# Evolution of Protein Regulation in the Vertebrate Glucose Sensor

**DOI:** 10.64898/2026.05.05.723016

**Authors:** S. Shirin Kamalaldinezabadi, Joshua I. Santiago, Johanna E. Papa, Yu Wang, Patrick A. Frantom, Hong Li, Robert Silvers, A. Carl Whittington, Brian G. Miller

## Abstract

Protein regulation is essential for cellular function and mis-regulation commonly causes disease. Despite this fact, we know little about how new regulatory strategies first emerge and how they evolve to act in concert to control complex physiological processes. Glucokinase (GCK), the body’s glucose sensor, lies at the heart of vertebrate glucose homeostasis and its activity is tightly controlled by multiple regulatory mechanisms. In the pancreas and liver, GCK is regulated by a unique form of monomeric allostery originating from the unliganded enzyme’s conformational dynamics. In the liver, GCK and GKRP form an inhibitory protein-protein interaction that sequesters GCK within the hepatocyte nucleus. Using a vertical, evolutionary approach, we resurrected extinct GCKs and GKRPs along correlated evolutionary trajectories. Using enzyme kinetics, limited proteolysis, hydrogen-deuterium exchange, high resolution NMR, and X-ray crystallography we determined the historical and molecular origins of protein regulation. Prior to the emergence of jawed vertebrates, a non-regulated GCK ancestor underwent a conformational expansion leading to monomeric allostery. This novel conformation includes an intrinsically disordered substrate binding loop. Paradoxically, the emergence of disorder did not require sequence change in the loop. The new GCK conformation also exposed a hydrophobic cleft. In the jawed vertebrate GKRP ancestor, a *de novo* loop insertion enabled exaptation of the pre-existing hydrophobic patch in GCK. Our results demonstrate how multiple, distinct regulatory strategies can arise at a central homeostatic control point through evolutionary addition of novel conformations. Additionally, our results provide a general mechanism for the emergence of heteromeric protein-protein interactions.

**Significance Statement:** Glucose homeostasis was a key innovation in vertebrate evolution. Here, we uncover the evolutionary basis of regulation in two key homeostatic proteins, glucokinase (GCK) and glucokinase regulatory protein (GKRP). We find that the unique cooperativity of vertebrate GCK resulted from an expansion of this enzyme’s conformational landscape. This expansion included sampling a new state and the emergence of intrinsic disorder, which did not require substitutions in the disordered region itself. We also discover that the GCK-GKRP interaction emerged when a pre-existing hydrophobic surface — a structural spandrel resulting from prior conformational expansion — was co-opted by loop insertion in GKRP, facilitating a new, inhibitory heteromeric interaction. Our results demonstrate how multiple, mechanistically distinct regulatory strategies arise from an ability to sample new protein conformations.

## Introduction

In vertebrates, the metabolic enzyme glucokinase (GCK) plays a central role in sensing and responding to whole-body glucose levels (1). As such, its activity is tightly controlled by multiple regulatory mechanisms (2). GCK catalyzes the ATP-dependent phosphorylation of glucose in the first and rate-limiting step of glycolysis in pancreatic β-cells (3, 4). Human GCK displays a sigmoidal kinetic response to glucose that is most sensitive to concentration changes near physiological glucose levels (Fig. 1A, red) (5). GCK activity dictates glucose-stimulated insulin release from the pancreas, thereby promoting systemic glucose uptake (3). In the liver, GCK activity triggers glycogen synthesis and is regulated by an inhibitory heteromeric protein-protein interaction with the glucokinase regulatory protein (GKRP) (6, 7). Interaction with GKRP also serves to sequester GCK within the hepatocyte nucleus, away from cytosolic glucose pools (8). The diverse regulatory mechanisms present in the GCK-GKRP system combine to provide exquisite control over blood glucose levels.

**Figure 1.**
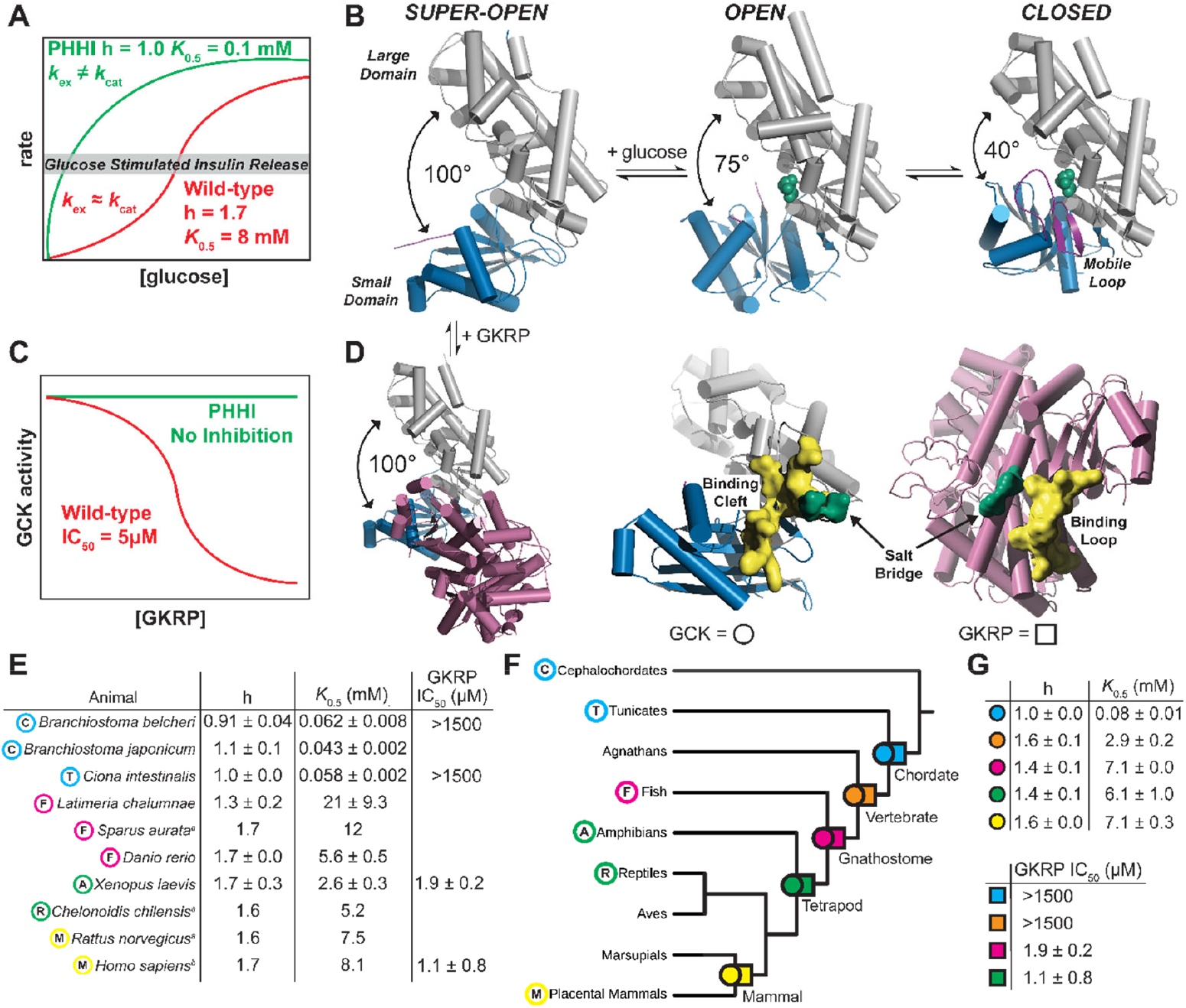
Structural and functional characteristics of extant and ancestral GCKs and GKRPs. (**A**) Human GCK displays a sigmoidal kinetic response to glucose characterized by a Hill coefficient of 1.7 and a *K*_0.5_ value of 8 mM (red), which regulates glucose stimulated insulin release (3). PHHI-associated variants eliminate cooperativity by disrupting the kinetic equivalency between the rate constant of conformational exchange, *k*_ex_, and *k*_cat_ (green) (10, 32). (**B**) The unliganded super-open GCK conformation (1V4T) and two glucose-bound states (glucose shown in green), open (4DCH) and closed (1V4S), differ by the magnitude of the interdomain opening angle. The intrinsically disordered mobile loop (purple) is only visible in the closed state. (**C**) GKRP is a competitive inhibitor of glucose binding to GCK (red) (7). Some PHHI variants are insensitive to GKRP inhibition (green). (**D**) GKRP binds exclusively to the super-open conformation in a 2060 Å^2^ cleft that includes a salt bridge between GCK residues E245/D247 and GKRP residues R297/R301 (green). (**E**) Functional characterization of extant GCK-GKRP pairs indicates that allosteric and GKRP-mediated regulation of GCK are vertebrate-specific traits. (**F**) Phylogenetic tree of GCK and GKRP evolution showing the positions of extant and reconstructed extinct proteins. (**G**) Characterization of reconstructed proteins identified functional transitions associated with allosteric regulation between chordate (blue) and vertebrate (orange) ancestors. GKRP-mediated regulation arose between the vertebrate (orange) and gnathostome (pink) ancestors. ^a^ – data from ref. (1); ^b^ – data from ref. (10); no superscript – data from this study.

GCK is the prototype for understanding monomeric kinetic cooperativity. It is autoregulated by its own conformational dynamics, which could be considered the “simplest” form of allosteric regulation (9, 10). Cooperativity arises from the temporal matching of the catalytic rate constant (*k*_cat_) with the rate constant for exchange between multiple unliganded GCK conformations (*k*_ex_) (9–12). Disruption of GCK autoregulation via point mutations leads to activation and loss of cooperativity, which causes persistent hyperinsulinemic hypoglycemia (PHHI) (Fig 1A, green) (13). Two mechanistically distinct activation mechanisms have been identified in PHHI variants. In α-type activation, cooperativity is abolished by substitutions that alter *k*_ex_, reduce conformational heterogeneity, and increase glucose binding affinity (10). In β-type activation, the rate of product release is elevated, reflected by a higher *k*_cat_ value, but conformational heterogeneity and glucose binding affinity remain unchanged (14). These two mechanistically distinct activation scenarios emphasize how precise matching of *k*_ex_ and *k*_cat_ is critical for normal glucose homeostasis.

GCK’s ability to sample multiple enzyme conformations is a key feature of its autoregulation (15). Crystal structures (Fig. 1B) show that unliganded GCK samples a super-open conformation characterized by a large interdomain opening angle that decreases upon glucose binding, and an intrinsically disordered, 30-residue mobile loop that folds into a β-hairpin in the closed, catalytically competent state (Fig. 1B; purple). The mobile loop is proteolytically susceptible in the unliganded state, providing a convenient proxy of GCK conformational heterogeneity and cooperativity (10). In addition to being required for kinetic cooperativity, the super-open conformation of GCK is necessary for regulation by GKRP. GKRP inhibits GCK (Fig. 1C) by binding exclusively to the super-open conformation of GCK (6) (Fig. 1D), and removal of the super-open conformation via PHHI-associated α-type activation disrupts GKRP inhibition (10).

To investigate the genetic, biochemical and biophysical origins of glucose homeostatic regulation, a hallmark of vertebrate physiology (16), we employed a vertical evolutionary investigation (17, 18) of GCK and GKRP. Our results reveal an evolutionary process that began before the appearance of jawed vertebrates, when a non-regulated GCK ancestor experienced conformational expansion, enabling a unique form of monomeric allostery. Expansion of GCK’s conformational landscape further included the emergence of an intrinsically disordered substrate binding loop, which did not require sequence changes within the loop. Conformational expansion in GCK exposed a cleft — a structural spandrel — that was subsequently co-opted by a GKRP loop insertion event to facilitate a new, inhibitory heteromeric interaction with GCK. This study provides unique insight into how genetic-based changes in protein sequence can lead to alterations in physical properties, including new conformational states, which facilitate new regulatory strategies including allostery and protein-protein interactions.

## Results and Discussion

### GCK Autoregulation and Inhibition by GKRP are Vertebrate Specific Traits

The cephalochordate (sister taxa to chordates) amphioxus contains a single hexokinase gene that is orthologous to vertebrate GCK, suggesting that vertebrate GCKs likely arose from a single locus prior to the diversification of chordates (19, 20). To determine the evolutionary timing of the emergence of regulatory mechanisms in the GCK-GKRP system, we sampled a series of extant species from three chordate subphyla and measured their GCK and GKRP functions (Fig. 1E; Fig. S1-S11). The tunicate *Ciona intestinalis* and two species from cephalochordate genus *Branchiostoma* contain non-cooperative GCKs (Hill coefficient (h) ∼ 1) with high sensitivity for glucose, suggesting these GCKs likely have functional and structural features similar to activated GCK variants. In contrast, all jawed vertebrates we sampled contain a cooperative GCK (h > 1) with moderate glucose sensitivity. The ability of GKRP to inhibit GCK shows a similar distribution among extant chordates. Tunicate and cephalochordate protein pairs failed to form an inhibitory interaction, whereas a functional GCK-GKRP interaction was observed in the tetrapod *Xenopus laevis* (Fig. S12-S14). Mapping of these parameters onto a chordate evolutionary tree supports a parsimonious model wherein both autoregulation and inhibition likely arose during the early diversification of vertebrates. Interestingly, the emergence of multiple regulatory mechanisms in the GCK-GKRP system correlates with the diversification of tissue types during early vertebrate evolution, including the appearance of the pancreas and liver (21–24).

The amino acid sequences of the cooperative GCKs among the jawed vertebrates are relatively highly conserved, with human GCK and the GCK from the teleost *Sparus aurata* having ≈ 80% identical residues. Comparison of the primary structures of human GCK and the non-cooperative GCK from *Branchiostoma belcheri* reveals a high degree of divergence with only ≈ 50% sequence identity. The situation is similar for comparing protein sequences of human GKRP and GKRP from *B. belcheri* (only ≈ 32% identity). This level of divergence precludes a traditional comparative, or horizontal, approach to determining the molecular origins of cooperativity. Thus, we employed a vertical approach to investigate the evolutionary and biophysical basis of regulation in the GCK-GKRP system by reconstructing extinct GCKs and GKRPs along parallel evolutionary trees (Fig. 1F, Fig. S15-S18) to identify functional and structural transitions associated with regulation *in situ*. We used PhyloBot (25) to reconstruct our ancestral sequences. We collected 194 extant GCK and hexokinase sequences across Chordata to generate our GCK phylogeny using the sequence from the echinoderm *Apostichopus japonicus* as an outgroup to the chordate sequences. Syntenic (gene-order conservation) analysis by Li, *et al*. (20) demonstrated that the vertebrate GCK locus and the hexokinase locus in the cephalochordate *Branchiostoma japonicum* are orthologous, enabling us to generate an orthologous group of sequences sequences (all other vertebrate hexokinases are paralogs to the vertebrate GCKs). Our best-fit phylogeny of GCK evolution suffers from long branches in the transition period during early vertebrate evolution (Fig. S15 & Fig. S16). Additionally, there is discordance in the GCK tree that leads to a mismatch with the species tree in multiple instances (SI Materials and Methods). Although ancestral sequence reconstruction has been repeatedly shown to be quite robust to a variety of sources of phylogenetic uncertainty including branch lengths and topology (26–28), long branch lengths and discordance may lead to uncertainty in our phylogenetic reconstruction that may affect our ancestral sequences (Fig. S18-23, Table S1 & S2). In addition, our strategy for reconstructing alternate ancestors could impact the degree to which biochemical properties are robust to phylogenetic uncertainty. We demonstrate below that our reconstructed ancestral phenotypes are robust to at least some sources of uncertainty. Our GKRP tree was generated from 98 extant eukaryote sequences spanning from Excavata to Mammals (Fig. S17). Only a single locus exists for GKRP across the extant taxa sampled ensuring orthology in our sequence set. Both trees generally conformed to known taxonomic relationships and had sampling that allowed reconstruction of approximately contemporaneous pairs of extinct GCKs and GKRPs.

### Evolution of Autoregulation in Glucokinase

Our previous work on regulated GCKs and activated variants demonstrates a very strong correlation of biochemical and biophysical parameters with the presence and absence of cooperativity and GKRP inhibition (6, 9, 10, 12, 14, 29–34). We postulated that this correlation will generally hold across the evolutionary histories of GCK and GKRP. Autoregulation in human GCK manifests as a cooperative kinetic response to glucose, with a midpoint value (*K*_0.5_) in the millimolar range. We resurrected extinct GCKs along the chordate evolutionary tree and found a clear transition in cooperativity (Fig. 1G) between the ancestral chordate GCK (cGCK) and ancestral vertebrate GCK (vGCK) that correlates with measurements of extant GCKs (Fig. S24-S27). The non-cooperative kinetics and high glucose sensitivity seen in the extant, non-vertebrate chordates is reflected in cGCK’s Hill coefficient (1.0) and *K*_0.5_ value (77 µM). Cooperativity in GCK arose by the time of the last common ancestor of vertebrates, represented by vGCK, which displays a Hill coefficient (1.6) and *K*_0.5_ value (2.9 mM) similar to extant vertebrate GCKs. The sigmoidal glucose response curve and characteristic moderate glucose sensitivity seen in human GCK was established early during vertebrate evolution and has remained highly conserved during the evolution of gnathostomes (gGCK), tetrapods (tGCK), and mammals (mGCK) (Figs. S28-S33). To test the robustness of our reconstructed ancestors, we also resurrected and characterized the “worst” versions of cGCK and vGCK from our PhyloBot reconstruction, as well as the AltAll (27) versions. These alternative versions of cGCK and vGCK were functionally identical to the most likely versions described above (Fig. S34-S4136), demonstrating that our reconstruction is robust to variability in alignment algorithm and model of sequence evolution (see SI Methods for more detail on robustness of the reconstruction).

One main feature distinguishing cooperative and non-cooperative GCKs is a difference in the structure and dynamics of the mobile loop in the unliganded state (10, 12, 32, 35). Prior work demonstrates that autoregulation in GCK is strongly correlated with proteolytic susceptibility of the loop (10), which requires local unfolding under limiting proteolysis conditions for cleavage by a generic protease (36–38). We measured the proteolytic susceptibility of extant GCKs (Fig. S42-45) and each resurrected GCK (Fig. 2A) and found that the transition in proteolytic susceptibility matched the transition in glucose kinetics. The non-cooperative cGCK undergoes no substantial proteolysis on the timescale of the measurement indicating minimal local unfolding of its mobile loop (Fig. S46). Conversely, vGCK displays proteolytic susceptibility similar to human GCK (τ_1/2_ = 1.54 ± 0.22 min; Fig. S47) (10).These data suggest the presence of a less ordered, more flexible loop in vGCK, whereas cGCK possesses a more structured mobile loop. The emergence of cooperativity appears to depend on an intrinsically disordered mobile loop. Once proteolytic susceptibility of the mobile loop in vGCK was established, it remained conserved across vertebrate evolution (Fig. 2A; Figs. S48-S50). To test whether sequence changes solely in the mobile loop were responsible for cooperativity, we installed the loop sequence from cooperative vGCK into non-cooperative cGCK. This chimeric ancestor lacked cooperativity (Fig. S51) and displayed proteolytic resistance similar to cGCK (Fig. S52). Thus, intrinsic disorder in the mobile loop is sequence-independent and must depend on contacts with other structural elements in the protein scaffold.

**Figure 2.**
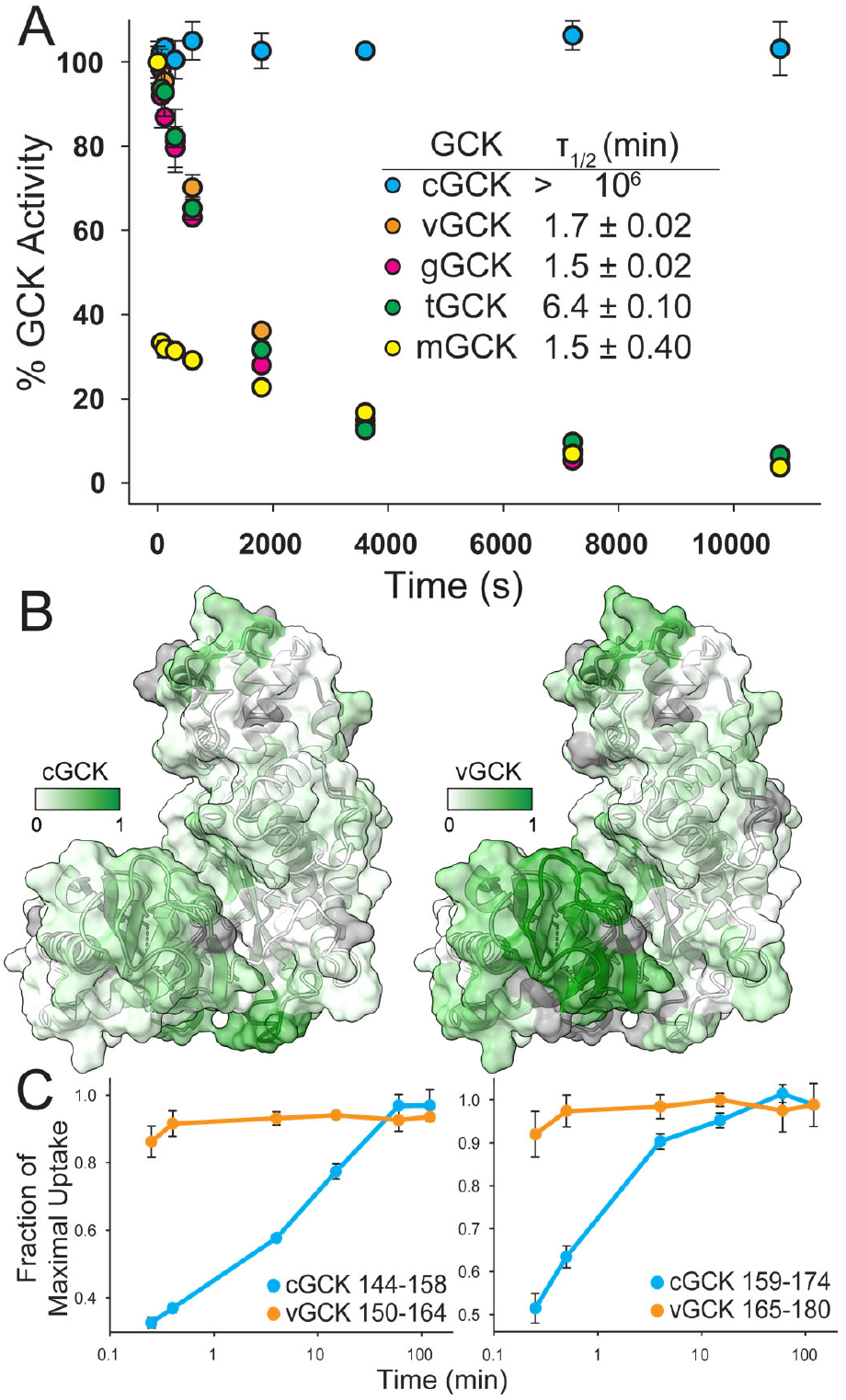
Evolutionary and biochemical origins of GCK autoregulation. (**A**) Proteolytic susceptibility of ancestral GCKs, indicative of mobile loop intrinsic disorder, correlates with the emergence of cooperativity. Once established, proteolytic susceptibility of the mobile loop was conserved across vertebrate evolution. (**B**) HDX-MS comparative analysis of cGCK and vGCK indicates altered regions of conformational heterogeneity and/or backbone dynamics coincidental with the emergence of cooperativity, most evident in the small domain. Fraction of maximal uptake is mapped onto the closed structure of human GCK (PDB: 1V4S) and indicated by the intensity of green shading. (**C**) The mobile loop spans residues 151 to 180 (human GCK numbering). Our HDX-MS data covers the majority of the mobile loop in both cGCK and vGCK and demonstrates that dynamics are elevated in the vGCK mobile loop, correlating with the emergence of proteolytic susceptibility and cooperativity.

Based on the established importance of conformational heterogeneity to cooperativity in human GCK, we hypothesized that the functional transition from the non-cooperative chordate ancestor to the cooperative vertebrate ancestor depended on an expansion of conformational space and/or dynamics. To test this postulate, we performed amide hydrogen/deuterium exchange mass spectrometry (HDX-MS) analysis on cGCK and vGCK. Based on common overlapping peptide fragments, the cooperative vGCK ancestor displays elevated HDX compared to the non-cooperative cGCK ancestor, consistent with an increase in conformational heterogeneity and/or dynamics (Fig. S53 & S54). In cGCK the highest levels of HDX are observed in three regions of the polypeptide distal to the active site and mobile loop (Fig. 2B, left). In vGCK, exchange is generally elevated and more widely distributed, also involving the small domain (Fig. 2B, right) and mobile loop (Fig. 2C) where conformational changes are known to be linked to cooperativity. Interestingly, the regions of highest exchange in non-cooperative cGCK are somewhat suppressed in cooperative vGCK. This observation is consistent with the possibility that some level of intrinsic dynamics is required for catalysis, but that these dynamics can be re-located within the GCK scaffold to facilitate regulation, while maintaining function.

Glucose binding to human GCK induces large chemical shift perturbations (CSPs) in ^13^C^δ1^-labeled isoleucine moieties observed in 2D ^1^H-^13^C HMQC NMR spectra, most notably in the small domain, the mobile loop, and residues directly involved in glucose binding (9, 10, 12). In the unliganded state, most of the small domain residues are invisible, indicating they are exchange-broadened and experience μs-ms timescale dynamics. Notably, two residues located in the mobile loop are visible as prominent, high intensity cross-peaks, indicating they experience rapid dynamics and the mobile loop is flexible. In contrast, all of the large domain residues are visible, indicating slow dynamics on the NMR timescale. Glucose binding to human GCK induces ordering as evidenced by an increase in chemical shift dispersion. This is also reflected in the appearance of cross-peaks for previously invisible small domain residues and a shift of the mobile loop residues to the ordered region of the spectrum associated with folding of the mobile loop into a β-hairpin. The μs-ms timescale dynamics present in the unliganded state of human GCK are essential for cooperativity, and the conversion in dynamics upon glucose binding that is observed through changes in NMR spectra is a hallmark behavior of cooperative GCK variants (9, 10, 12). In non-cooperative, α-type activated human GCK variants, the differences between the unliganded and glucose-bound NMR spectra are minimal as the μs-ms timescale dynamics in the unliganded state are suppressed, suggesting the α-type unliganded conformational distribution favors a binding competent state that resembles the glucose-bound state (12).

Changes in structure and dynamics in cGCK and vGCK were tracked using 2D ^1^H-^13^C ALSOFAST-HMQC spectra of ^13^C^δ1^-labeled isoleucine samples (Fig. 3A and 3B, respectively, Fig. S55-58). Despite differences in the number and position of isoleucine residues among cGCK, vGCK, and human GCK, the isoleucine residues in both ancestral GCKs display a similar distribution to human GCK with many identical positions across all three allowing a robust comparison (Fig. 3). Isoleucines residing within the large domain display visible peaks with little to no CSPs upon glucose binding in both cGCK and vGCK (Fig. 3C and 3D, respectively). This indicates that the chemical environment within the large domain in both unliganded and liganded states is similar despite the amino acid differences between cGCK and vGCK. Differences in NMR spectra of the two unliganded ancestral proteins are consistent with a shift in dynamics and/or conformational heterogeneity that occurred during the evolution of cooperativity. Most peaks in cooperative vGCK are clustered near ∼ 0.5-1.0 ppm and ∼12-16 ppm for ^1^H and ^13^C, respectively, which is very similar to human GCK (12). Positions 126, 253, 293, and 452 in the small domain display weak, broadened peaks indicating conformational exchange on the µs-ms timescale. Positions 163 and 165 located in the mobile loop display sharp peaks indicating rapid dynamics. The peaks observed in unliganded cGCK are more dispersed and have a smaller line width, especially in the congested area where peaks are resolved in cGCK. This signifies a loss or significant reduction in conformational exchange on the μs-ms timescale, correlating with the lack of cooperativity in cGCK. In vGCK, probes located at the core of the small domain (residues 110, 126, and 189) show significant CSPs, indicating a more substantial change in chemical environment and dynamics within the small domain upon glucose binding. Additionally, very large CSPs in the α13 helix region (residues 211 and 452) are absent in cGCK. In cGCK, moderate CSPs are only observed for 110 and 199 located in the small domain and 165 located in the mobile loop, which are all in close proximity to bound glucose. Isoleucines in the core of the small domain (residues 107, 126, and 189) and the α13 helix and its surrounding contacts (residues 91 and 452) display no substantial CSPs. Our NMR results demonstrate a scenario where a non-cooperative ancestral GCK that resembled an α-type activated modern GCK shifted to a cooperative GCK through the emergence of increased conformational heterogeneity and dynamics. Once established along the evolutionary trajectory, this structural and dynamic phenotype was likely completely conserved across vertebrate evolution.

**Figure 3.**
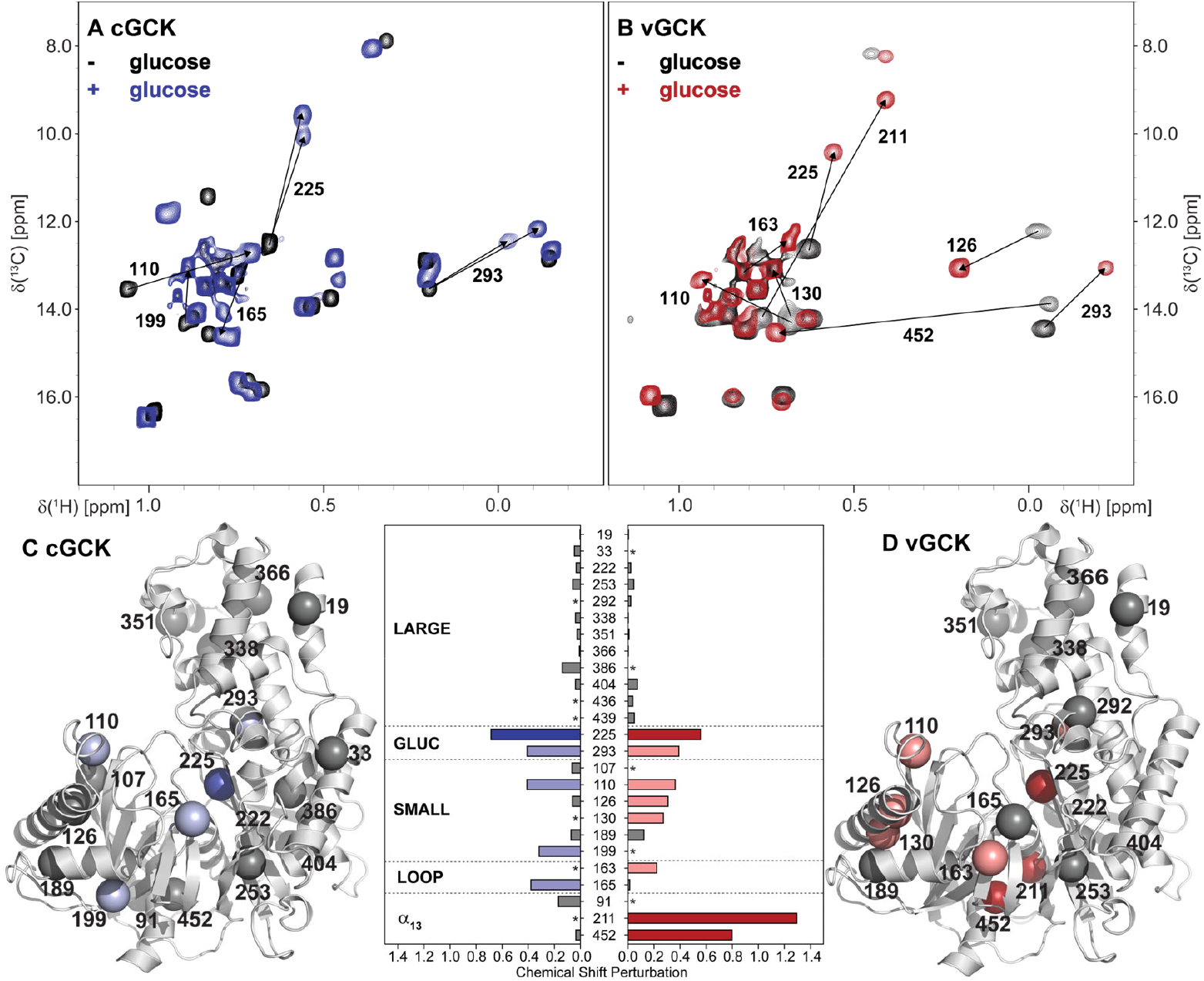
NMR-spectroscopic hallmarks of cGCK and vGCK. (**A**) ^1^H-^13^C ALSOFAST-HMQC spectra of unliganded (black) and liganded (blue) non-cooperative cGCK. (**B**) ^1^H-^13^C ALSOFAST-HMQC spectra of unliganded (black) and liganded (red) cooperative vGCK. Large changes in chemical shift upon binding are denoted with arrows. Structural distribution of chemical shift perturbations observed upon binding of glucose for cGCK (**C**) and vGCK (**D**). Insignificant CSPs below 0.2 ppm are shown in gray, moderate CSPs between 0.2 and 0.5 ppm are shown in light blue/light red, and strong CSPs above 0.5 ppm are shown in dark blue/dark red. The location of isoleucine probes (spheres) is mapped onto the closed structure of human GCK (1V4S). Color coding of the isoleucine probes matches the classification of CSPs. Numbering is based on human GCK (GenBank: AAA52562.1; see SI alignment files for corresponding numbering in cGCK and vGCK).

Overall, cGCK and vGCK share 72.4% sequence identity; 128 amino acids differentiate the non-cooperative ancestor from its cooperative descendent. This large number of positional variations confounds efforts to identify the residue(s) responsible for the emergence of monomeric kinetic cooperativity. Notably, however, cGCK contains six residues resembling or matching non-cooperative, PHHI-associated human GCK variants. To test whether these substitutions are sufficient to install cooperativity into cGCK, we created and characterized the L65T-I91V-R99W-D180N-M211I-A214Y variant (human GCK numbering). These substitutions resulted in a non-cooperative enzyme with a reduced *k*_cat_ value (6.9 s^-1^) and a glucose *K*_m_ value of 53 mM (Fig. S59). We also characterized a hybrid cGCK in which small domain residues matched those of vGCK, since past NMR studies of human GCK demonstrated this region is the major site of conformational exchange in the unliganded state. This small-domain cGCK variant was also non-cooperative, displaying a glucose *K*_m_ value of 114 mM (Fig. S60). Along with the results of the mobile loop replacement variant, these findings suggest that cooperativity cannot be engineered simply via targeted alterations to a specific polypeptide region, consistent with a complex, historically contingent evolutionary trajectory. In this type of scenario, amino acid substitutions that alter function may only be operational in the presence of prior substitutions that potentially have no effect on function, leading to an unpredictable evolutionary trajectory (39–45)..

### Evolution of Inhibition by GKRP

The GCK-GKRP interaction depends on the presence of the super-open conformation in unliganded GCK (6, 46). The hydrophobic GKRP binding cleft (Fig. 3A) forms in unliganded GCK as the opening angle increases to ≈ 100°, moving two negatively charged residues (Glu245 and Asp247) into position to form coulombic interactions with two positively charged GKRP residues (Arg301 and Arg297). The opening angle closes to ≈ 75° upon glucose binding, which disrupts the GKRP binding cleft (Fig. 4A). To investigate whether unliganded cGCK samples the super-open conformation, and thus might be susceptible to GKRP inhibition, we pursued the crystal structure of this enzyme. We were unable to crystallize the cGCK sequence from our PhyloBot reconstruction. Instead, we were able to obtain crystals for the cGCK sequence from our FastML reconstruction (Fig. 4A; Table S3). These two proteins were functionally identical (Phylobot cGCK *k*_cat_ = 47 ± 8s^-1^, *K*_m_ = 77 ± 7 µM, h = 1.0 ± 0.0; FastML cGCK *k*_cat_ = 58 ± 19s^-1^, *K*_m_ = 73 ± 6 µM, h = 0.92 ±

**Figure 4.**
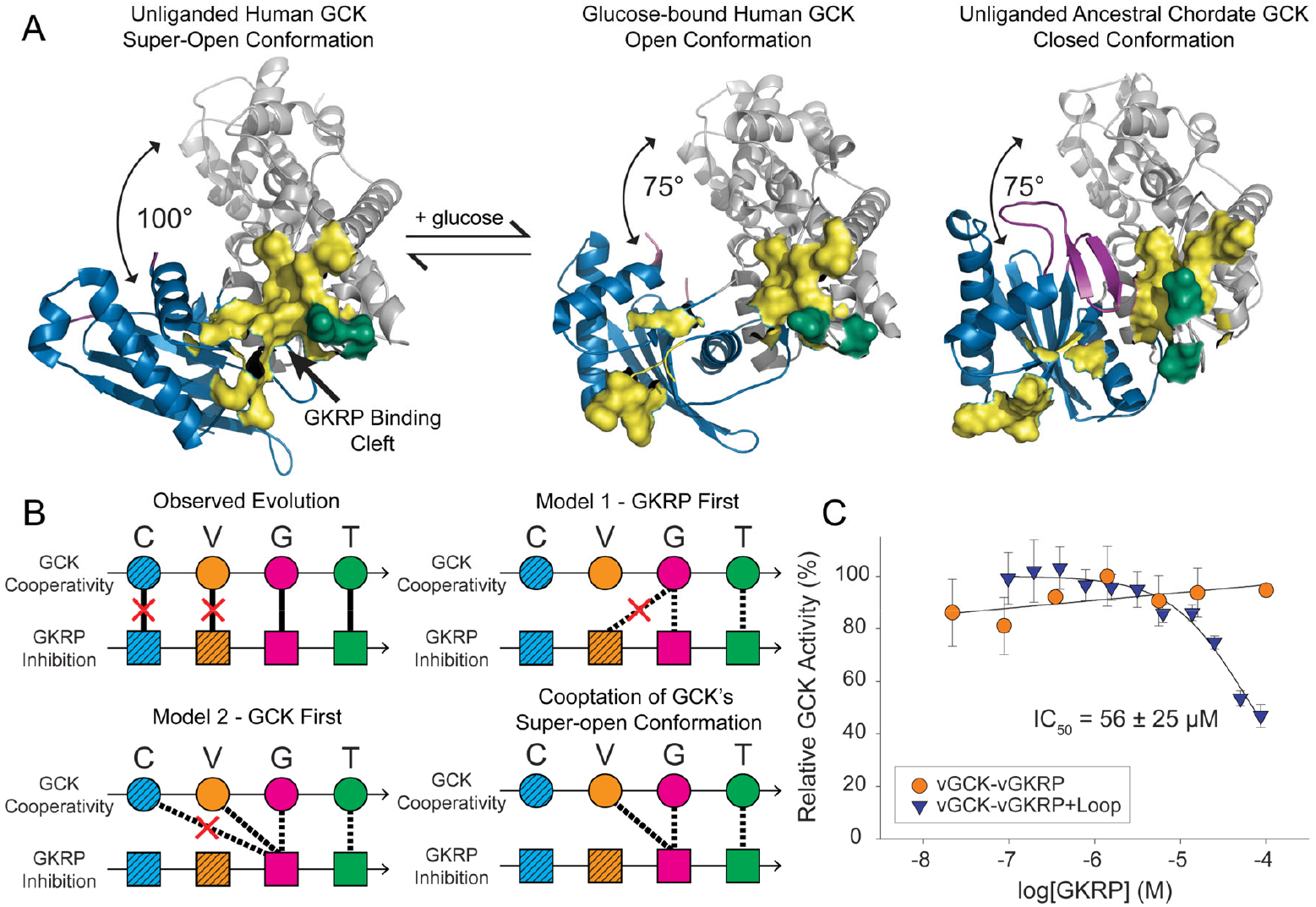
Evolutionary origins of GKRP-mediated GCK regulation. (**A**) The GKRP binding cleft in human GCK is formed along with the large opening angle in the unliganded state (PDB: 1V4T). Glucose binding closes the opening angle and disrupts the GKRP binding cleft, shown in the “open” conformation (PDB: 4DCH). The crystal structure of unliganded cGCK shows a more closed opening angle of 75°, an ordered mobile loop, and no GKRP binding cleft. Coloring is identical to Fig. 1. Bound glucose is not shown here. (**B**) We observed an evolutionary trajectory in which reconstructed chordate (blue) and vertebrate (orange) ancestral GCK-GKRP pairs failed to interact (red X), but gnathostome (pink) and tetrapod (green) ancestors did interact. We tested two models for the evolutionary timing of the GCK-GKRP interaction. In the GKRP first model, which was not supported (red X), vGKRP is expected to inhibit all GCK ancestors. In the GCK first model, susceptibility to GKRP inhibition arose before the ability of GKRP to inhibit. In support of this model, gGKRP inhibits vGCK which can adopt the super-open state, but gGKRP cannot inhibit cGCK. Our data support a model involving co-optation of GCK’s super-open conformation by GKRP following loop insertion. (**C**) Inserting a six-residue loop into vGKRP (orange) was sufficient to introduce inhibitory activity into vGKRP (blue, + loop).

0.4; Figs. S61 & S62). The structure of unliganded cGCK reveals a structure similar to the glucose-bound, “open” structure of human GCK (Fig 1B), with an interdomain opening angle of ≈ 75° and an ordered mobile loop. These data are consistent with our observed proteolytic resistance, HDX-MS exchange data, and NMR measurements of this ancestor and offers a structural explanation for the lack of autoregulation in this enzyme, as conformational heterogeneity in the unliganded enzyme via sampling of the super-open conformation is required for kinetic cooperativity (Fig. 1). The crystal structure of cGCK provides the first example of an unliganded vertebrate glucokinase with the mobile loop in a folded conformation.

Consistent with the inability of cGCK to sample the super-open conformation, we found it was unable to be inhibited by its contemporaneous regulatory protein, cGKRP (Fig. S63). Our results suggest that the super-open conformation, and monomeric cooperativity, had evolved by the time vGCK emerged in the last common ancestor of vertebrates. Intriguingly, vGCK, with a Hill coefficient of 1.6, is not inhibited by its contemporaneous regulatory protein, vGKRP, despite sampling the super-open conformation (Fig. 1; Fig. S64). The next two pairs of ancestral GCK-GKRPs corresponding to the gnathostome ancestor (gGCK-gGKRP; Fig. S65) and tetrapod ancestor (tGCK-tGKRP; Fig. S66) show Hill coefficients and inhibitory constants resembling their extant descendants (Fig. 1), suggesting that the GCK-GKRP inhibitory interaction was present since at least the last common ancestor of jawed vertebrates.

To determine when and how a functional GCK-GKRP interaction emerged, we tested two competing models describing the evolutionary timing of the GCK-GKRP interaction (Fig. 3B; Figs. S67-69). In the GKRP first model, vGKRP is expected to be capable of inhibiting GCKs from descendent ancestors, e.g. gGCK, but the interaction required further mutation in GCK to reach physiologically relevant inhibition. In the GCK first model, the susceptibility of GCK to inhibition evolved before the ability of GKRP to inhibit — *i*.*e*. the super-open conformation was able to be inhibited before GKRP was able to bind to it. Indeed, gGKRP inhibits its ancestor vGCK (Fig. S68). Our model demonstrates that the super-open conformation already existed in GCK before the GKRP attained the capacity to interact. The GKRP binding cleft in GCK represents a biophysical spandrel, as it emerged as a physical side product of the super-open conformation rather than as an adaptive structure itself (47–49).

Exaptation (50) of the GCK spandrel formed by the super-open conformation must have required further GKRP mutation to enable inhibition, as the binding cleft is highly conserved between vGCK and gGCK. We compared the sequences of vGKRP and gGKRP with those of their modern descendants and found that a six-residue loop insertion in the gGKRP ancestor can account for the emergence of the GCK-GKRP interaction. Among non-vertebrate chordates, this region of the alignment is highly variable, suggesting there may have been rapid sequence change prior to the evolution of the GCK-GKRP interaction. We used the method described by Kaltenbach *et al* (51) for informed reconstruction of indels. Although the loop sequence varies, the presence of the loop is completely conserved among extant vertebrate GKRPs and can be seen in the GCK-GKRP crystal structure (Fig. 1; S70). We added the six-residue loop from gGKRP into the vGKRP background and assayed vGKRP with its contemporaneous GCK, vGCK. Insertion of the loop, which could be accomplished in a single genetic event, is sufficient to introduce inhibitory activity into GKRP (Fig. 4C; Fig. S71), although it is weaker inhibition than seen in extant interacting pairs. Due to the low sequence sampling density in this region of the GKRP tree, the exact timing of the loop-insertion event is difficult to pinpoint. However,.our results suggest that exaptation enabled by *de novo* evolution was required for the emergence of the GCK-GKRP heteromeric interaction.

### Disorder, Duplication, and Exaptation Enable Regulatory Complexity

Regulation via intrinsic disorder in proteins is often associated with cell signaling, where one protein can interact with multiple binding partners due to the shape-shifting capabilities of intrinsic disorder (52). Here, we demonstrate that the emergence of an intrinsically disordered binding loop was critical for the evolution of monomeric cooperativity in vertebrate GCK. Autoregulation in GCK specifically, and intrinsically disordered regions generally, could be considered the simplest form of protein regulation. In the case of vertebrate GCK, conformational heterogeneity and intrinsic disorder provided dynamics on a timescale matching the rate of turnover, leading to its characteristic sigmoidal kinetic response. Autoregulation in GCK provides exquisite substrate sensitivity and tunability (53). However, even mild disruption of conformational heterogeneity and mobile loop dynamics causes disease by disrupting the timescale matching required for cooperativity (10). Elevated dynamics in the mobile loop, beyond the level of wild-type, increases the rate of product release, causing β-type activation (14). In contrast, decreased disorder in the mobile loop is associated with α-type activation, reverting the enzyme back to its ancestral state, a phenotypic atavism.

The exaptation of a pre-existing hydrophobic surface during the formation of the GCK-GKRP interaction could be a general mechanism underlying the evolution of heteromeric protein-protein interactions. Previously, Whittington *et al*. (54) demonstrated *in silico* that exaptation of pre-existing hydrophobic surfaces in phospholipase A2 paralogs was responsible for the evolution of a heteromeric neurotoxin in rattlesnakes. Subsequently, Pillai *et al*. (55) showed experimentally that a similar mechanism underlies the evolution of hemoglobin. In both examples, the interactions were between closely related and structurally similar paralogs, suggesting that the mechanism of exaptation of pre-existing surfaces may be a result of symmetric expansion akin to homomeric interactions (56). A recent follow up study supports this hypothesis for hemoglobin (57). Our results demonstrate that even in protein-protein interactions between non-related proteins where no symmetry exists, exaptation of a pre-existing surface is critical. A similar mechanism facilitated by horizontal gene transfer underlies the formation of a heteromeric interaction in cyanobacteria (58). In a pair of yeast DNA-binding proteins a heteromeric protein-protein interaction likely evolved from previously non-interacting interfaces via co-localization at adjacent DNA-binding sites and co-optation of a surface of one of the pair relatively enriched in hydrophobic residues (59). Similar mechanisms underlying the formation of multimeric protein-protein interactions across diverse protein systems suggest that exaptation of pre-existing surfaces is a general and genetically simple mechanism for the emergence of heteromeric interactions.

Subfunctionalization after gene duplication may lead to a period of neutral evolution facilitating subsequent neofunctionalization in the daughter gene copies (60, 61), providing a tidy model consistent with our results. The extant GCKs assayed in this study are members of the larger vertebrate hexokinase family, which contains four paralogous enzymes that phosphorylate hexose sugars and are differentially expressed in various tissues (62, 63). GCK (HK IV) consists of a single kinase domain, while the other three hexokinase genes consist of two kinase domains, suggesting a series of gene duplication events and a fusion event during hexokinase evolution (1). Four HK genes were present before the vertebrate diversification, as they are present in all classes of jawed vertebrate (19, 64). The vertebrate hexokinases are generally regulated by a diverse set of mechanisms in addition to those identified in the present study (62). This increase in regulatory complexity correlates with the diversification of tissue types in early vertebrate evolution (21–24) where the presence of multiple hexokinase paralogs likely allowed tissue-specific regulatory features. Specifically, the regulatory mechanisms of the GCK-GKRP system likely emerged along with pancreatic islet cells and hepatic cells to provide a central control hub mediating systemic glucose homeostasis in vertebrates. Our results show that within the GCK-GKRP system, novel regulatory mechanisms arose from conformational heterogeneity, disorder, and exaptation, which were all likely facilitated by gene duplication.

## Materials and Methods

### Phylogenetics

Sequences of extant GCKs and GKRPs were obtained from the NCBI non-redundant protein database and EMBL’s Ensembl in 2018. In total, 194 sequences from across Deuterostomia were used in the GCK reconstruction, with the echinoderm *Apostichopus japonicus* GCK set as the outgroup. 98 sequences from Amoebas and Holozoa were used in the GKRP reconstruction. PhyloBot was used to perform sequence alignments, phylogenetic reconstruction, and estimation of ancestral sequences (25). Separate reconstructions were performed for the GCK and GKRP sequences, and gaps were reconstructed according to Kaltenbach, *et al* (51).

### Recombinant protein production and purification

Ancestral GCK and GKRP coding sequences cloned into pET-22b (+) and codon-optimized for expression in *Escherichia coli* were ordered from GenScript (Piscataway, NJ). N-terminal hexahistidine-tagged recombinant proteins were produced and purified as previously described (29). When working with vGKRP, chromatography buffers were supplemented with L-arginine (50 mM), L-glutamate (50 mM), and DTT (10 mM) to mitigate aggregation and increase protein stability.

### Enzyme kinetics, inhibition assays, and limited proteolysis studies

GCK activity and GKRP inhibition were measured via an enzyme-coupled spectrophotometric assay in which the production of glucose-6-phosphate (G6P) is linked to the conversion of NADP^+^ to NADPH via glucose 6-phosphate dehydrogenase (G6PDH) from *Leuconostoc mesenteroides* (Sigma-Aldrich) as previously described. Proteolysis assays of unliganded GCKs were performed with thermolysin (Sigma-Aldrich) as previously described (10, 35).

### Crystallography and structure determination

Crystals of FastML-cGCK were obtained via the hanging drop method by mixing 1 µL of protein solution (30 mg/mL, purified as described above) with 1 µL of a well solution (0.1 M Tris-HCl pH 7.5, 25% w/v PEG 2000 MME, and 0.3 M sodium acetate) and equilibrating the drop against the same well solution at room temperature for 3-5 days. The structure of cGCK was determined using molecular replacement via the MRage package in Phenix (65) by serially searching for the solution using the individual domains of human GCK (PDB: 1V4S) as probes. Structure refinement was carried out with Phenix until satisfactory crystallographic residual factors and stereochemical parameters were reached (Table S1).

### Hydrogen-deuterium exchange mass spectrometry

Protein samples were concentrated to 50 µM and 5 µL of each sample was added to 45 µL potassium phosphate buffer (25 mM, pH 7.0 prepared in H_2_O or D_2_O) with incubation at 25°C. After a set time, the sample was quenched with 25 µL of an ice-cold solution containing 200 mM TCEP, 8M urea and 1.0% formic acid (pH ∼ 2.3). A 25 µL aliquot of ice-cold 1.0% formic acid was added to the sample, followed by addition of 400 µL of ice-cold 0.1% formic acid. 100 µL aliquots were injected onto a Waters HDX Manager UPLC system coupled to a Xevo G2-XS Q-Tof mass spectrometer. Protein samples were digested with an in-line pepsin column (Waters Enzymate BEH pepsin column, 2.1 × 30 mm, maintained at 15 °C), followed by desalting (2.1 × 5 mm ACQUITY UPLC BEH C-18 VanGuard trapping column) and then separation on 1.0 × 100 mm ACQUITY UPLC BEH C-18 analytical column maintained at 4 °C. Peptides were separated with a 5-minute gradient from 5%-85% acetonitrile in 0.1% formic acid and a flow rate of 30 µL/min. Peptide ion and MS^E^ fragment data from non-deuterated samples were analyzed with ProteinLynx Global Server software (Waters Corp., Milford, MA) to generate a peptide coverage map. Mass shifts in deuterium exchanged peptides were determined using DynamX 3.0 (Waters Corp., Milford, MA) to generate deuterium incorporation plots. The maximum relative exchange for each peptide was determined using a fully deuterated control sample. Data and error bars shown are the average of three replicate experiments (Fig. 2.; Fig. S47).

### Nuclear Magnetic Resonance

Methyl groups on isoleucine side chains of cGCK and vGCK were radio-labeled using the isoleucine precursor, alpha-ketobutyric acid (methyl-^13^C, 99%; 3,3-D_2_, 98% CDLM-7318-0; Cambridge Isotope Laboratories) as previously described (10, 12, 32). Enzymes were purified as described above and dialyzed against 25 mM potassium phosphate pH 8.0, 50 mM KCL and 10 mM DTT at 4°C. All spectra were acquired on a Bruker AVANCE III 700-MHz NMR spectrometer equipped with a TCI cryogenic probe. 2D ^1^H-^13^C ALSOFAST-HMQC NMR experiments (pulse program: afhmqcgpphsf) were used to obtain proton and carbon correlations. The data were recorded as matrices of 1536 × 160 complex datapoints. The spectral widths were 15.0012 ppm and 22.0000 ppm for F2 and F1 directions respectively. The recycling delay d1 was 0.3 seconds. Proton chemical shifts were referenced using DSS as an internal reference, while carbon-13 chemical shifts were reference indirectly. All spectra were apodised with a QSIN function with an SSB of 3.5 in each dimension before zero filling. The experiments were performed at 298 K. The spectra were collected using either 512 or 1024 scans depending on sample concentration. Site-specific assignments of Ile resonances were accomplished by single-site substitution of Ile to Val followed by expression, purification, and the collection of the 2D spectra to identify the missing cross peak, as previously described (10, 12, 32). NMR data processing and spectral analysis were performed using TopSpin 4.1.4 program. The chemical shift perturbation (CSP) upon glucose binding for each Ile residue were calculated using the equation CSP = {0.5 × [(Dd_1H_)^2^ + (0.25 × Dd_13C_)^2^]}^0.5^, where Dd_1H_ is the change in chemical shift in ^1^H and Dd_13C_ is the change in chemical shift in ^13^C.

Additional experimental methods are provided is Supporting Information.

## Data Sharing

All data, including the full sequences of extant and reconstructed proteins, are provided in Supporting Information. The crystal structure for ancestral GCK has been deposited in the Protein Data Bank (PDB entry 9XYT).

## Acknowledgments

Research reported in this publication was supported by the National Institute of General Medical Sciences of the National Institutes of Health under Award Numbers R01GM133843 (B.G.M.), R35GM142912 (R.S.) R01GM112919 (P.A.F.) and R35GM152081 (H.L). The content is solely the responsibility of the authors and does not necessarily represent the official views of the National Institutes of Health. Additional funding was provided by the FSU Council on Research and Creativity (B.G.M). The authors thank Dr. Thayumanasamy Somasundaram for assistance with X-ray data collection and analysis.

## References

1. M. L. Cárdenas, A. Cornish-Bowden, T. Ureta, Evolution and regulatory role of the hexokinases. Biochim. Biophys. Acta Mol. Cell Res. 1401, 242–264 (1998).

2. F. M. Matschinsky, D. F. Wilson, The Central Role of Glucokinase in Glucose Homeostasis: A Perspective 50 Years After Demonstrating the Presence of the Enzyme in Islets of Langerhans. Front. Physiol. 10, 148 (2019).

3. F. M. Matschinsky, Glucokinase as glucose sensor and metabolic signal generator in pancreatic beta-cells and hepatocytes. Diabetes 39, 647–652 (1990).

4. S. M. Sternisha, B. G. Miller, Molecular and cellular regulation of human glucokinase. Arch. Biochem. Biophys. 663, 199–213 (2019).

5. A. Cornish-Bowden, M. L. Cardenas, “Glucokinase: A Monomeric Enzyme with Positive Cooperativity” in Glucokinase and Glycemic Disease: From Basics to Novel Therapeutics, (KARGER, 2004), pp. 125–134.

6. T. Beck, B. G. Miller, Structural basis for regulation of human glucokinase by glucokinase regulatory protein. Biochemistry 52, 6232–6239 (2013).

7. E. Van Schaftingen, A. Vandercammen, M. Detheux, D. R. Davies, The regulatory protein of liver glucokinase. Adv. Enzyme Regul. 32, 133–148 (1992).

8. C. Shiota, J. Coffey, J. Grimsby, J. F. Grippo, M. A. Magnuson, Nuclear import of hepatic glucokinase depends upon glucokinase regulatory protein, whereas export is due to a nuclear export signal sequence in glucokinase. J. Biol. Chem. 274, 37125–37130 (1999).

9. M. Larion, R. K. Salinas, L. Bruschweiler-Li, R. Brüschweiler, B. G. Miller, Direct evidence of conformational heterogeneity in human pancreatic glucokinase from highresolution nuclear magnetic resonance. Biochemistry 49, 7969–7971 (2010).

10. A. C. Whittington, et al., Dual allosteric activation mechanisms in monomeric human glucokinase. Proceedings of the National Academy of Sciences 112, 11553–11558 (2015).

11. C. Frieden, Kinetic aspects of regulation of metabolic processes. The hysteretic enzyme concept. Journal of Biological Chemistry 245, 5788–5799 (1970).

12. M. Larion, R. K. Salinas, L. Bruschweiler-Li, B. G. Miller, R. Brüschweiler, Order-Disorder Transitions Govern Kinetic Cooperativity and Allostery of Monomeric Human Glucokinase. PLoS Biol. 10, e1001452 (2012).

13. H. B. T. Christesen, et al., Activating glucokinase (GCK) mutations as a cause of medically responsive congenital hyperinsulinism: Prevalence in children and characterisation of a novel GCK mutation. Eur. J. Endocrinol. 159, 27–34 (2008).

14. S. M. Sternisha, P. Liu, A. G. Marshall, B. G. Miller, Mechanistic Origins of Enzyme Activation in Human Glucokinase Variants Associated with Congenital Hyperinsulinism. Biochemistry 57, 1632–1639 (2018).

15. K. Kamata, M. Mitsuya, T. Nishimura, J. I. Eiki, Y. Nagata, Structural basis for allosteric regulation of the monomeric allosteric enzyme human glucokinase. Structure 12, 429–438 (2004).

16. S. Polakof, T. P. Mommsen, J. L. Soengas, Glucosensing and glucose homeostasis: from fish to mammals. Comp. Biochem. Physiol. B Biochem. Mol. Biol. 160, 123–149 (2011).

17. G. K. A. Hochberg, J. W. Thornton, Reconstructing Ancient Proteins to Understand the Causes of Structure and Function. Annu. Rev. Biophys. 46, 247–269 (2017).

18. A. C. Whittington, S. Kamalaldinezabadi, J. I. Santiago, B. G. Miller, Vertical Investigations of Enzyme Evolution Using Ancestral Sequence Reconstruction. Comprehensive Natural Products III 640–653 (2020). 10.1016/B978-0-12-409547-2.14650-5.

19. D. M. Irwin, H. Tan, Molecular evolution of the vertebrate hexokinase gene family: Identification of a conserved fifth vertebrate hexokinase gene. Comp. Biochem. Physiol. Part D Genomics Proteomics 3, 96–107 (2008).

20. M. Li, Z. Gao, Y. Wang, H. Wang, S. Zhang, Identification, expression and bioactivity of hexokinase in amphioxus: insights into evolution of vertebrate hexokinase genes. Gene 535, 318–326 (2014).

21. O. D. Madsen, Pancreas Phylogeny and Ontogeny in Relation to a “Pancreatic Stem Cell.” C. R. Biol. 330, 534 (2007).

22. N. Shiojiri, et al., Phylogenetic analyses of the hepatic architecture in vertebrates. J. Anat. 232, 200 (2018).

23. R. Annunziata, C. Andrikou, M. Perillo, C. Cuomo, M. I. Arnone, Development and evolution of gut structures: from molecules to function. Cell Tissue Res. 377, 445–458 (2019).

24. S. Nakayama, T. Sekiguchi, M. Ogasawara, Molecular and evolutionary aspects of the protochordate digestive system. Cell Tissue Res. 377, 309–320 (2019).

25. V. Hanson-Smith, A. Johnson, PhyloBot: A Web Portal for Automated Phylogenetics, Ancestral Sequence Reconstruction, and Exploration of Mutational Trajectories. PLoS Comput. Biol. 12, e1004976 (2016).

26. V. Hanson-Smith, B. Kolaczkowski, J. W. Thornton, Robustness of Ancestral Sequence Reconstruction to Phylogenetic Uncertainty. Mol. Biol. Evol. 27, 1988 (2010).

27. G. N. Eick, J. T. Bridgham, D. P. Anderson, M. J. Harms, J. W. Thornton, Robustness of Reconstructed Ancestral Protein Functions to Statistical Uncertainty. Mol. Biol. Evol. 34, 247–261 (2017).

28. R. N. Randall, C. E. Radford, K. A. Roof, D. K. Natarajan, E. A. Gaucher, An experimental phylogeny to benchmark ancestral sequence reconstruction. Nat. Commun. 7, 12847 (2016).

29. P. Pal, B. G. Miller, Activating mutations in the human glucokinase gene revealed by genetic selection. Biochemistry 48, 814–816 (2009).

30. M. Larion, B. G. Miller, 23-Residue C-terminal alpha-helix governs kinetic cooperativity in monomeric human glucokinase. Biochemistry 48, 6157–6165 (2009).

31. J. A. Martinez, M. Larion, M. S. Conejo, C. M. Porter, B. G. Miller, Role of connecting loop 1 in catalysis and allosteric regulation of human glucokinase. Protein Science 23, 915–922 (2014).

32. M. Larion, et al., Kinetic Cooperativity in Human Pancreatic Glucokinase Originates from Millisecond Dynamics of the Small Domain. Angew. Chem. Int. Ed. 54, 8129–8132 (2015).

33. S. M. Sternisha, et al., Nanosecond-Timescale Dynamics and Conformational Heterogeneity in Human GCK Regulation and Disease. Biophys. J. 118 (2020).

34. J. A. Martinez, Q. Xiao, A. Zakarian, B. G. Miller, Antidiabetic Disruptors of the Glucokinase-Glucokinase Regulatory Protein Complex Reorganize a Coulombic Interface. Biochemistry 56, 3150–3157 (2017).

35. B. H. Gordon, P. Liu, A. C. Whittington, R. Silvers, B. G. Miller, Biochemical methods to map and quantify allosteric motions in human glucokinase. Methods Enzymol. 685, 433–459 (2023).

36. C. Park, S. Marqusee, Probing the high energy states in proteins by proteolysis. J. Mol. Biol. 343, 1467–1476 (2004).

37. A. Fontana, et al., Probing protein structure by limited proteolysis. Acta Biochim. Pol. 51, 299–321 (2004).

38. Y. Chang, C. Park, Mapping Transient Partial Unfolding by Protein Engineering and Native-State Proteolysis. J. Mol. Biol. 393, 543–556 (2009).

39. Z. R. Sailer, M. J. Harms, Molecular ensembles make evolution unpredictable. Proceedings of the National Academy of Sciences 114, 201711927 (2017).

40. M. J. Harms, J. W. Thornton, Historical contingency and its biophysical basis in glucocorticoid receptor evolution. Nature 512, 203–207 (2014).

41. Z. R. Sailer, M. J. Harms, High-order epistasis shapes evolutionary trajectories. PLoS Comput. Biol. 13, e1005541 (2017).

42. T. N. Starr, J. W. Thornton, Epistasis in protein evolution. Protein Science 25, 1204–1218 (2016).

43. D. M. McCandlish, E. Rajon, P. Shah, Y. Ding, J. B. Plotkin, The role of epistasis in protein evolution. Nature 497, E1–E2 (2013).

44. M. S. Breen, C. Kemena, P. K. Vlasov, C. Notredame, F. A. Kondrashov, Epistasis as the primary factor in molecular evolution. Nature 490, 535–538 (2012).

45. C. M. Miton, N. Tokuriki, How mutational epistasis impairs predictability in protein evolution and design. Protein Science 25, 1260–1272 (2016).

46. J. M. Choi, M. H. Seo, H. H. Kyeong, E. Kim, H. S. Kim, Molecular basis for the role of glucokinase regulatory protein as the allosteric switch for glucokinase. Proc. Natl. Acad. Sci. U. S. A. 110, 10171–10176 (2013).

47. S. J. Gould, R. C. Lewontin, The spandrels of San Marco and the Panglossian paradigm: a critique of the adaptationist programme. Proceedings of the Royal Society of London - Biological Sciences 205, 581–598 (1979).

48. S. J. Gould, The exaptive excellence of spandrels as a term and prototype. Proc. Natl. Acad. Sci. U. S. A. 94, 10750–10755 (1997).

49. A. C. Whittington, D. R. Rokyta, Biophysical Spandrels form a Hot-Spot for Kosmotropic Mutations in Bacteriophage Thermal Adaptation. J. Mol. Evol. 87 (2019).

50. S. J. Gould, E. S. Vrba, Exaptation—a Missing Term in the Science of Form. Paleobiology 1, 4–15 (1982).

51. M. Kaltenbach, et al., Evolution of chalcone isomerase from a noncatalytic ancestor. Nat. Chem. Biol. 14, 548–555 (2018).

52. P. E. Wright, H. J. Dyson, Intrinsically disordered proteins in cellular signalling and regulation. Nat. Rev. Mol. Cell Biol. 16, 18–29 (2015).

53. V. J. Hilser, J. A. Anderson, H. N. Motlagh, Allostery vs. “allokairy.” Proc. Natl. Acad. Sci. U. S. A. 112, 11430–11431 (2015).

54. A. C. Whittington, A. J. Mason, D. R. Rokyta, A single mutation unlocks cascading exaptations in the origin of a potent pitviper neurotoxin. Mol. Biol. Evol. 35, 887–898 (2018).

55. A. S. Pillai, et al., Origin of complexity in haemoglobin evolution. Nature 2020 581:7809 581, 480–485 (2020).

56. J. A. Marsh, S. A. Teichmann, Structure, Dynamics, Assembly, and Evolution of Protein Complexes. Annu. Rev. Biochem. 84, 551–575 (2015).

57. C. R. Cortez-Romero, J. Lyu, A. S. Pillai, A. Laganowsky, J. W. Thornton, Symmetry facilitated the evolution of heterospecificity and high-order stoichiometry in vertebrate hemoglobin. Proc. Natl. Acad. Sci. U. S. A. 122, e2414756122 (2025).

58. N. Steube, et al., Fortuitously compatible protein surfaces primed allosteric control in cyanobacterial photoprotection. Nature Ecology & Evolution 2023 7:5 7, 756–767 (2023).

59. C. R. Baker, L. N. Booth, T. R. Sorrells, A. D. Johnson, Protein modularity, cooperative binding, and hybrid regulatory states underlie transcriptional network diversification. Cell 151, 80–95 (2012).

60. S. Rastogi, D. A. Liberles, Subfunctionalization of duplicated genes as a transition state to neofunctionalization. BMC Evol. Biol. 5, 28 (2005).

61. S. Ohno, Evolution by Gene Duplication (Springer Berlin Heidelberg, 1970).

62. J. E. Wilson, Isozymes of mammalian hexokinase: structure, subcellular localization and metabolic function. J. Exp. Biol. 206, 2049–57 (2003).

63. J. E. Wilson, “The Hexokinase Gene Family” in Glucokinase and Glycemic Disease: From Basics to Novel Therapeutics, (KARGER, 2004), pp. 18–30.

64. T. Ureta, The comparative isozymology of vertebrate hexokinases. Comparative Biochemistry and Physiology -- Part B: Biochemistry and 71, 549–555 (1982).

65. D. Liebschner, et al., Macromolecular structure determination using X-rays, neutrons and electrons: recent developments in Phenix. Acta Crystallogr. D Struct. Biol. 75, 861–877 (2019).

