## Supplementary Text for "Evolution of Protein Regulation in the Vertebrate Glucose Sensor"

**This PDF file includes:**

Materials and Methods
Figures S1 to S71
Table S1 to S3
SI References

**Other supporting materials for this manuscript include the following:**

Phylogenetics Files
PDB Validation File

### Supporting Information Text

### Materials and Methods

**Phylogenetics** Sequences of extant GCKs and GKRPs were obtained from the NCBI non-redundant protein database and EMBL's Ensembl in 2018. Redundant sequences, incomplete sequences, sequences with ambiguous codons, and sequences not obtained through DNA sequencing were discarded. In total, 194 sequences from across Deuterostomia were used in the GCK reconstruction, with the echinoderm *Apostichopus japonicus* GCK set as the outgroup. 98 sequences from Amoebas and Holozoa were used in the GKRP reconstruction. PhyloBot was used to perform sequence alignments, phylogenetic reconstruction, and estimation of ancestral sequences (*1*). PhyloBot takes a set of input sequences and aligns them with MSAprobs and Muscle, estimates trees using RAxML with a variety of commonly used amino acid evolution models with two types of rate heterogeneity categories (CAT and GAMMA). CODEML is used to infer ancestral sequences. PhyloBot used MSAprobs for the GCK and GKRP alignments. The 148 GCK amino acid sequences from chordates yielded an alignment length of 513 residues anchored by 61 completely conserved sites. There were a few species' GCKs with relatively long N- and C-terminal deletions leading to 258 informative sites, i.e. sites with no indels and not completely conserved. The GKRP alignment length was 1057 residues anchored by 32 completely conserved sites. The 98 GKRP sequences used span all of eukaryote evolution and thus show higher divergence. There were 416 informative sites. Separate reconstructions were performed for the GCK and GKRP sequences, and gaps were reconstructed according to Kaltenbach, et al (*2*).

PhyloBot provides the output for ten alignment/model combinations. We used the "best" (top ranked model) and "worst" (lowest ranked model) for robustness analysis. The best fit alignment for both the GCK and GKRP sequences was MSAprobs. The GCK alignment best model was JTT with CAT and the worst was WAG with GAMMA. The GKRP alignment best model was LG with CAT and the worst was WAG with GAMMA. PhyloBot assesses topology using the SH-aLRT in PhyML GCK and GKRP trees showing branch lengths and SH-aLRT branch supports are shown in Figs. S15-S18.. PhyloBot uses CODEML to estimate ancestral sequences using the empirical Bayesian method and reconstructs indels by parsimony. Posterior probabilities across sites for chordate, vertebrate, and gnathostome ancestral GCKs and GKRPs for the best model are given as histograms (Fig. S18-S23). Average posterior probabilities for the best GCK and GKRP reconstructions are shown in Table S1.

Robustness of phenotypes was determined by generating and assaying the best and worst reconstructions of the chordate and vertebrate GCK ancestors from Phylobot and the AltAll (*3*) of the best Phylobot reconstructions of both of those ancestors. For our AltAll version, sites which were reconstructed with a posterior probability below 0.6 (P1) were considered to be ambiguously reconstructed. The posterior probability of the second most probable amino acid at that site was named P2. For the ambiguously reconstructed sites, if (P2/P1) > 0.5, then the most probable amino acid was replaced by the second most probable amino acid for that site. We must acknowledge that this is a less conservative version than 0.2 posterior probability cutoff. Our version has 58 residues that are different from the best cGCK, whereas the 0.2 AltAll would have 68 differences (sequence identities and number of residue differences can be found in Table S2). Although unlikely, these differences could result in uncertainty regarding the number of ambiguously reconstructed sites and the properties of the alternate ancestor. We do note, however, that the functional properties of the "worst" ancestors were robust to phenotypic uncertainty. The best and worst PhyloBot cGCKs were 97.58 % identical. The best and AltAll cGCKs were 87.25 % identical. These were phenotypically identical. Similarly, the best and worst vGCKs (97.85 % identical) were phenotypically identical. The maximum likelihood version of cGCK (88.11 % identical to the best cGCK) from the FastML reconstruction was also identical (Figs. S49 & S50). There is an incidence of discordance of the GCK best tree with the species tree where the coelacantch (*Latimeria chalumnae*) is sister taxa to the jawed vertebrates while the elasmobranch (*Callorhinchus milii*) is sister taxa to the tetrapods. This discordance may arise from long-branch attraction and/or the low sequence sampling density available for these species and regions of the tree. However, our tests of robustness described above demonstrate that our phenotypic reconstructions are robust to relatively large sequence variation. The relative position of these two discordant taxa would likely have a smaller effect on the sequence than the difference between the best and AltAll versions of cGCK. Additionally, the FastML tree (see SI_alignment_files) has frog GCKs (*Xenopus*) as the sister taxa to the jawed vertebrates and that cGCK version is identical to the others, further showing that our reconstruction is robust to some amount of gene tree/species tree discordance. Alignments with accession number, trees, and ancestral sequences are provided in SI files (SI_phylogenetics_files).

**Recombinant protein production and purification** Ancestral GCK and GKRP coding sequences cloned into pET-22b (+) and codon-optimized for expression in *Escherichia coli* were ordered from GenScript (Piscataway, NJ). N-terminal hexahistidine-tagged recombinant GCKs were expressed in BL21 (DE3) cells as previously described (*5*). Recombinant GCKs were purified via $Ni^{2+}$-affinity and size exclusion chromatography (SEC) as previously described (*6*). C-terminal hexahistidine-tagged recombinant GKRPs were expressed in *E. coli* BM5340 (DE3) cells. BM5340 (DE3) is a glucokinase deficient strain to avoid contaminating activity from endogenous *E. coli* glucokinases (*7*). Cultures were inoculated to an $OD_{600}$ of 0.01 in LB supplemented with ampicillin (100 µg/mL), kanamycin (20 µg/mL), and chloramphenicol (10 µg/mL). Cells were shaken at 250 rpm, 37 °C until the OD600 reached 0.50, at which point the temperature was reduced to 16 °C. Once the incubator reached 16 °C, IPTG (0.5 mM) was added to induce gene expression. Growth continued for 40 h, after which cells were harvested by centrifugation for 10 min at 6,000 × g and 4 °C. GKRPs were purified via $Ni^{2+}$-affinity chromatography and SEC as described previously (*8–10*). When working with vGKRP, chromatography buffers were supplemented with L-arginine (50 mM), L-glutamate (50 mM), and DTT (10 mM) to mitigate aggregation and increase protein stability. Chemicals for protein purification were purchased from Thermo Fisher Scientific, Sigma-Aldrich, and Gold Biotechnology.

**Enzyme kinetics, inhibition assays, and limited proteolysis studies** GCK activity was measured via an enzyme-coupled spectrophotometric assay in which the production of glucose-6-phosphate (G6P) is linked to the conversion of $NADP^+$ to NADPH via glucose 6-phosphate dehydrogenase (G6PDH) from *Leuconostoc mesenteroides* (Sigma-Aldrich). Measurements were performed in a 1 mL assay containing 200 mM HEPES pH 8.0, 50 mM NaCl, 10 mM DTT, 2 mM $NADP^+$, 7.5 units of G6PDH with variable concentrations of ATP or glucose. $MgCl_2$ was included at concentrations 1 mM higher than ATP concentrations. Assays were performed using a UV-Visible spectrophotometer at 340 nm (CARY 100 Bio). The reaction mixture was equilibrated for 2 minutes at 25°C, after which time the baseline was recorded for 1 min, and then reactions were initiated via ATP addition. Each experiment was carried out at least twice, and at least two replicates were performed at each substrate concentration in each experiment. Data were fit to the Michaelis-Menten equation or the Hill equation using GraphPad Prism. The reported kinetic parameters ($k_{cat}$, $K_{0.5}$, Hill coefficient) represent the average and standard deviation from two individual experiments.

To evaluate GKRP inhibition of GCK, steady-state kinetics assays were performed using GCKs at a final concentration of 25–100 nM, such that the uninhibited rate of glucose conversion corresponded to a rate of G6P production of ~540 nM per second. Ancestral GCKs were mixed with ancestral GKRPs at various GKRP concentrations (0–100 µM), and the proteins were incubated in GKRP SEC buffer supplemented with 10 mM DTT and, if applicable, sorbitol 6-phosphate (final assay concentration of 2 mM) for 5 min at 25 °C to allow binding to reach equilibrium. Incubation took place in a 100 µL quartz cuvette to minimize the risk of sample loss during transfer. Following equilibration, 1 unit of G6PDH was added to the reaction mixture, followed by a master mix containing HEPES (250 mM, pH 7.1), KCl (25 mM), NADP+ (0.5 mM), DTT (10 mM), $MgCl_2$ (6 mM), and glucose. The final concentration of glucose was 5 mM when working with vGCK, gGCK, or tGCK, and 50 µM when working with cGCK, due to cGCK's lower $K_{0.5}$ value for glucose. The reaction mixtures were incubated 3 min prior to initiating the reaction with ATP (5 mM). The rate of glucose conversion was plotted as a function of GKRP concentration and the $IC_{50}$ values were calculated using GraphPad PRISM by fitting the data to a sigmoidal ligand dose-response. Data that could not be fitted with an $R^2$ value of at least 0.9 were fit to a linear equation instead to demonstrate lack of inhibition and were interpreted as having an $IC_{50}$ >1500 µM, the maximum $IC_{50}$ value possible in the assay.

Proteolysis assays of unliganded GCKs were performed with thermolysin (Sigma-Aldrich) as previously described (*11*, *12*). Plots of percentage of GK activity versus incubation time with thermolysin were fit to a single-phase exponential decay equation using GraphPad Prism to extract an observed rate ($k_{obs}$) that was converted to half-life.

**Crystallography and structure determination** Diffraction quality crystals of FastML-cGCK were obtained via the hanging drop method by mixing 1 µL of protein solution (30 mg/mL, purified as described above) with 1 μL of a well solution (0.1 M Tris-HCl pH 7.5, 25% w/v PEG 2000 MME, and 0.3 M sodium acetate) and equilibrating the drop against the same well solution at room temperature for 3-5 days. Crystals were cryoprotected by soaking briefly in the mother liquor supplemented with 15% v/v glycerol, before being flash frozen in liquid nitrogen on a loop. 1200 frames of diffraction data over 240 degrees were collected from a single frozen crystal at the NSLS-II beamline 17-ID-2 equipped with the FMX-16MEiger detector. Following integration and scaling of data using HKL2000 (*13*), the structure of cGCK was determined using molecular replacement via the MRage package in Phenix (*14*) by serially searching for the solution using the individual domains of human GCK (PDB 1V4S) as probes. Structure refinement was carried out with Phenix until satisfactory crystallographic residual factors and stereochemical parameters were reached (Table S3).

**Hydrogen-deuterium exchange mass spectrometry** Samples of cGCK and vGCK were produced and purified as described above. Protein samples were concentrated to 50 μM using an Amicon Ultra-15 Centrifugal Filter Unit (MilliporeSigma). 5 μL of each sample was added to 45 μL potassium phosphate buffer (25 mM, pH 7.0 prepared in $H_2O$ or $D_2O$) and the sample was incubated at 25˚C in a water bath. After a set incubation period, the sample was quenched with 25 μL of an ice-cold solution containing 200 mM TCEP, 8M urea and 1.0% formic acid (pH ~ 2.3). A 25 μL aliquot of ice-cold 1.0% formic acid was added to the sample, followed by addition of 400 μL of ice-cold 0.1% formic acid. 100 μL aliquots were injected onto a Waters HDX Manager UPLC system coupled to a Xevo G2-XS Q-Tof mass spectrometer. Protein samples were digested with an in-line pepsin column (Waters Enzymate BEH pepsin column, 2.1 x 30 mm, maintained at 15 °C), followed by desalting (2.1 x 5 mm ACQUITY UPLC BEH C-18 VanGuard trapping column) and then separation on 1.0 x 100 mm ACQUITY UPLC BEH C-18 analytical column maintained at 4 °C. Peptides were separated with a 5-minute gradient from 5%-85% acetonitrile in 0.1% formic acid and a flow rate of 30 μL/min. Peptide ion and $MS^E$ fragment data from non-deuterated samples were analyzed with ProteinLynx Global Server software (Waters Corp., Milford, MA) to generate a peptide coverage map. Mass shits in deuterium exchanged peptides were determined using DynamX 3.0 (Waters Corp., Milford, MA) to generate deuterium incorporation plots. The maximum relative exchange for each peptide was determined using a fully deuterated control sample. Data and error bars shown are the average of three replicate experiments (Fig. 2.; Fig. S52).

**Nuclear Magnetic Resonance** Methyl groups on isoleucine side chains of cGCK and vGCK were radio-labeled using the isoleucine precursor, alpha-ketobutyric acid (methyl-$^{13}C$, 99%; 3,3-$D_2$, 98% CDLM-7318-0; Cambridge Isotope Laboratories) as previously described (*11*, *15*, *16*). Enzymes were purified as described above and dialyzed against 25 mM potassium phosphate pH 8.0, 50 mM KCL and 10 mM DTT at 4°C. All spectra were acquired on a Bruker AVANCE III 700-MHz NMR spectrometer equipped with a TCI cryogenic probe. 2D $^1H$-$^{13}C$ ALSOFAST-HMQC NMR experiments (pulse program: afhmqcgpphsf) were used to obtain proton and carbon correlations. The data were recorded as matrices of 1536 × 160 complex datapoints. The spectral widths were 15.0012 ppm and 22.0000 ppm for F2 and F1 directions respectively. The recycling delay d1 was 0.3 seconds. Proton chemical shifts were referenced using DSS as an internal reference, while carbon-13 chemical shifts were reference indirectly. All spectra were apodised with a QSIN function with an SSB of 3.5 in each dimension before zero filling. The experiments were performed at 298 K. The spectra were collected using either 512 or 1024 scans depending on sample concentration. Site-specific assignments of Ile resonances were accomplished by single-site substitution of Ile to Val followed by expression, purification, and the collection of the 2D spectra to identify the missing cross peak, as previously described (*11*, *15*, *16*). NMR data processing and spectral analysis were performed using TopSpin 4.1.4 program. For chemical shift perturbation analysis, the peak picking function of TopSpin was used. For the peaks that were not automatically picked, peak selection was performed manually. The data were transferred to an excel sheet for further analysis. The chemical shift perturbation (CSP) upon glucose binding for each Ile residue were calculated using the equation CSP = $\{0.5 \times [(Dd_{1H})^2 + (0.25 \times Dd_{13C})^2]\}^{0.5}$, where $Dd_{1H}$ is the change in chemical shift in $^1H$ and $Dd_{13C}$ is the change in chemical shift in $^{13}C$.

Replicate 1

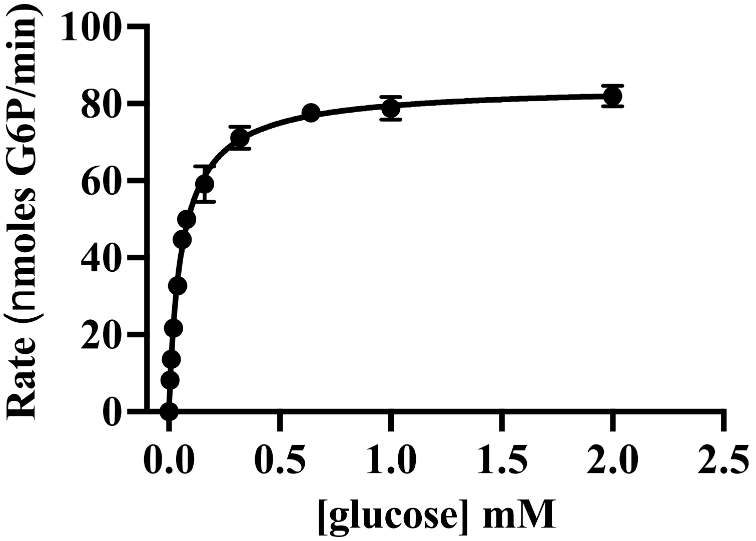

Replicate 2

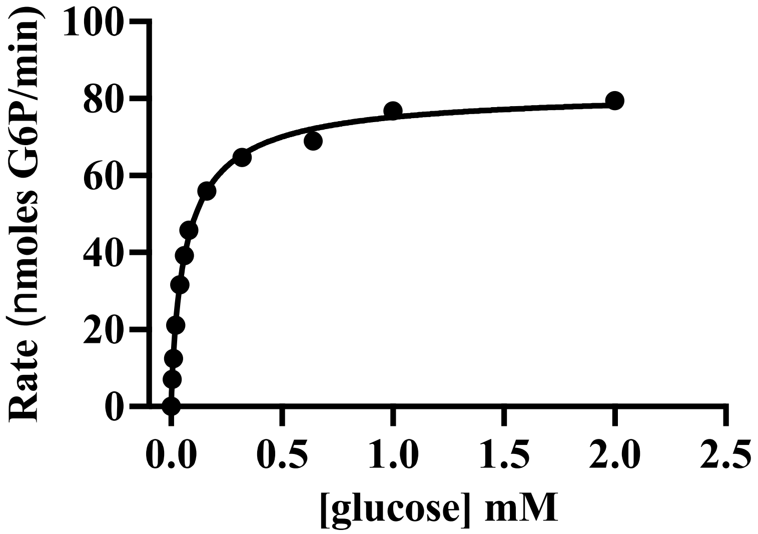

**Fig. S1**. Steady-state kinetic assays of *Ciona intestinalis* GCK with variable glucose concentrations.

Replicate 1

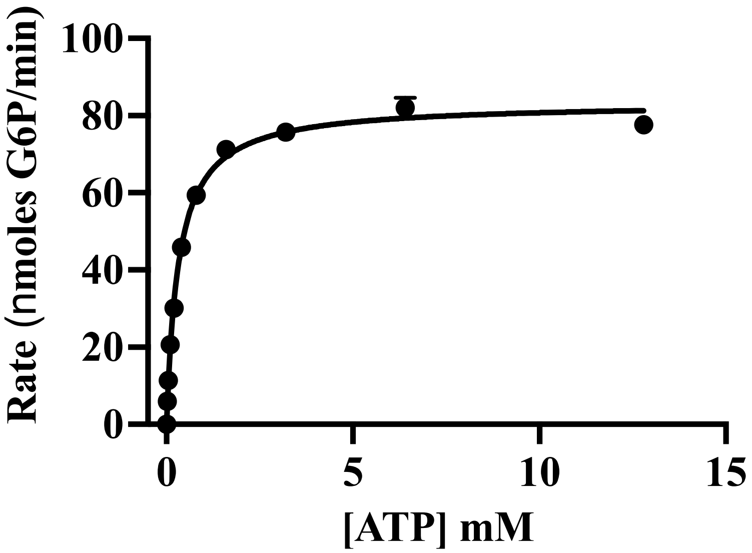

Replicate 2

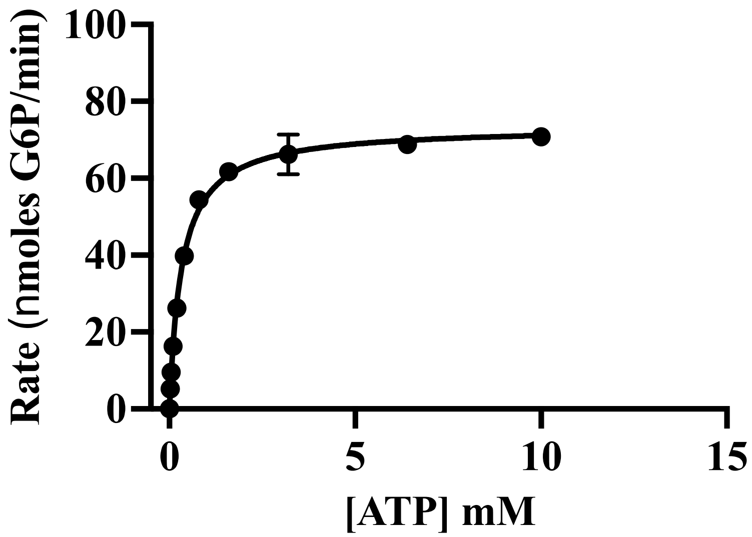

**Fig. S2**. Steady-state kinetic assays of *C. intestinalis* GCK with variable ATP concentrations.

Replicate 1

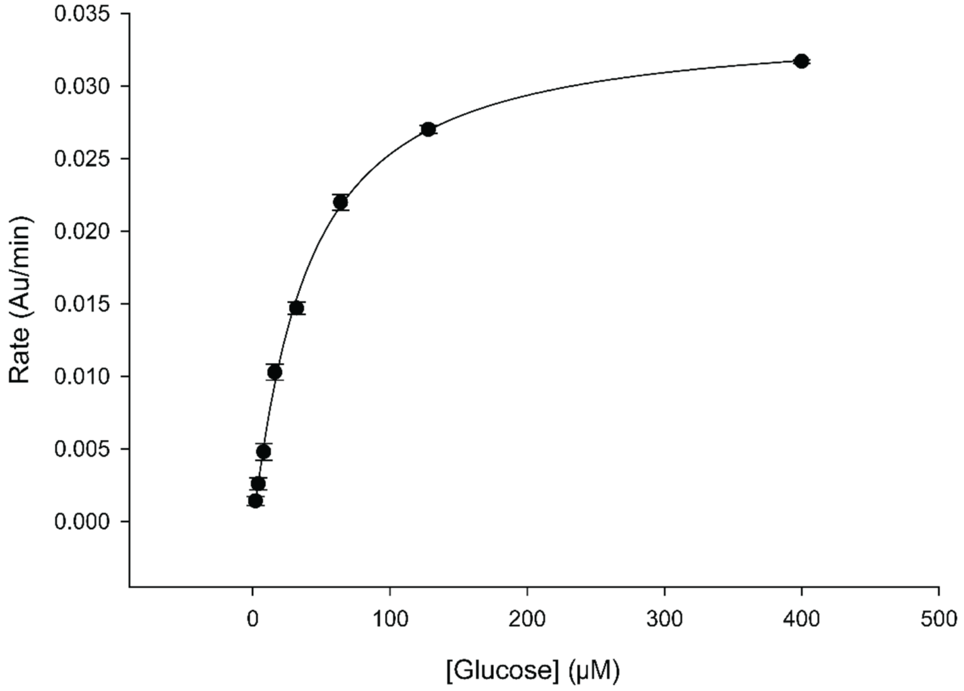

Replicate 2

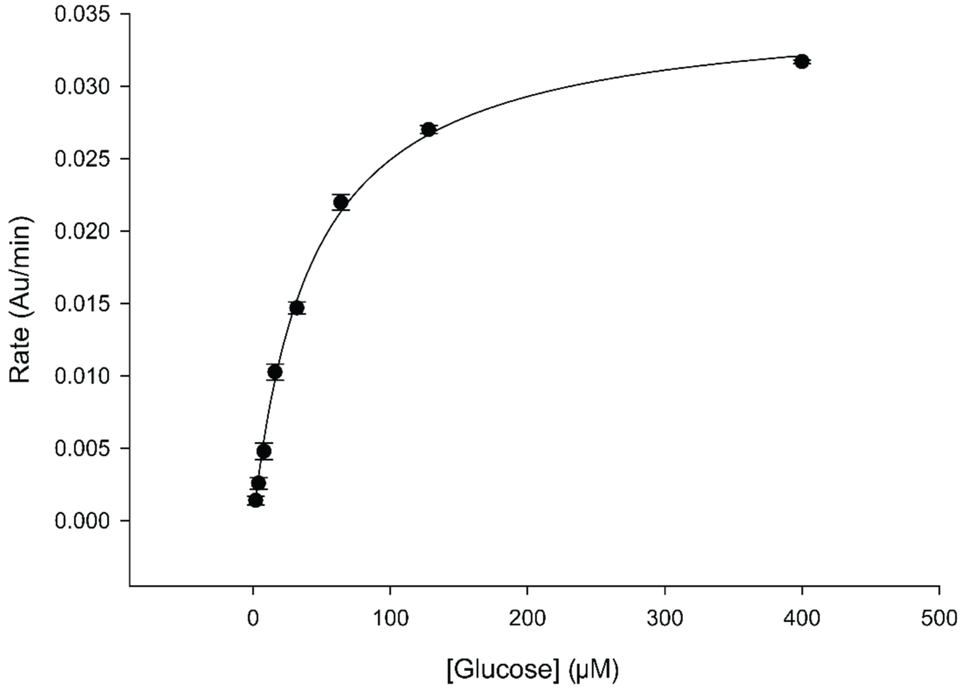

**Fig. S3**. Steady-state kinetic assays of *Branchiostoma japonicum* GCK with variable glucose concentrations.

Replicate 1

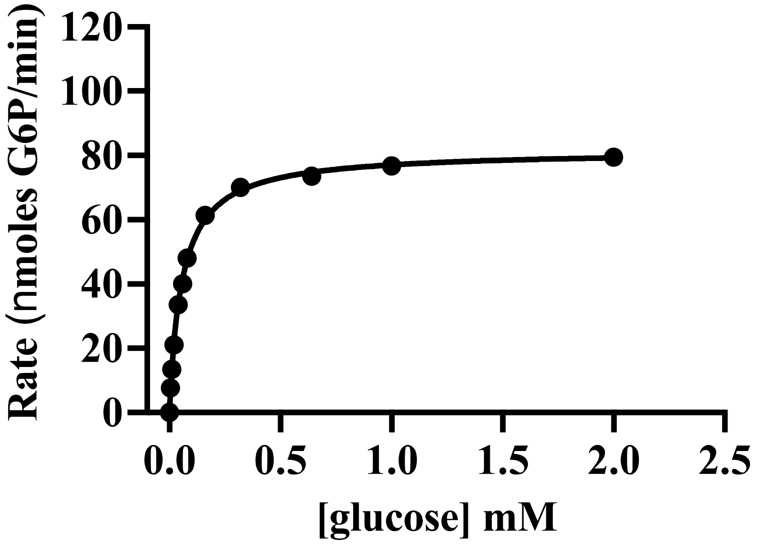

Replicate 2

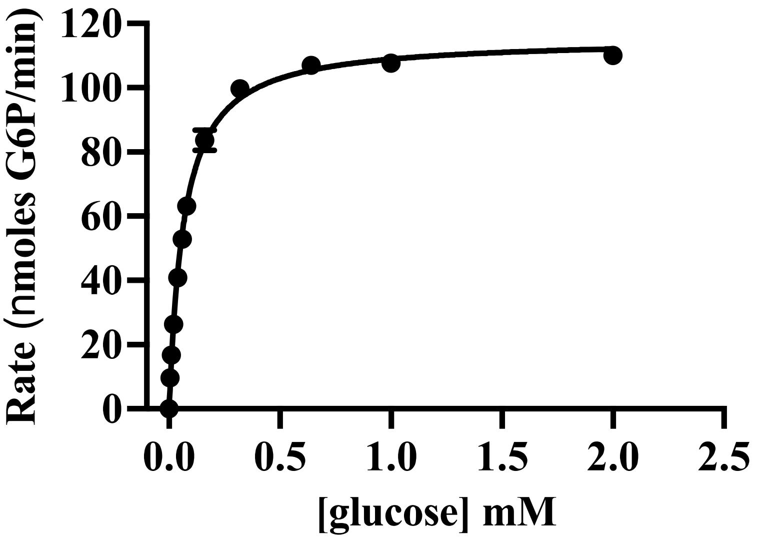

**Fig. S4**. Steady-state kinetic assays of *Branchiostoma belcheri* GCK with variable glucose concentrations.

Replicate 1

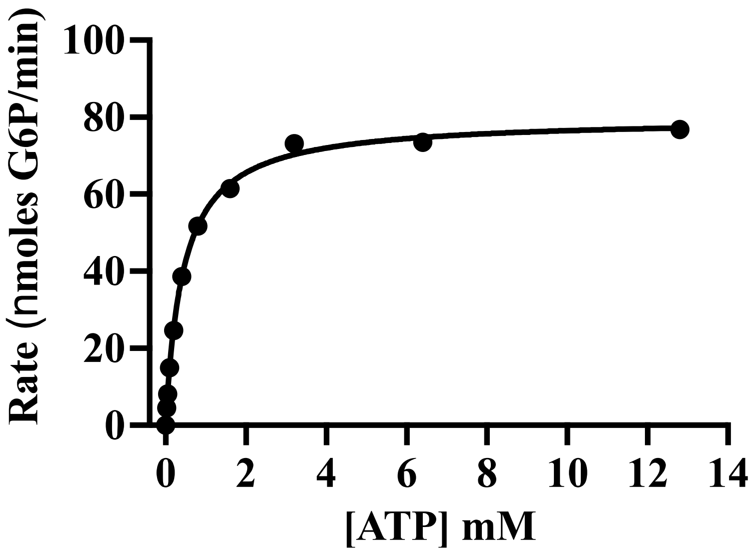

Replicate 2

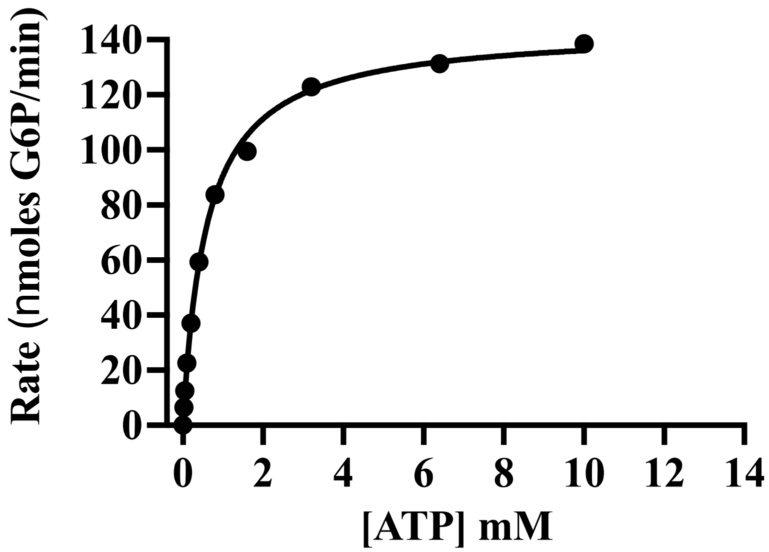

**Fig. S5**. Steady-state kinetic assays of *B. belcheri* GCK with variable ATP concentrations.

Replicate 1

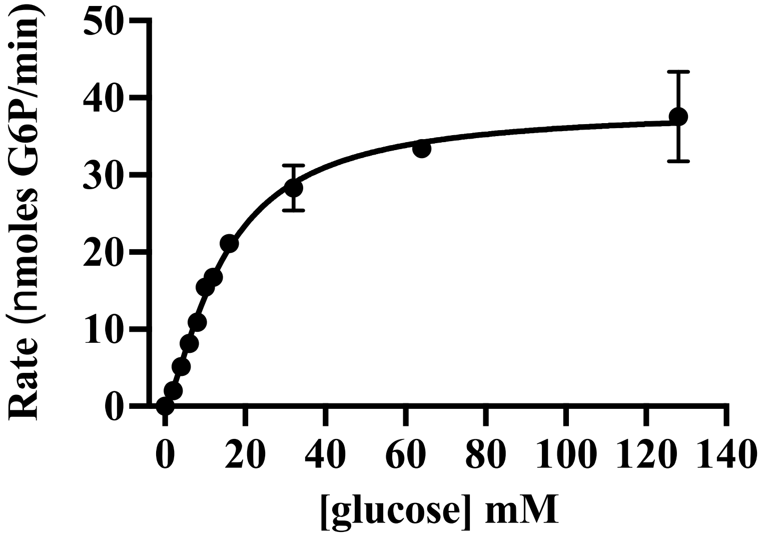

Replicate 2

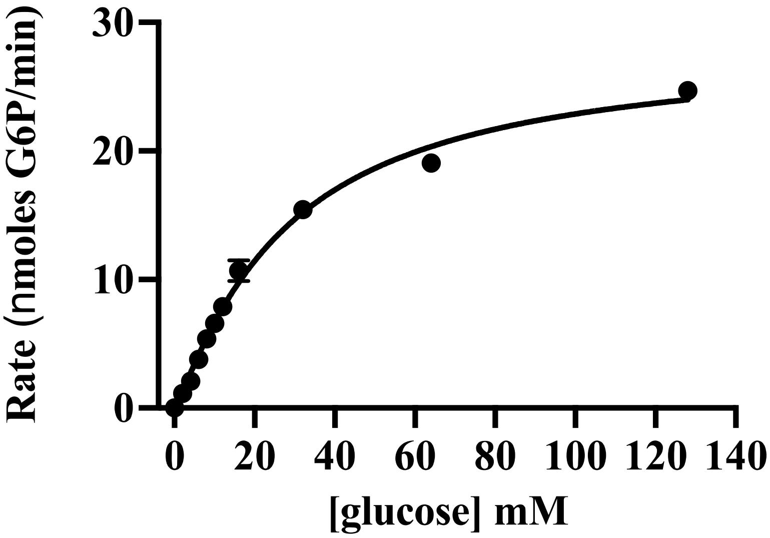

**Fig. S6**. Steady-state kinetic assays of *Latemeria chalumnae* GCK with variable glucose concentrations.

Replicate 1

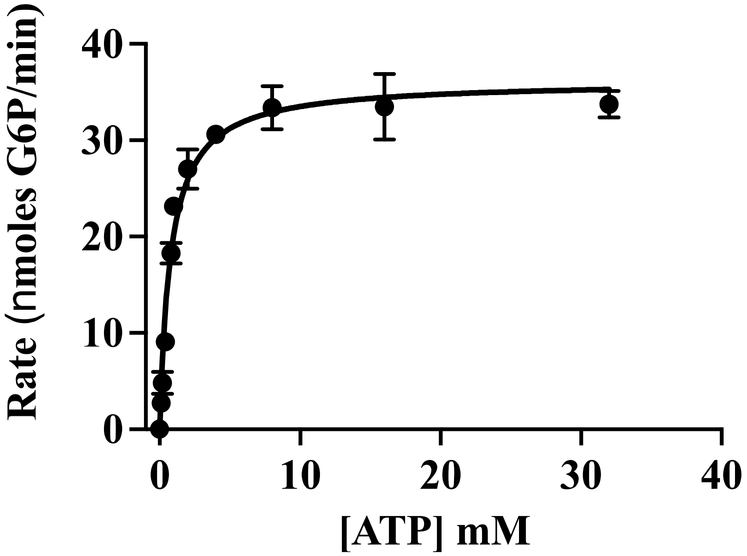

Replicate 2

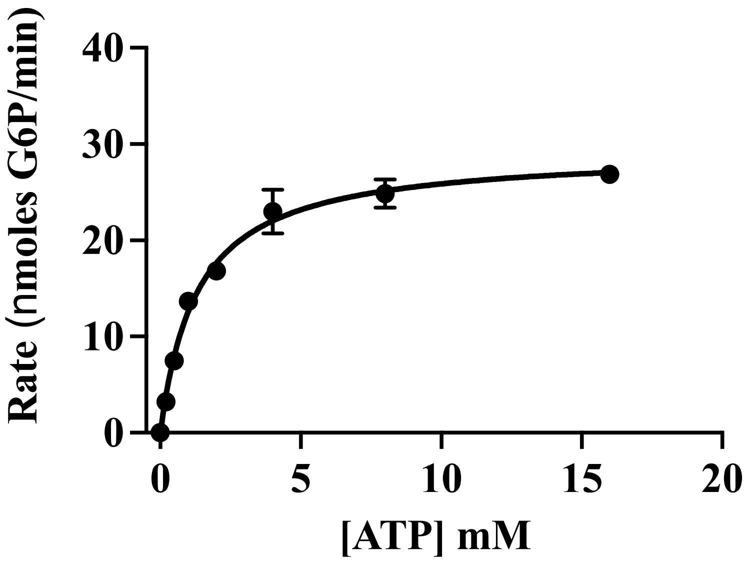

**Fig. S7**. Steady-state kinetic assays of *L. chalumnae* GCK with variable glucose concentrations.

Replicate 1

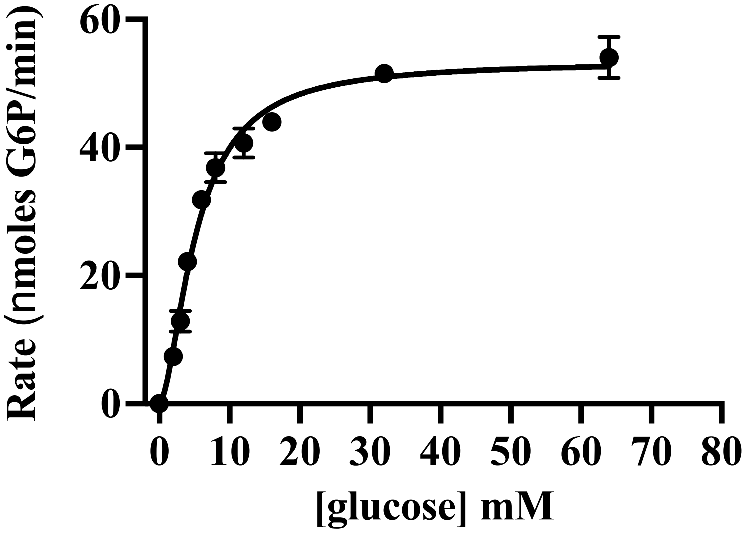

Replicate 2

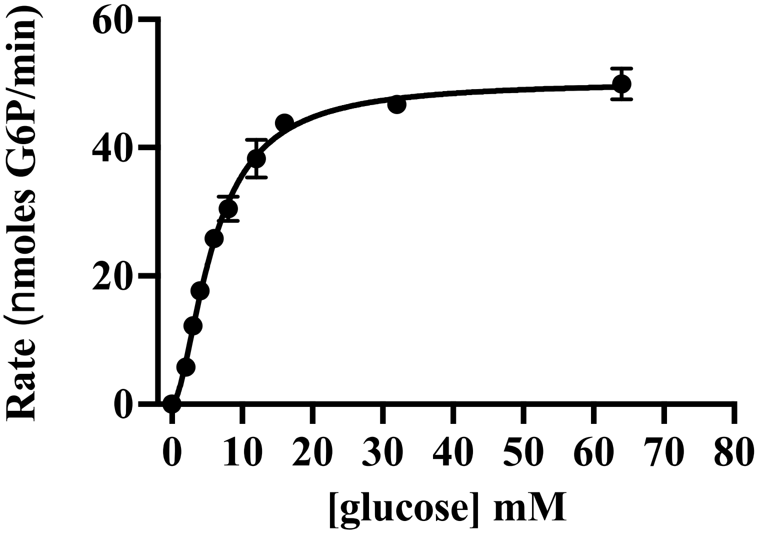

**Fig. S8**. Steady-state kinetic assays of ***Danio*** *rerio* GCK with variable glucose concentrations.

Replicate 1

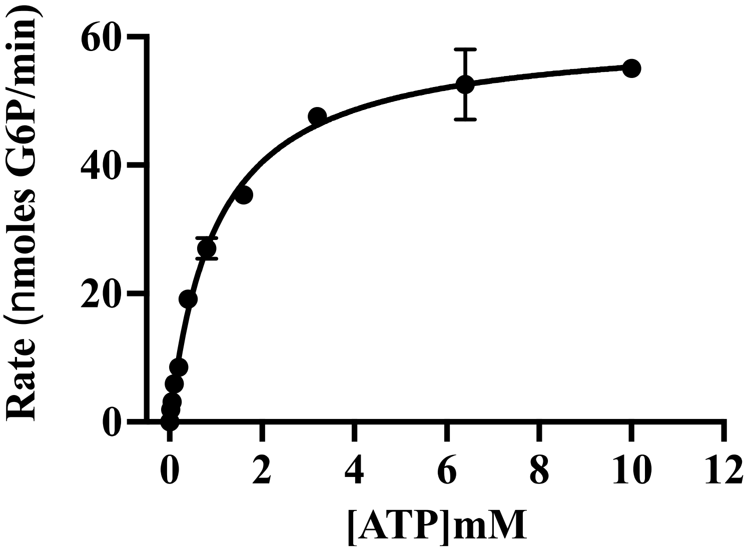

Replicate 2

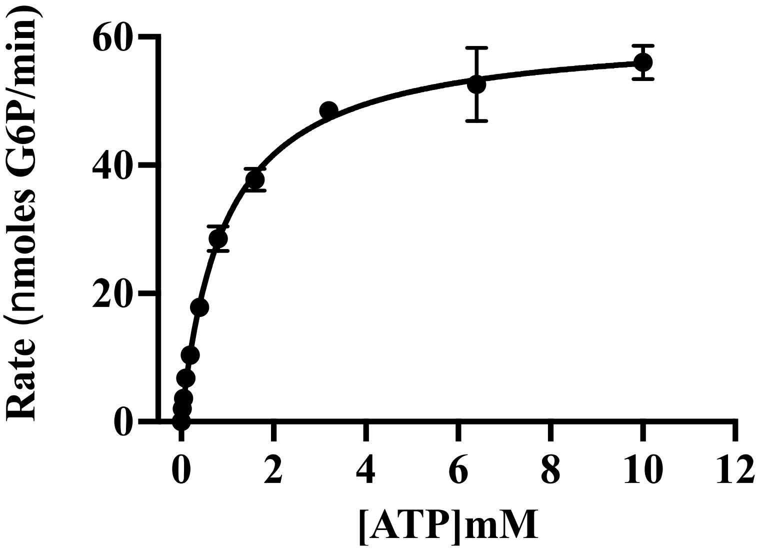

**Fig. S9**. Steady-state kinetic assays of *D. rerio* GCK with variable ATP concentrations.

Replicate 1

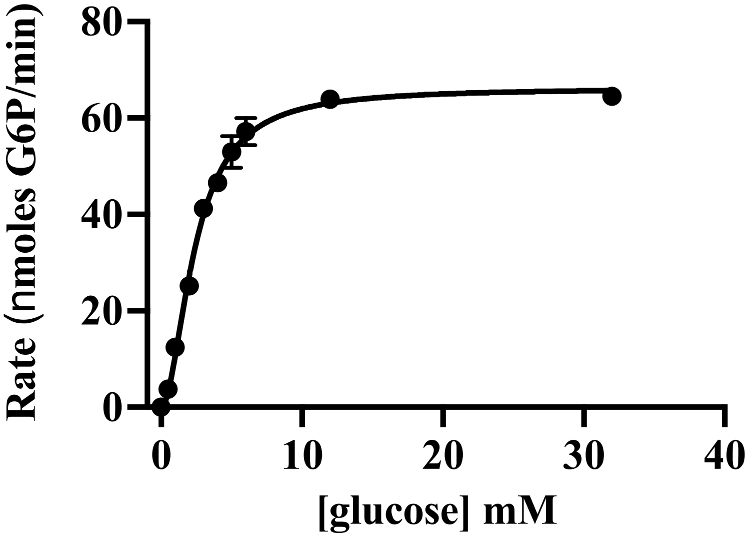

Replicate 2

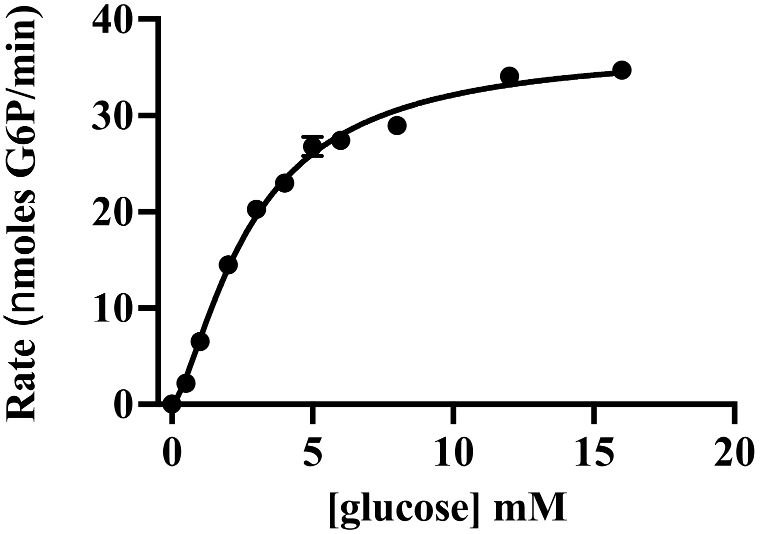

**Fig. S10**. Steady-state kinetic assays of *Xenopus laevis* GCK with variable glucose concentrations.

Replicate 1

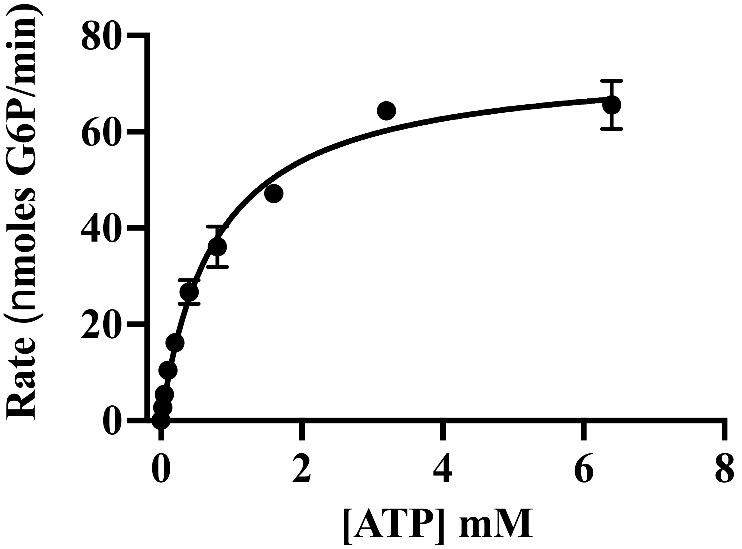

Replicate 2

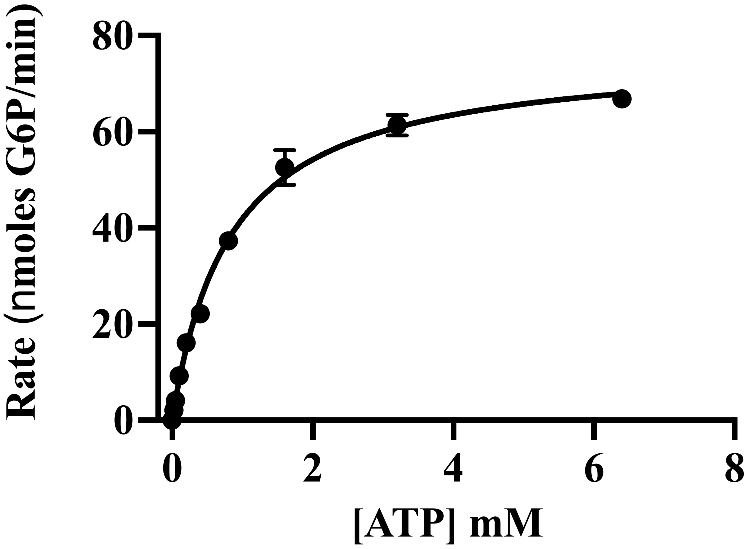

**Fig. S11**. Steady-state kinetic assays of *X. laevis* GCK with variable ATP concentrations.

Replicate 1

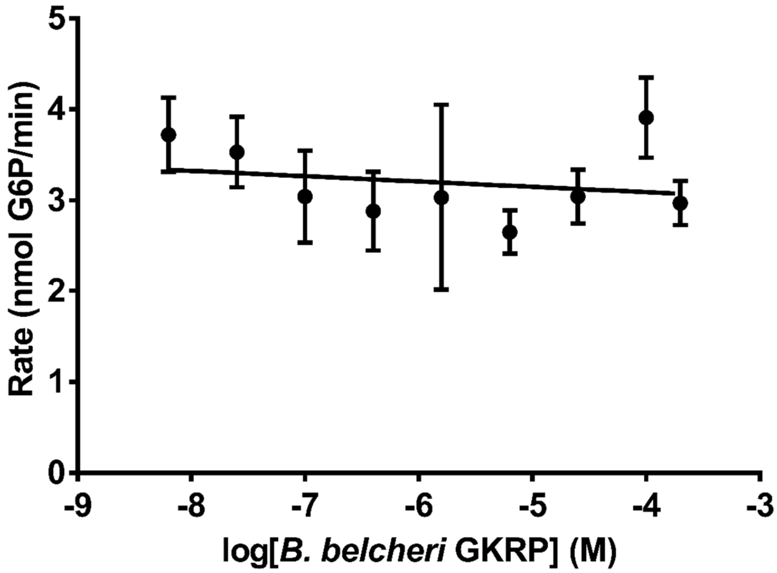

Replicate 2

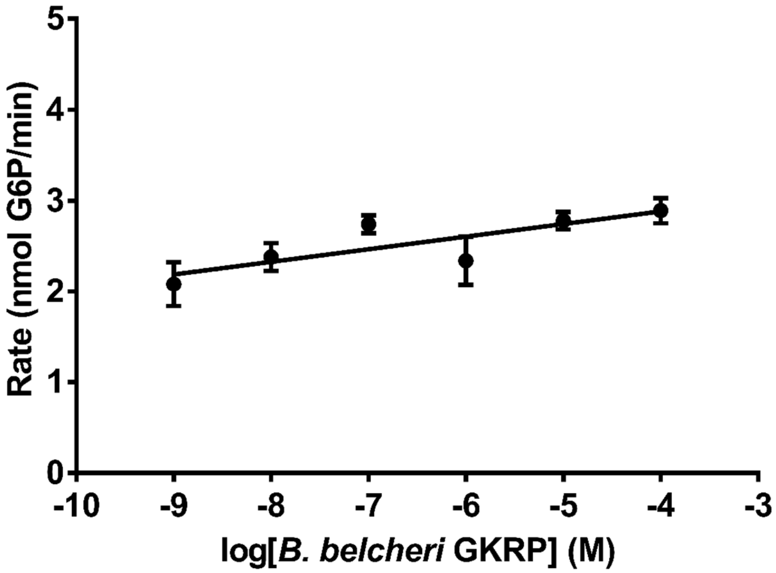

**Fig. S12**. GCK and GKRP protein pairs from *B. belcheri* do not form an inhibitory interaction. Each data point represents the average of three technical replicates. Error bars represent standard deviations. Where not visible, the error bars are smaller than the data points.

Replicate 1

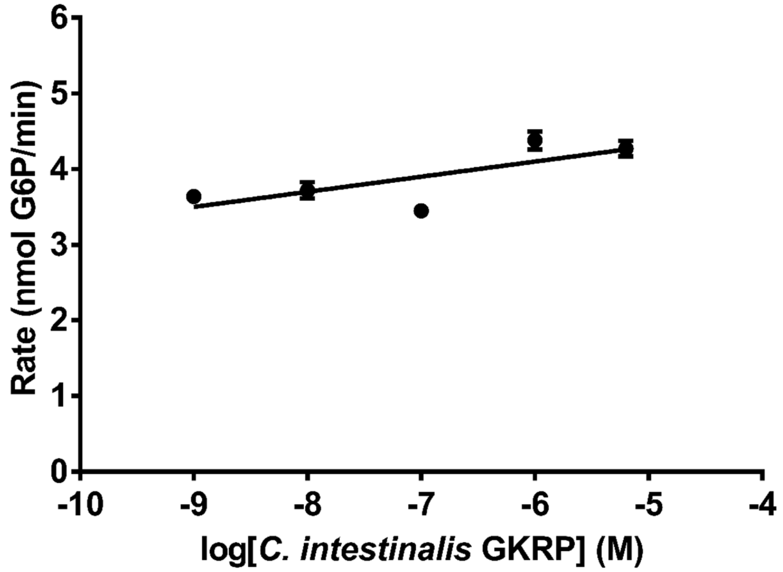

Replicate 2

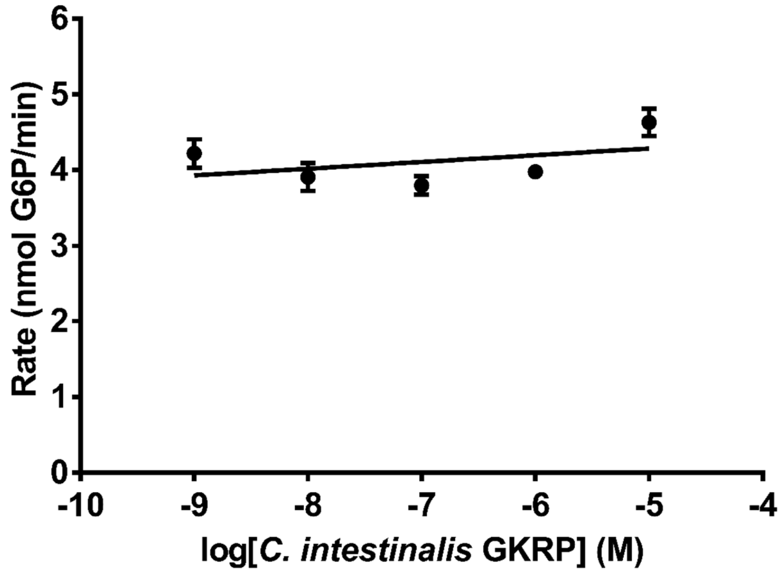

**Fig. S13**. GCK and GKRP protein pairs from *C. intestinalis* do not form an inhibitory interaction. Each data point represents the average of three technical replicates. Error bars represent standard deviations. Where not visible, the error bars are smaller than the data points.

Replicate 1

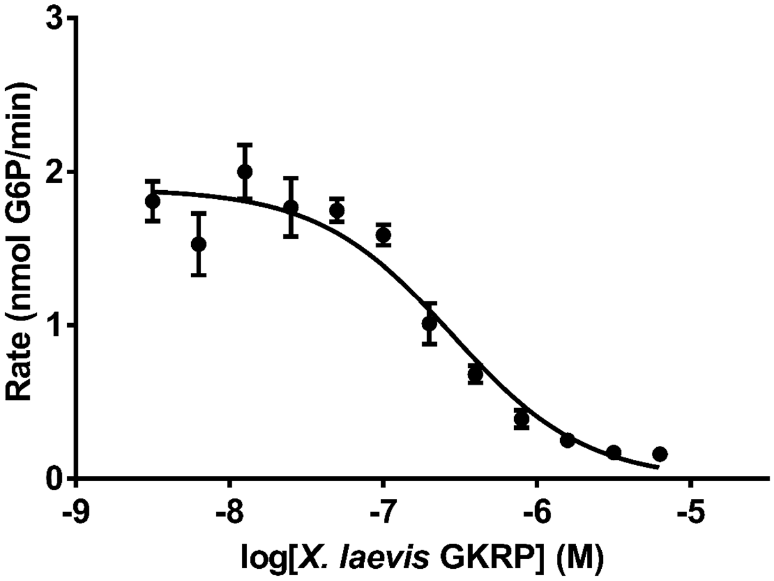

Replicate 2

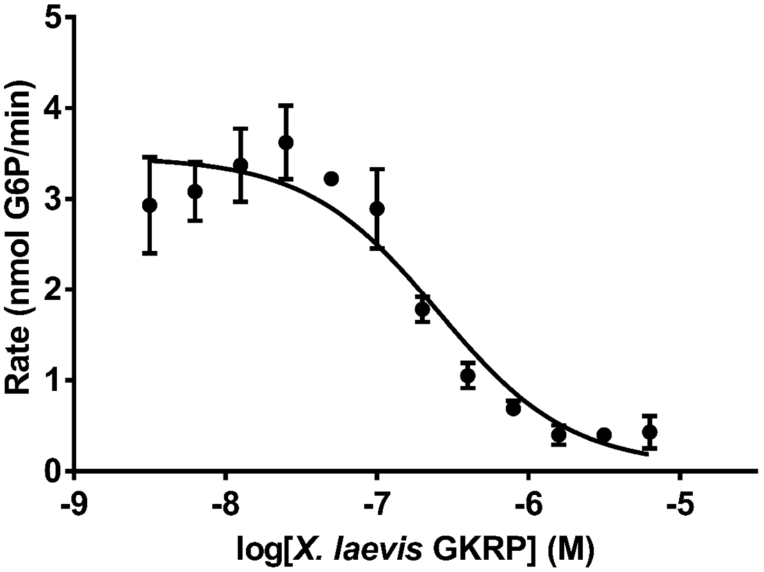

**Fig. S14**. GCK and GKRP protein pairs from *X. laevis* form an inhibitory interaction. Each data point represents the average of three technical replicates. Error bars represent standard deviations. Where not visible, the error bars are smaller than the data points.

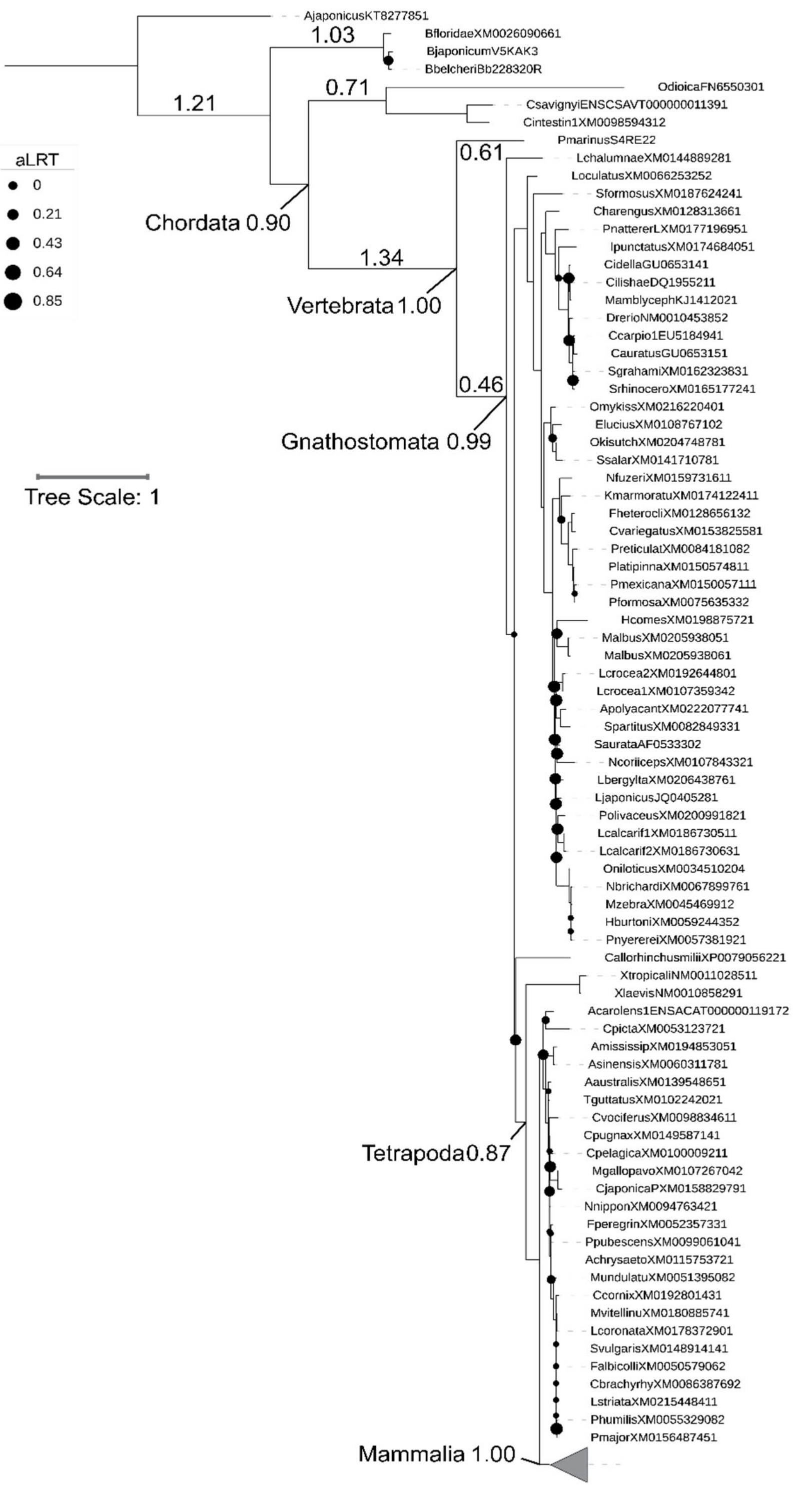

**Fig. S15**. GCK tree used for ASR with ancestral nodes labeled. Mammal clade collapsed for clarity. Branch lengths and SH-aLRT values are labeled for ancestral nodes surrounding functional transition points. SH-aLRT values ≥ 0.85 not shown for clarity. An SH-aLRT value of 0 indicates a very short branch length or that the presence/absence of the branch does not alter the likelihood of the tree.

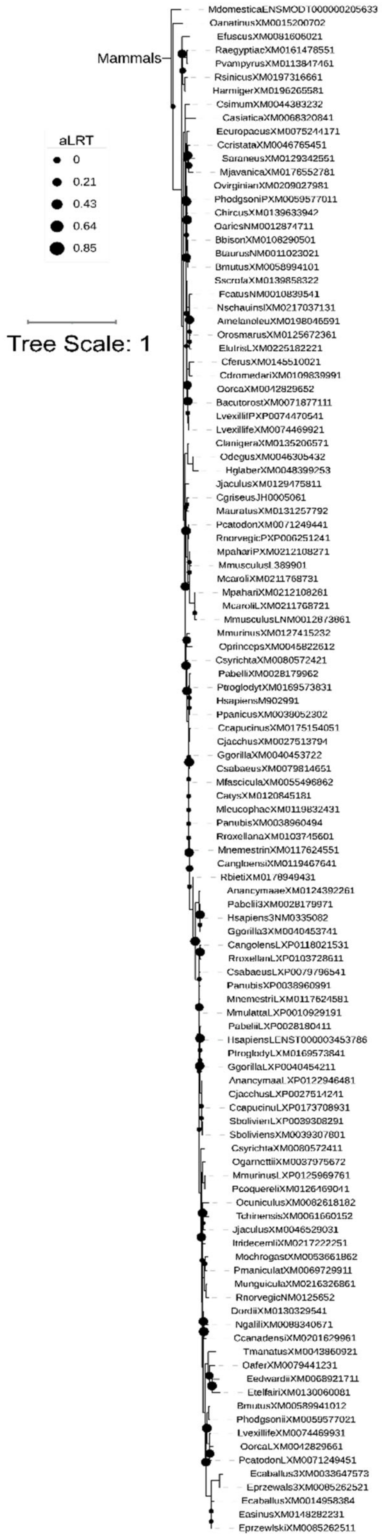

**Fig. S16**. The mammal section of the GCK tree shown in Fig. S15.

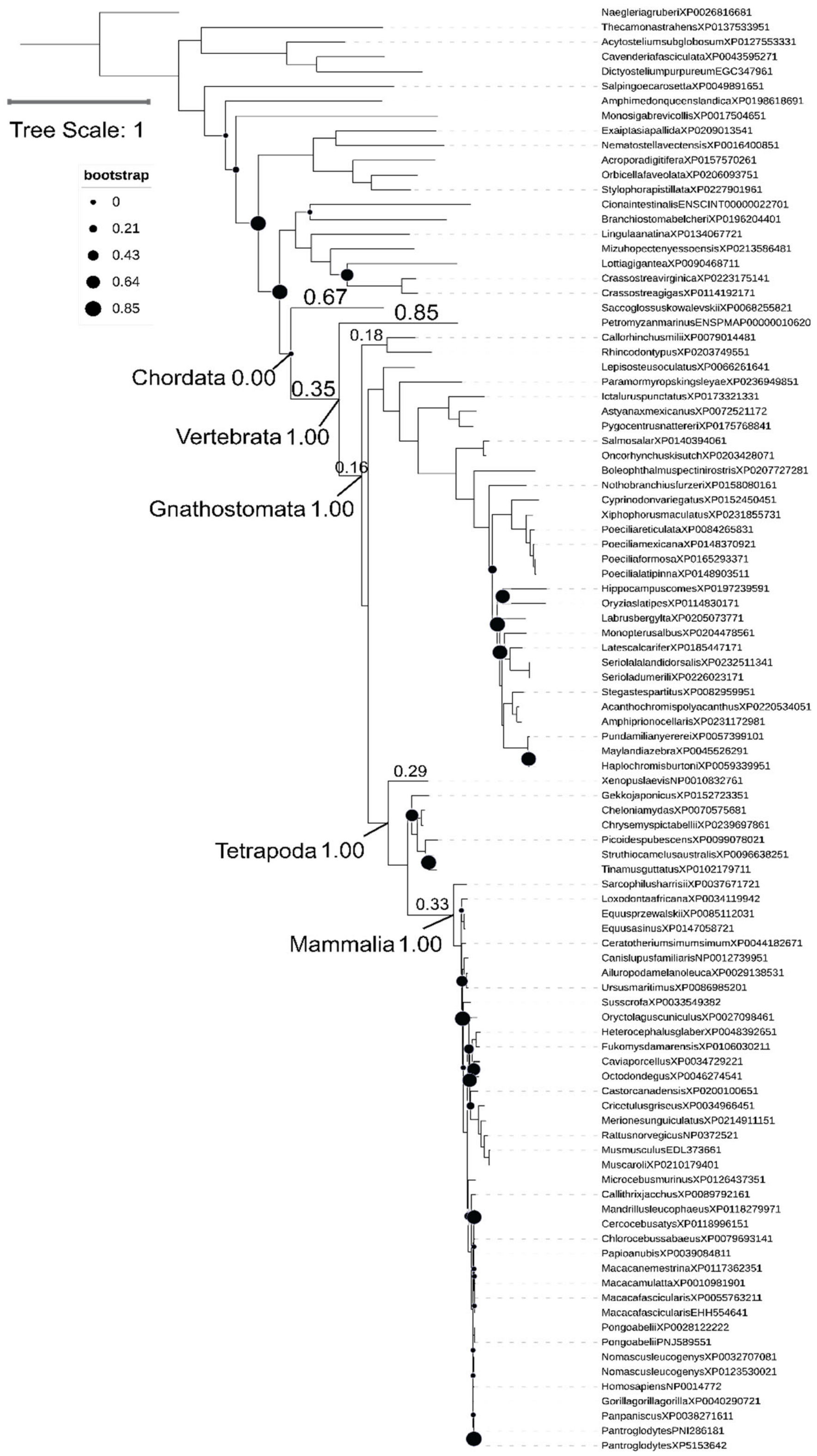

Fig. S17. GKRP tree used for ASR with ancestral nodes labeled. Branch lengths and SH-aLRT values are labeled for ancestral nodes surrounding functional transition points. SH-aLRT values ≥ 0.85 not shown for clarity. An SH-aLRT value of 0 indicates a very short branch length or that the presence/absence of the branch does not alter the likelihood of the tree.

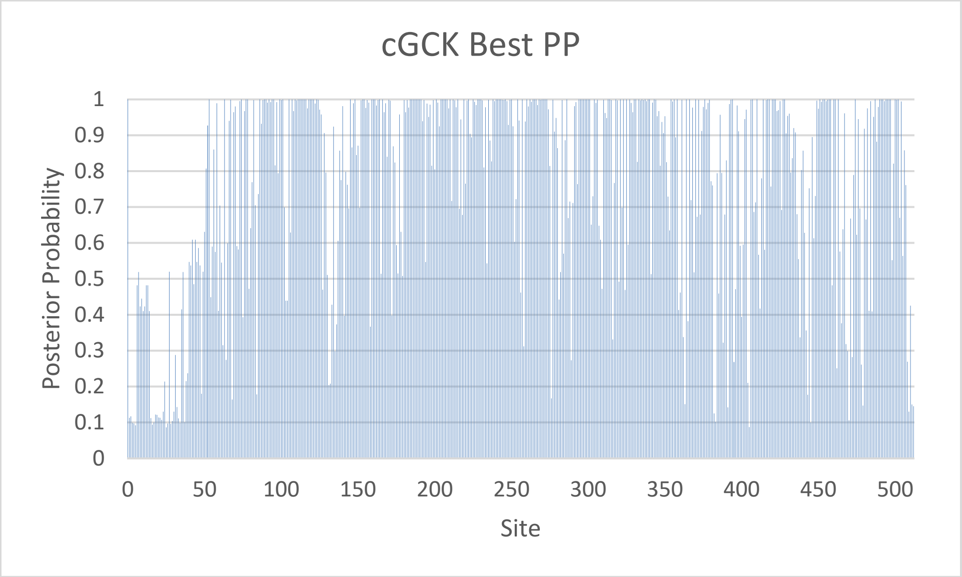

Fig. S18. Posterior probability across sites for the chordate GCK ancestor from the PhyloBot best tree reconstruction.

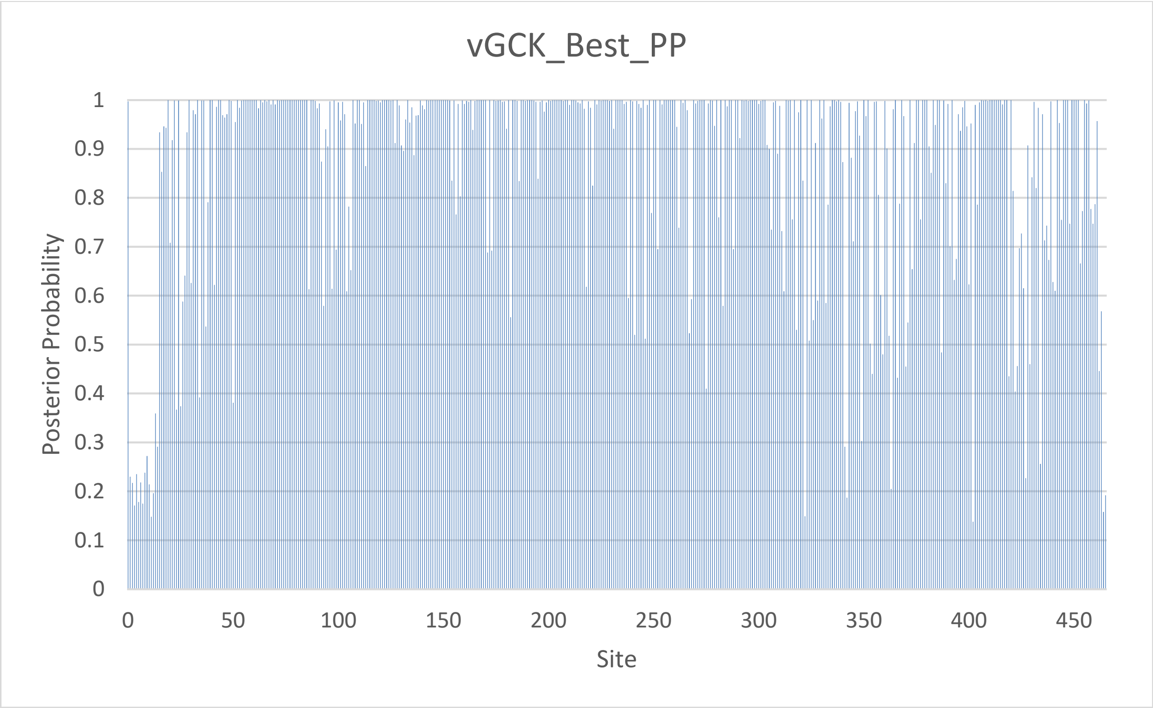

Fig. S19. Posterior probability across sites for the vertebrate GCK ancestor from the PhyloBot best tree reconstruction.

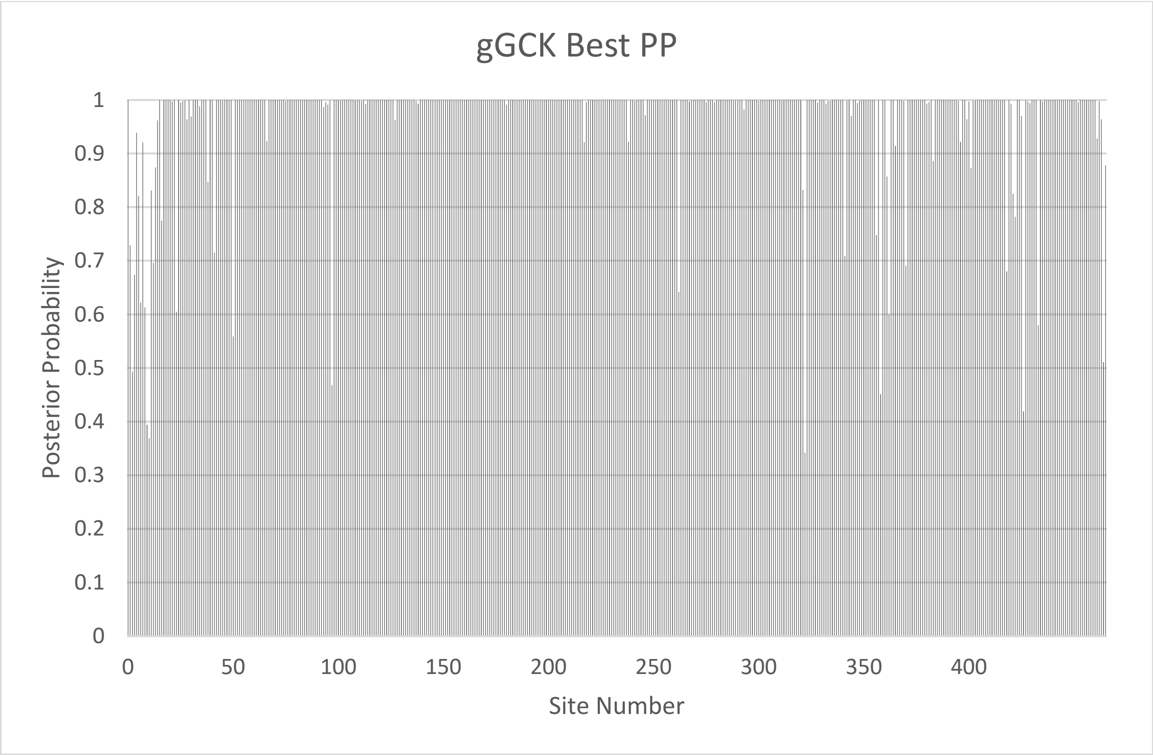

Fig. S20. Posterior probability across sites for the gnathostome GCK ancestor from the PhyloBot best tree reconstruction.

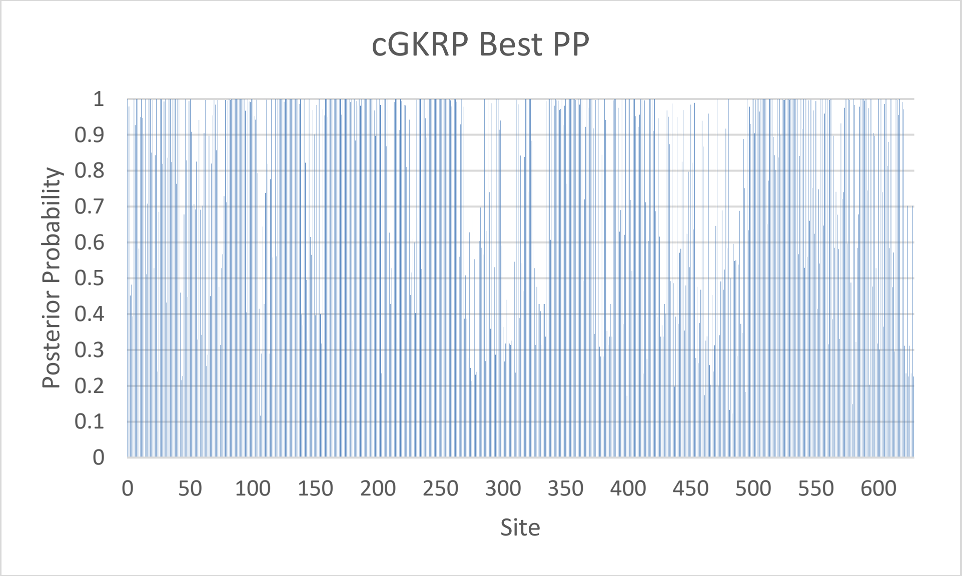

Fig. S21. Posterior probability across sites for the chordate GKRP ancestor from the PhyloBot best tree reconstruction.

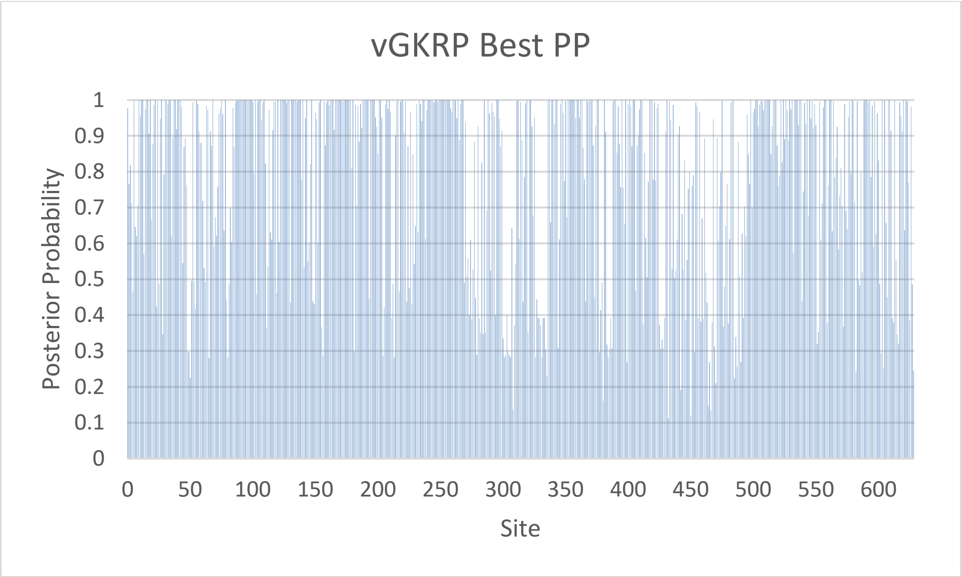

Fig. S22. Posterior probability across sites for the vertebrate GKRP ancestor from the PhyloBot best tree reconstruction.

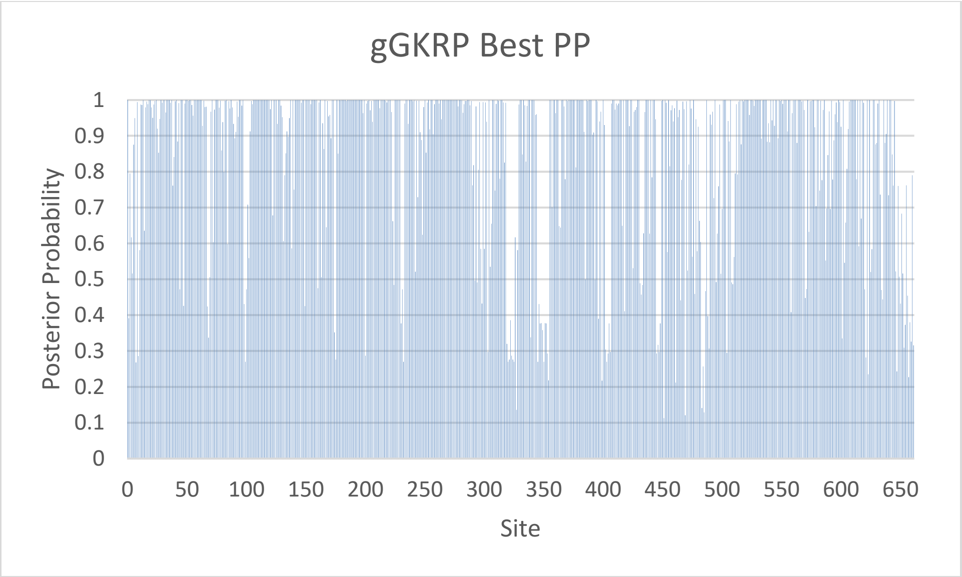

Fig. S23. Posterior probability across sites for the gnathostome GKRP ancestor from the PhyloBot best tree reconstruction.

Replicate 1

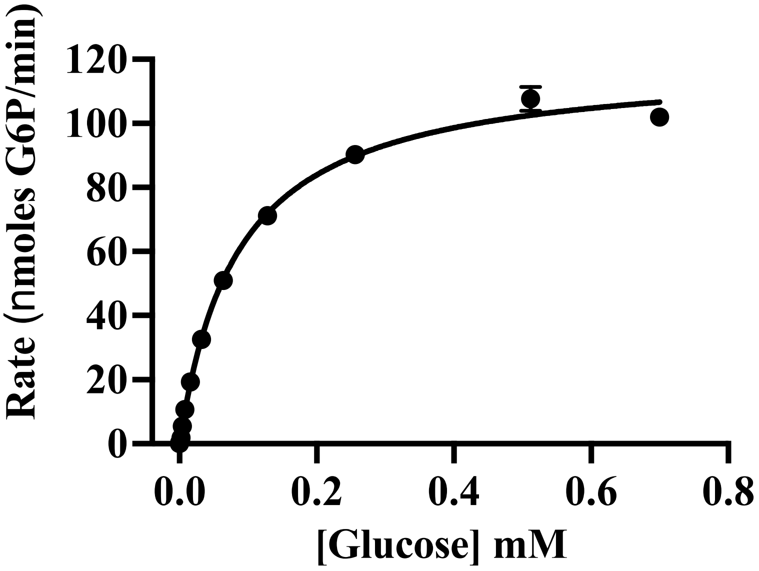

Replicate 2

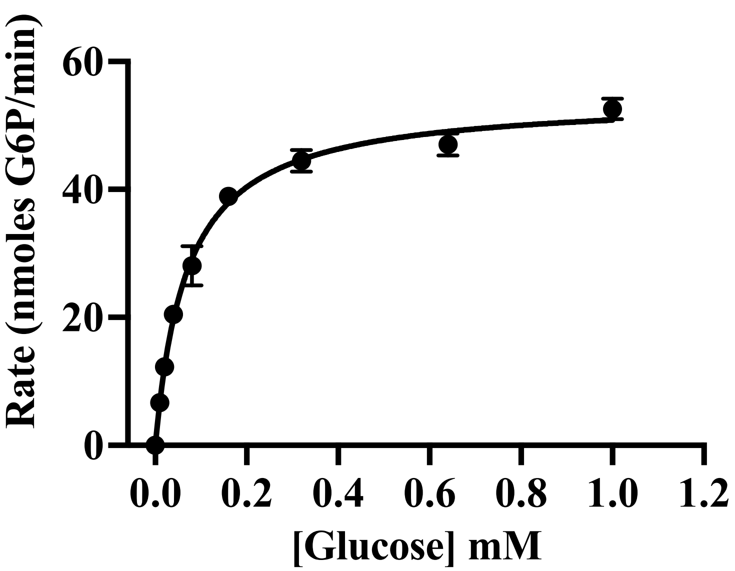

**Fig. S24**. Steady-state kinetic assays of ancestral chordate GCK (cGCK) with variable glucose concentrations.

Replicate 1

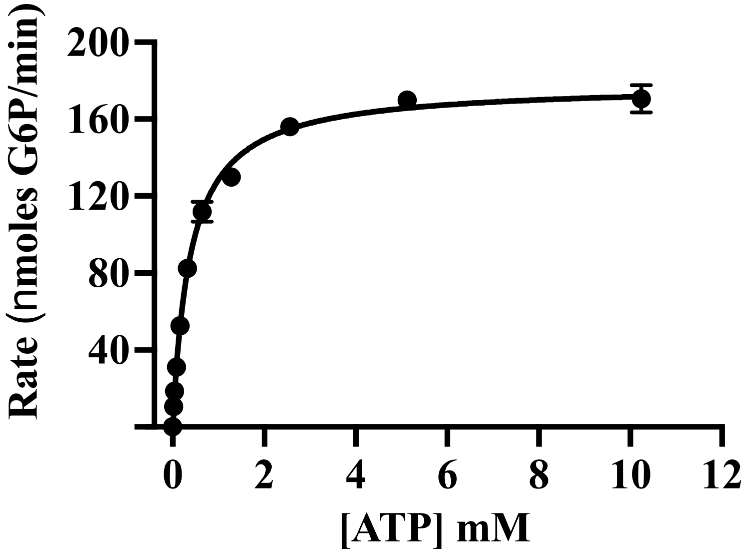

Replicate 2

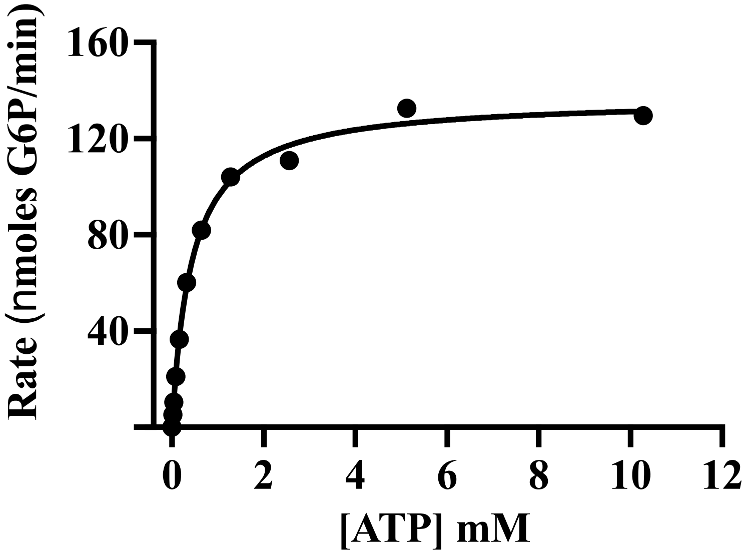

**Fig. S25**. Steady-state kinetic assays of cGCK with variable ATP concentrations.

Replicate 1

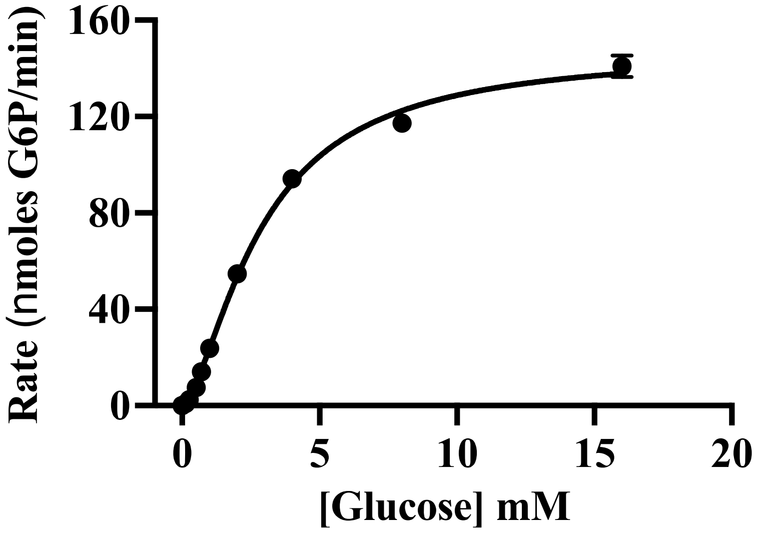

Replicate 2

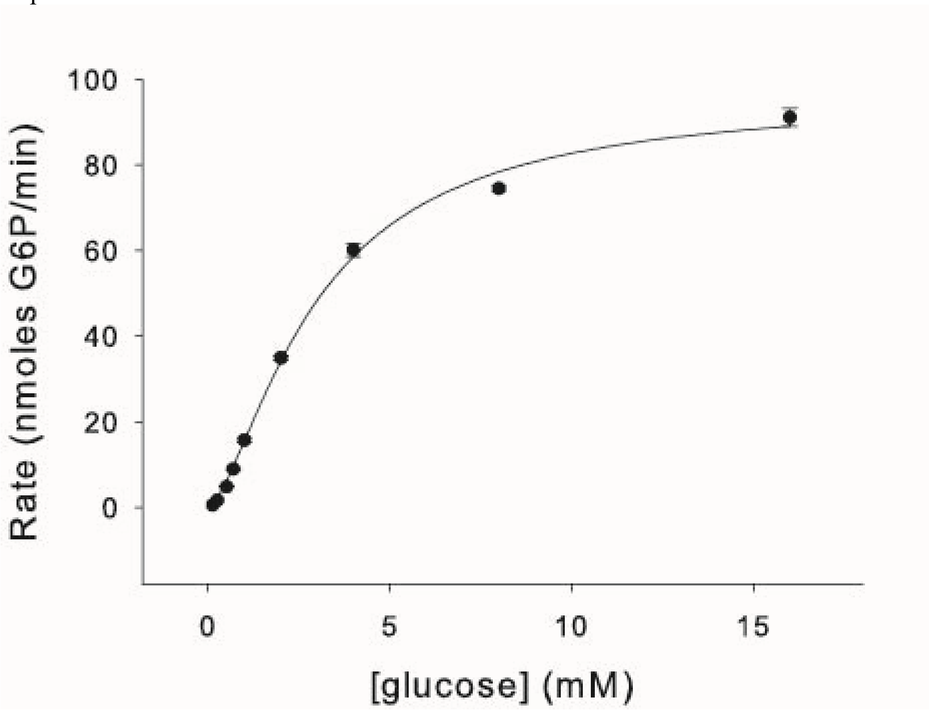

**Fig. S26**. Steady-state kinetic assays of ancestral vertebrate GCK (vGCK) with variable glucose concentrations.

Replicate 1

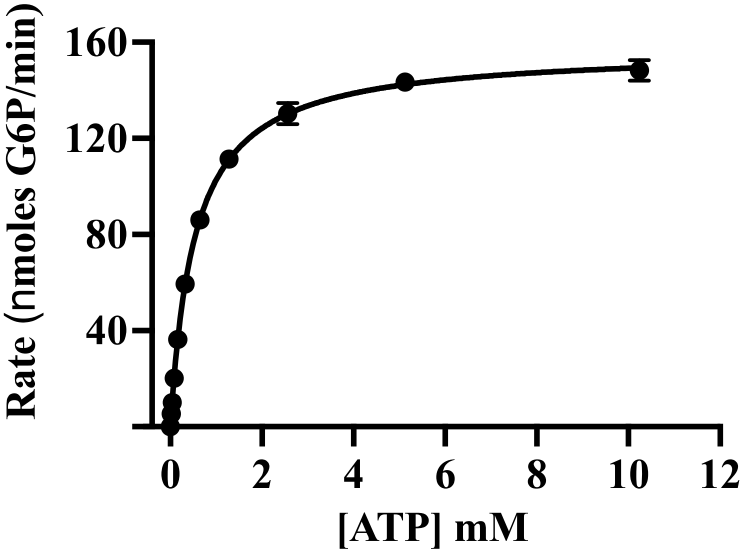

Replicate 2

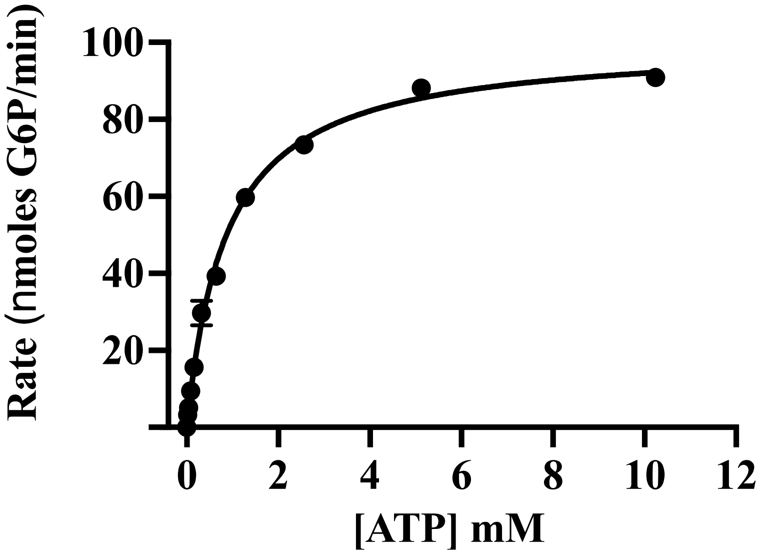

**Fig. S27**. Steady-state kinetic assays of ancestral vGCK with variable ATP concentrations.

Replicate 1

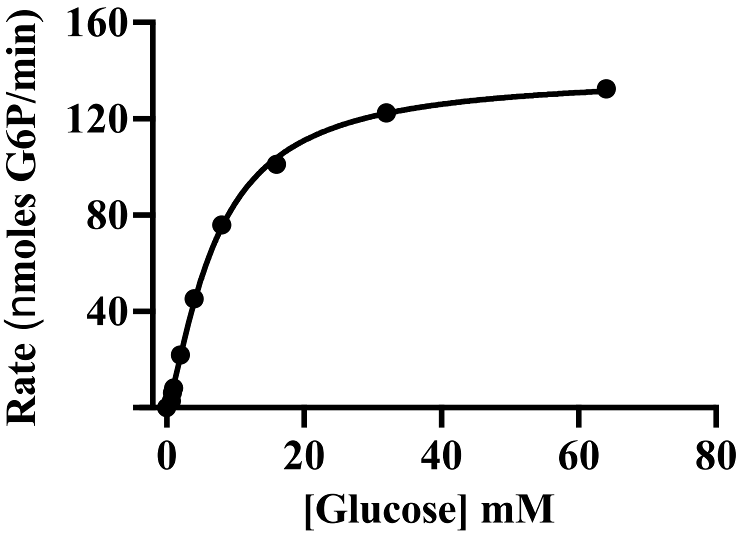

Replicate 2

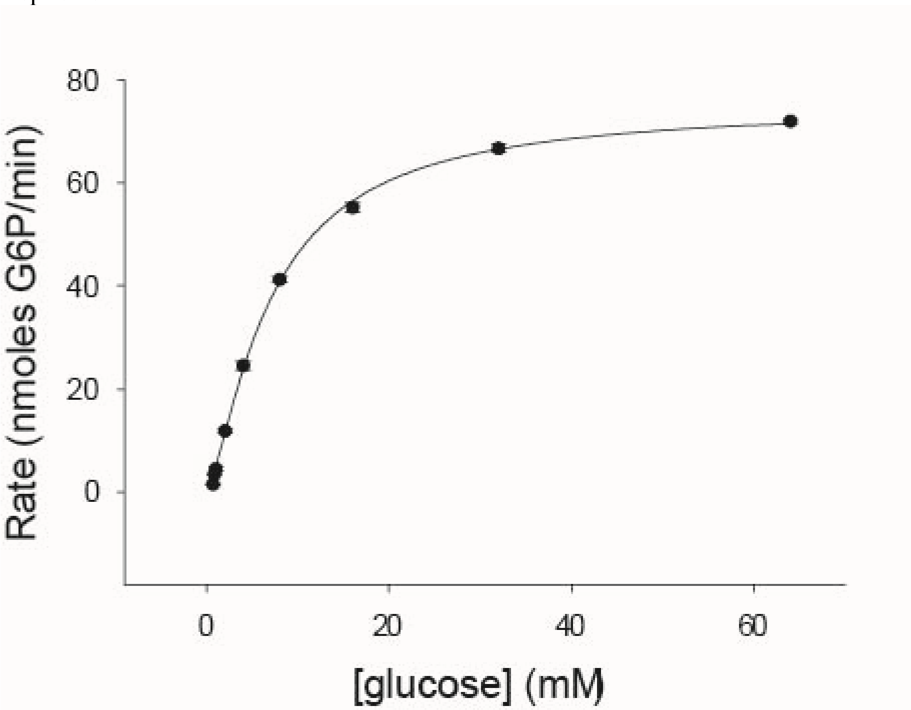

**Fig. S28**. Steady-state kinetic assays of ancestral gnathostome GCK (gGCK) with variable glucose concentrations.

Replicate 1

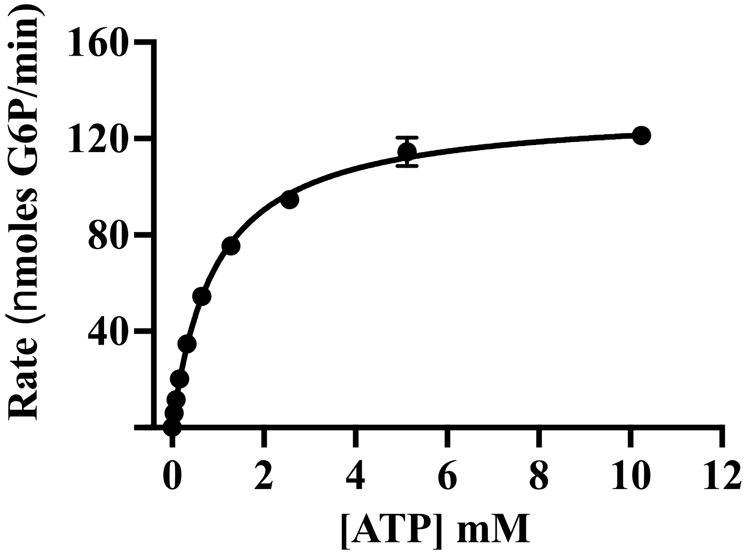

Replicate 2

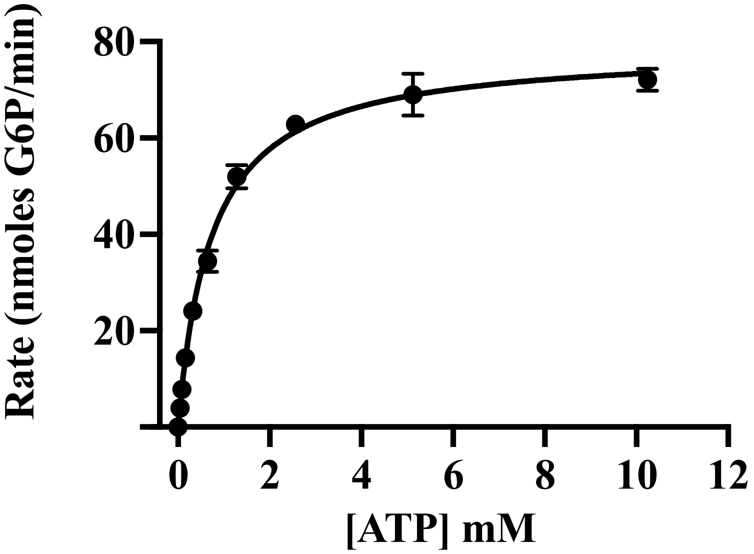

**Fig. S29**. Steady-state kinetic assays of ancestral gGCK with variable ATP concentrations.

Replicate 1

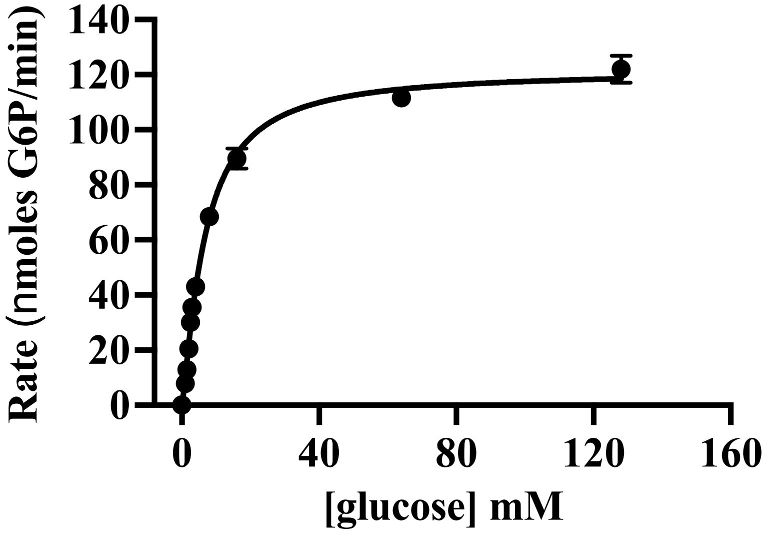

Replicate 2

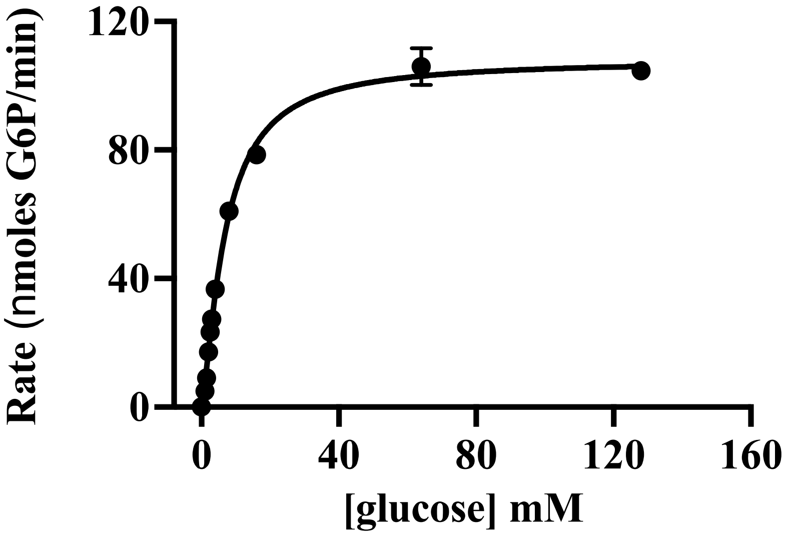

**Fig. S30**. Steady-state kinetic assays of ancestral tetrapod GCK (tGCK) with variable glucose concentrations.

Replicate 1

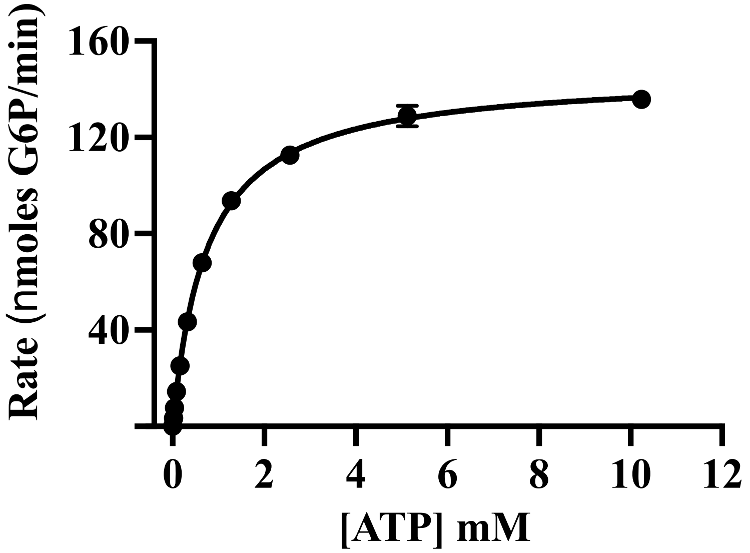

Replicate 2

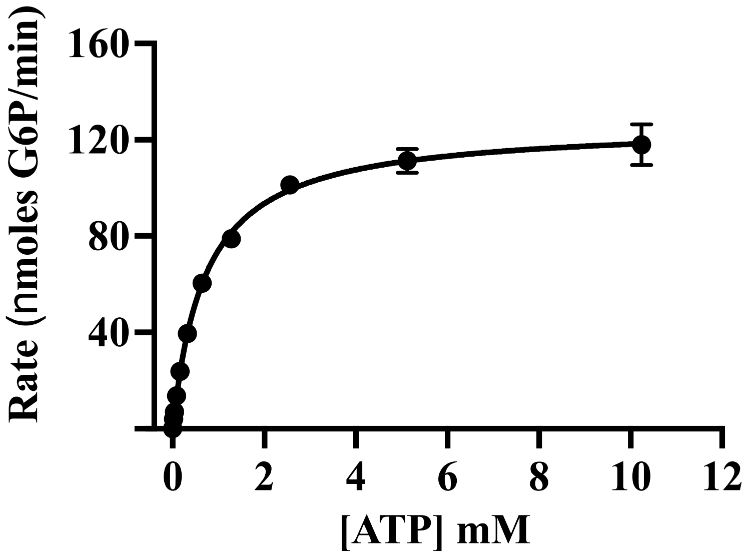

**Fig. S31**. Steady-state kinetic assays of ancestral tGCK with variable ATP concentrations.

Replicate 1

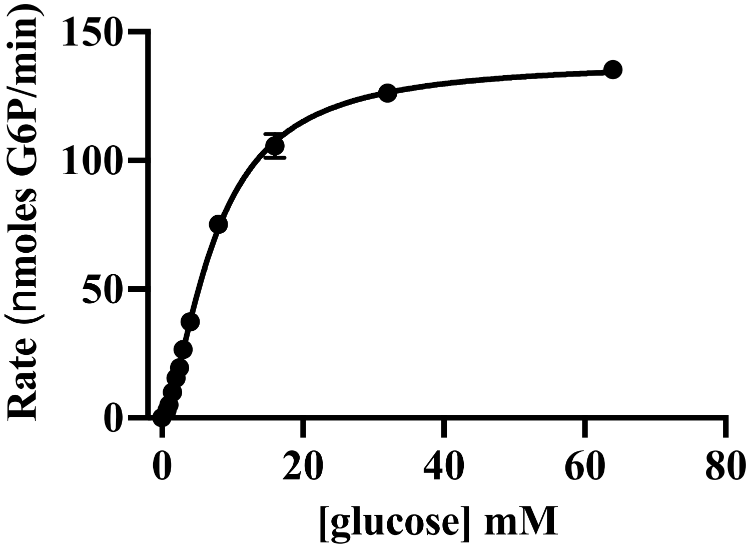

Replicate 2

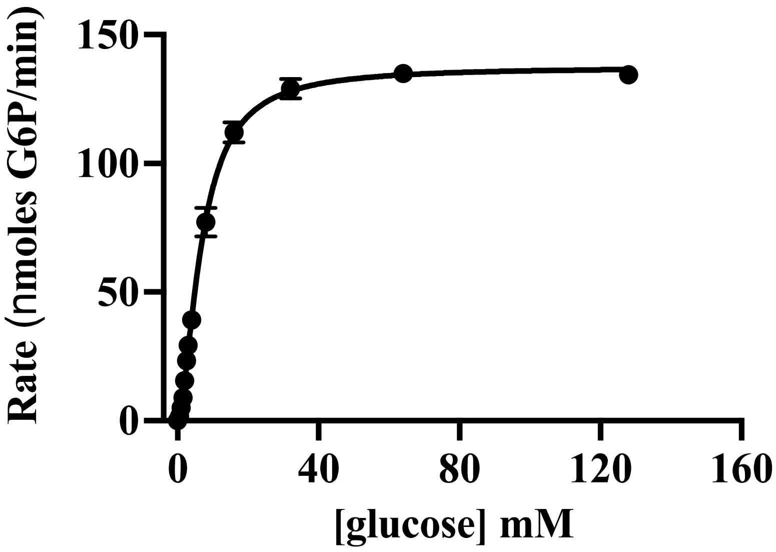

**Fig. S32**. Steady-state kinetic assays of ancestral mammal GCK (mGCK) with variable glucose concentrations.

**y**

Replicate 1

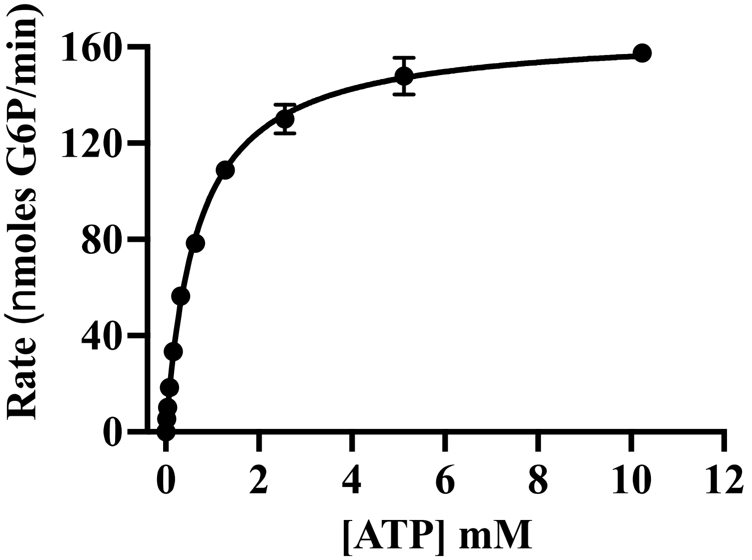

Replicate 2

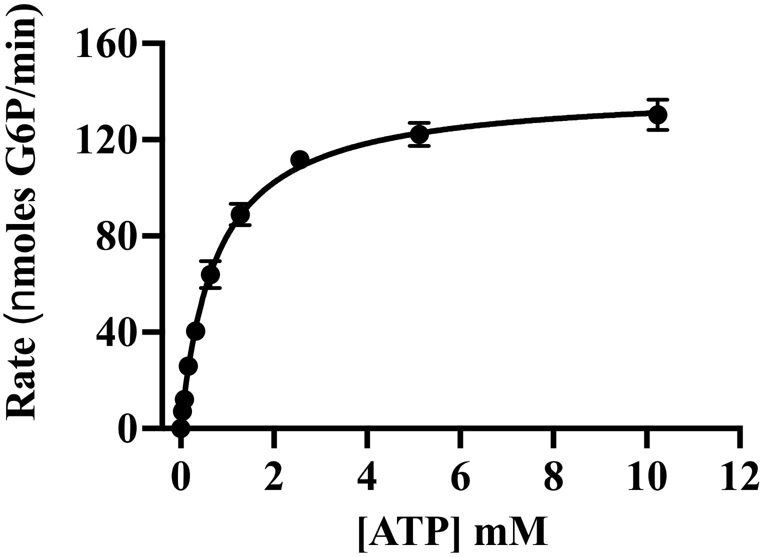

**Fig. S33**. Steady-state kinetic assays of ancestral mGCK with variable ATP concentrations.

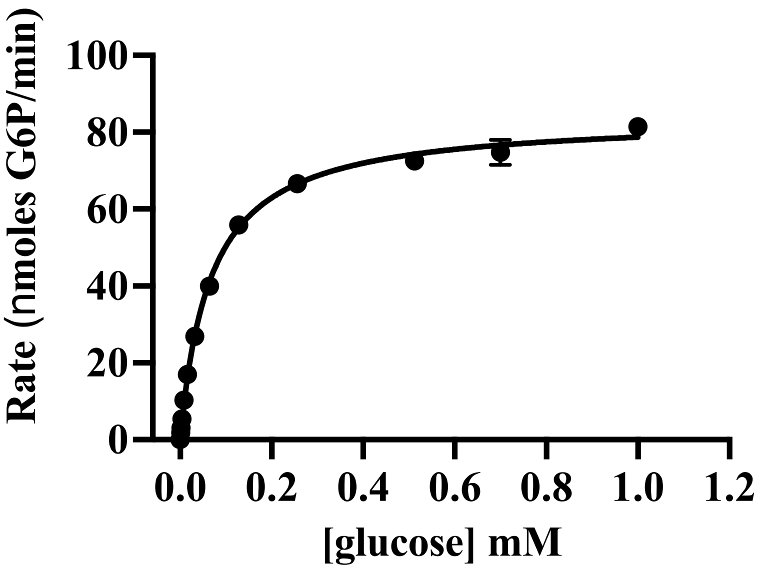

**Fig. S34**. Steady state measurements of the glucokinase reaction rates at varying glucose concentrations for cGCK Worst. Each data point represents the average of triplicate measurements of the highest rate of glucose 6-phosphate production observed over at least one minute of data collection.

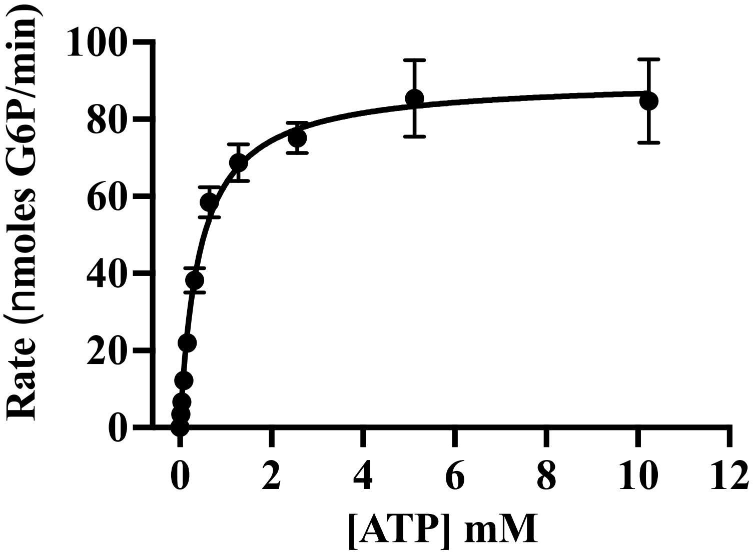

**Fig. S35**. Steady state measurements of the glucokinase reaction rates at varying ATP concentrations for cGCK Worst. Each data point represents the average of triplicate measurements of the highest rate of glucose 6-phosphate production observed over at least one minute of data collection.

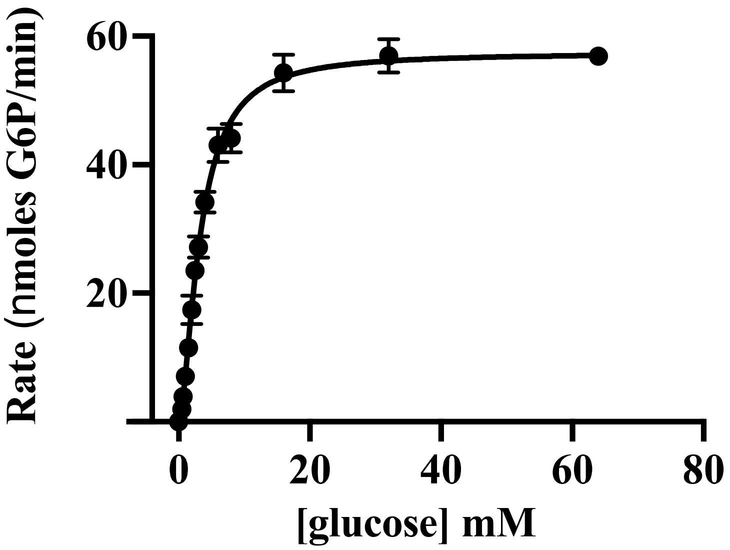

**Fig. S36**. Steady state measurements of the glucokinase reaction rates at varying glucose concentrations for vGCK Worst. Each data point represents the average of triplicate measurements of the highest rate of glucose 6-phosphate production observed over at least one minute of data collection.

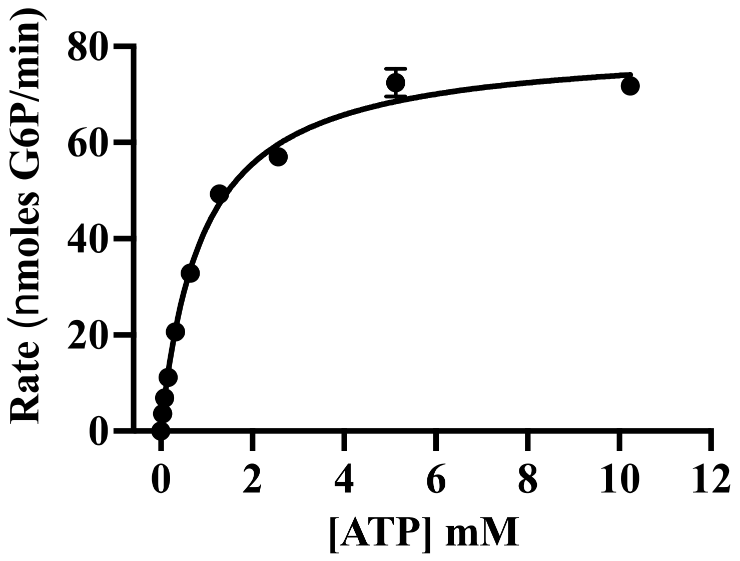

**Fig. S37**. Steady state measurements of the glucokinase reaction rates at varying ATP concentrations for vGCK Worst. Each data point represents the average of triplicate measurements of the highest rate of glucose 6-phosphate production observed over at least one minute of data collection.

Replicate 1

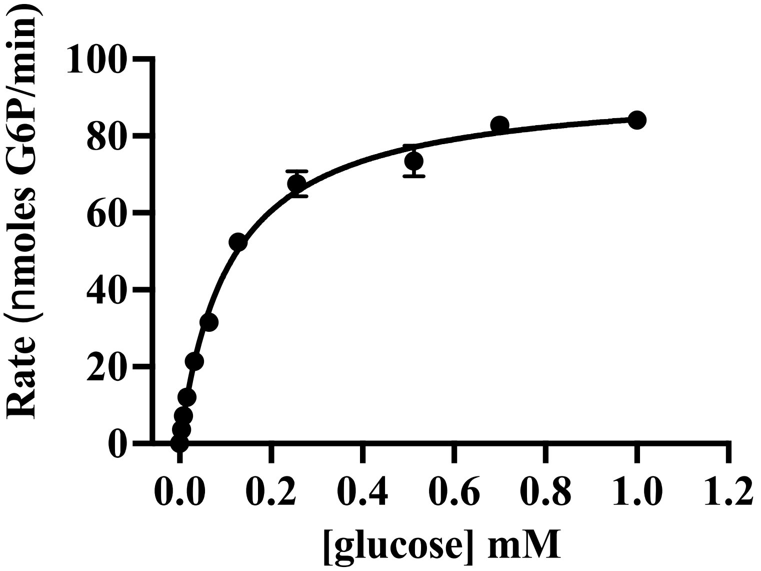

Replicate 2

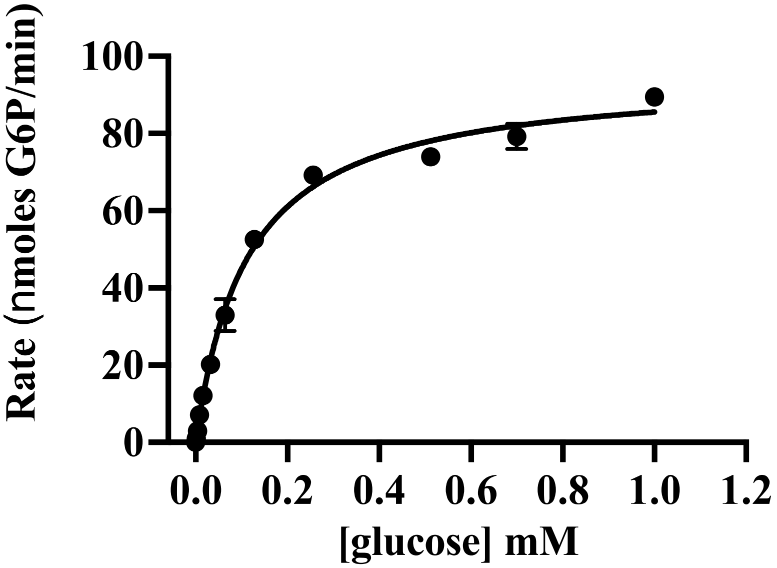

**Fig. S38**. Steady state measurements of the glucokinase reaction rates at varying glucose concentrations for cGCK AltAll. Each data point represents the average of triplicate measurements of the highest rate of glucose 6-phosphate production observed over at least one minute of data collection.

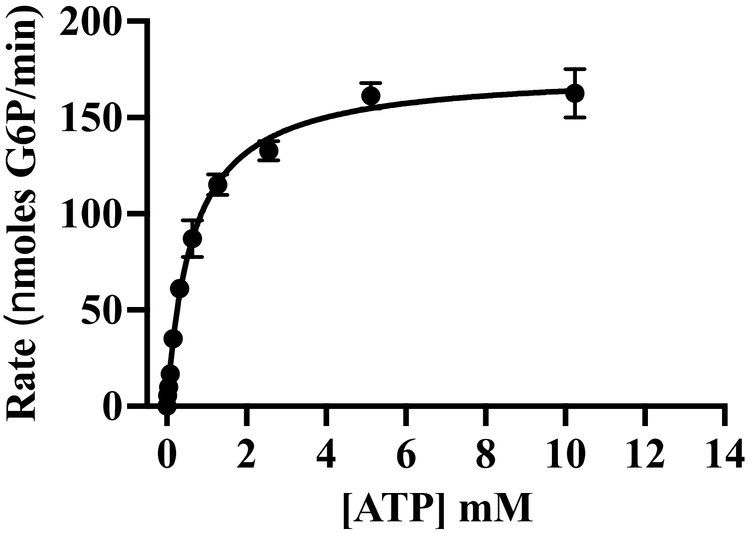

**Fig. S39**. Steady state measurements of the glucokinase reaction rates at varying ATP concentrations for cGCK AltAll. Each data point represents the average of triplicate measurements of the highest rate of glucose 6-phosphate production observed over at least one minute of data collection.

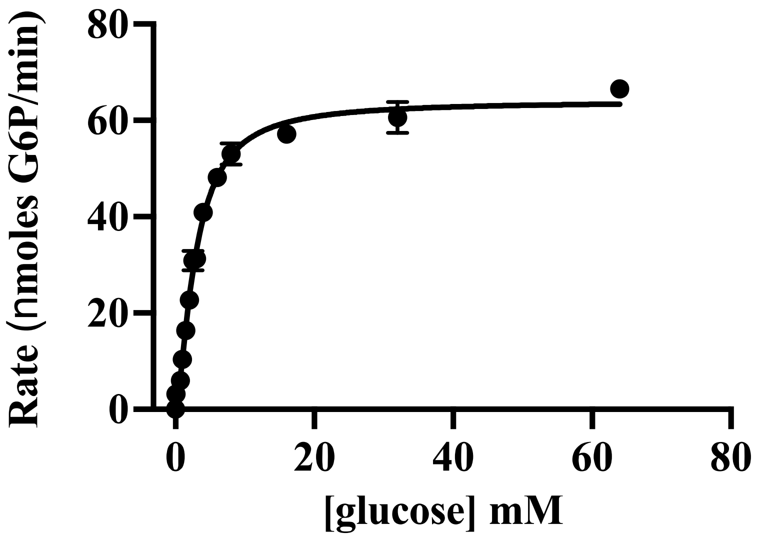

**Fig. S40**. Steady state measurements of the glucokinase reaction rates at varying glucose concentrations for vGCK AltAll. Each data point represents the average of triplicate measurements of the highest rate of glucose 6-phosphate production observed over at least one minute of data collection.

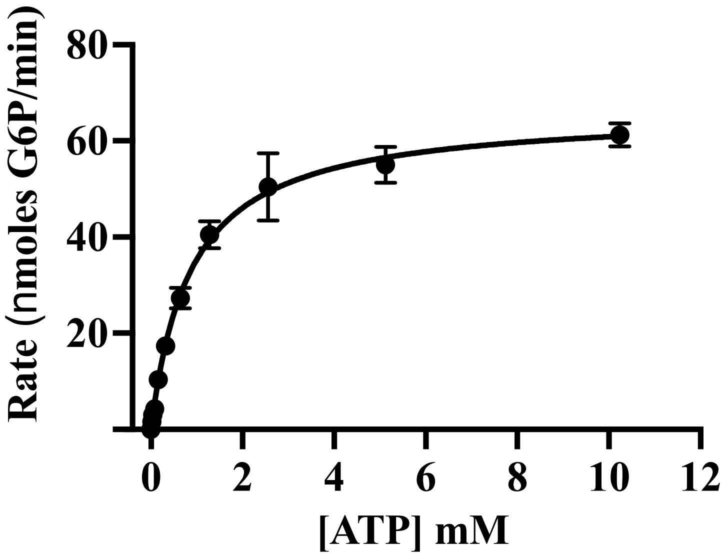

**Fig. S41**. Steady state measurements of the glucokinase reaction rates at varying ATP concentrations for vGCK AltAll. Each data point represents the average of triplicate measurements of the highest rate of glucose 6-phosphate production observed over at least one minute of data collection.

Replicate 1

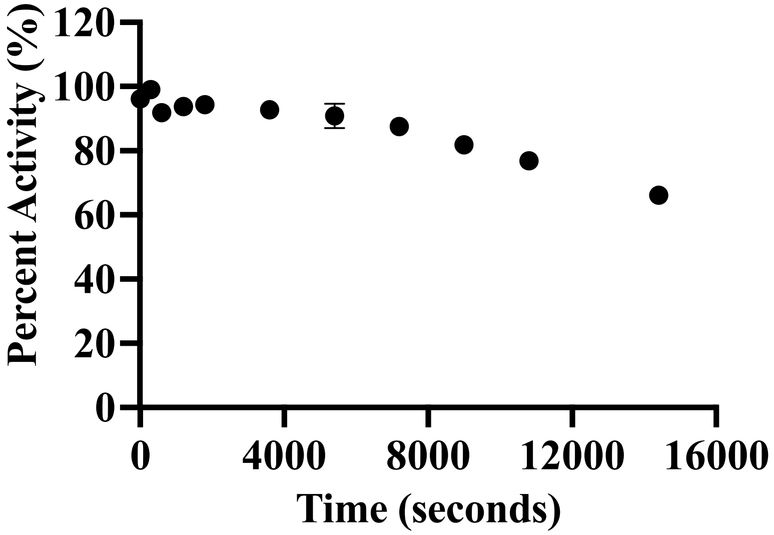

Replicate 2

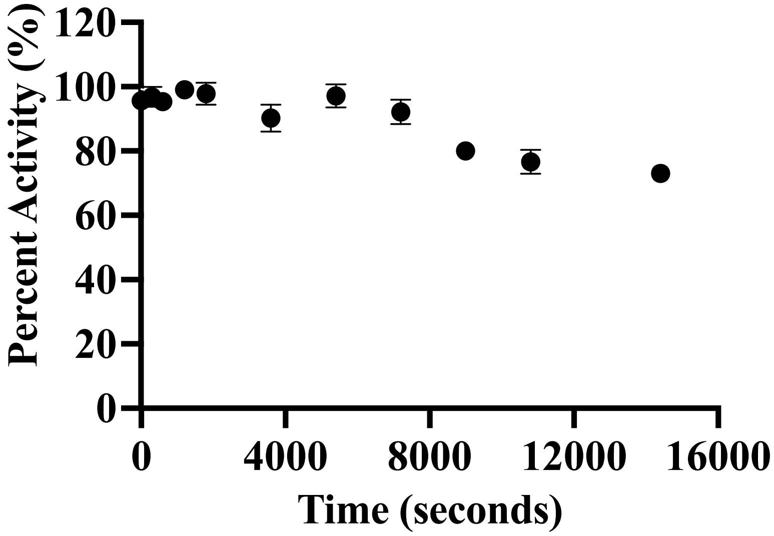

**Fig. S42**. Results of limited proteolysis experiments on *C. intestinalis* GCK.

Replicate 1

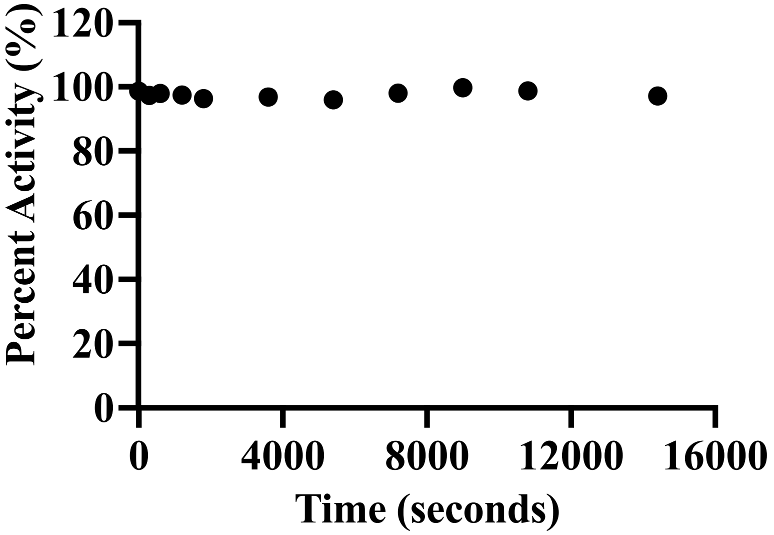

Replicate 2

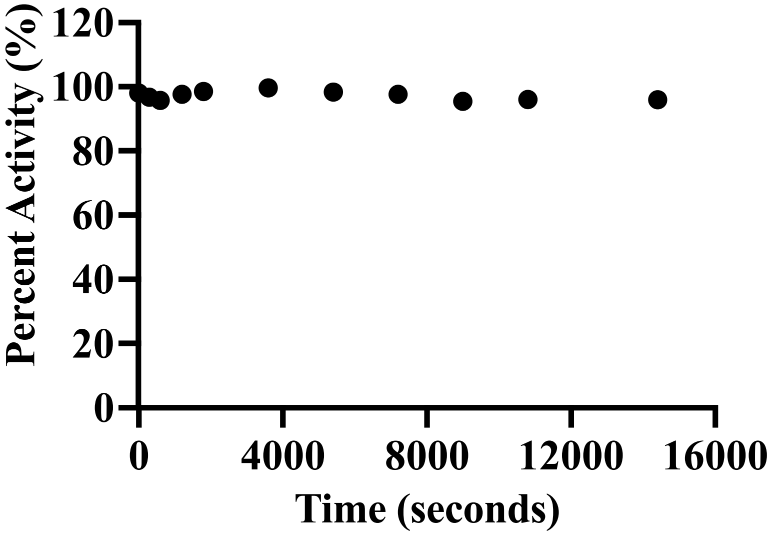

**Fig. S43**. Results of limited proteolysis experiments on *B. belcheri* GCK.

Replicate 1

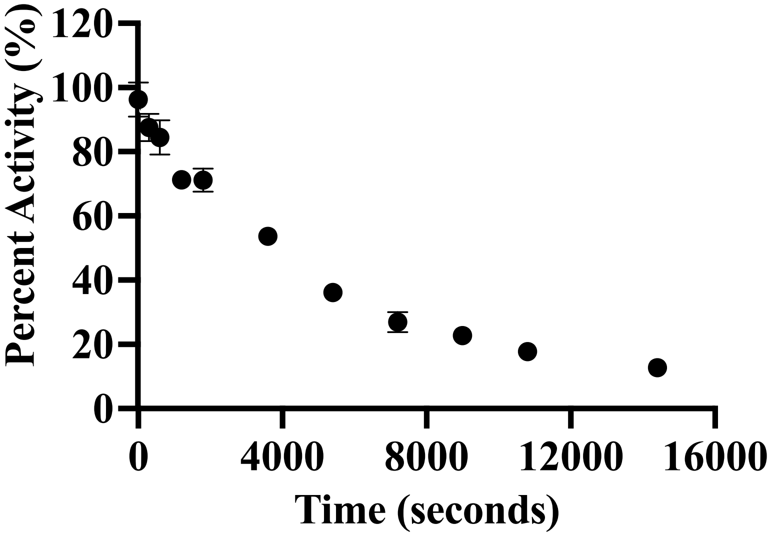

Replicate 2

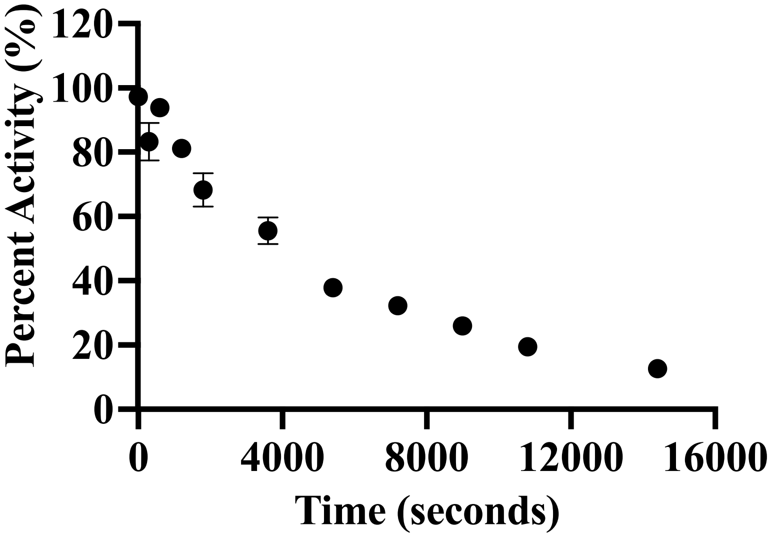

**Fig. S44**. Results of limited proteolysis experiments on *D. rerio* GCK.

Replicate 1

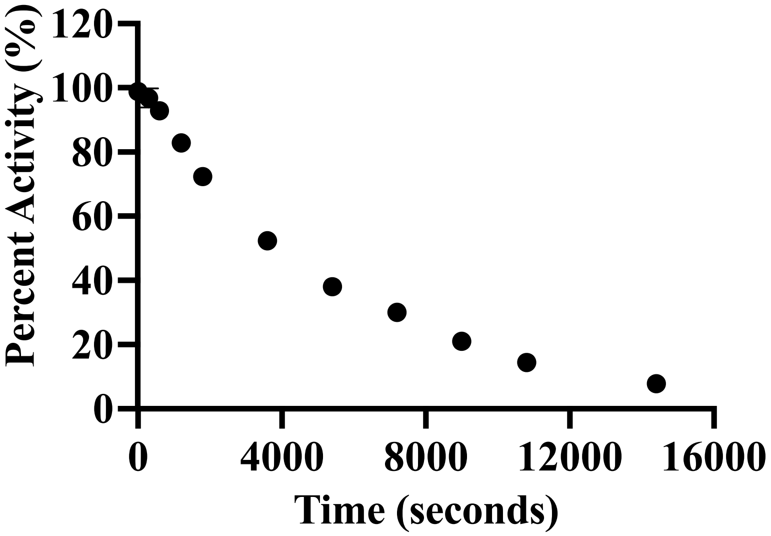

Replicate 2

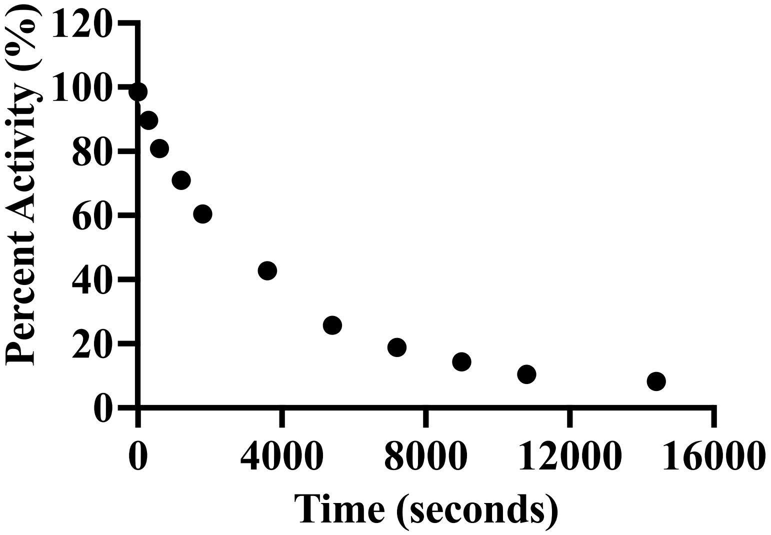

**Fig. S45**. Results of limited proteolysis experiments on *X. laevis* GCK.

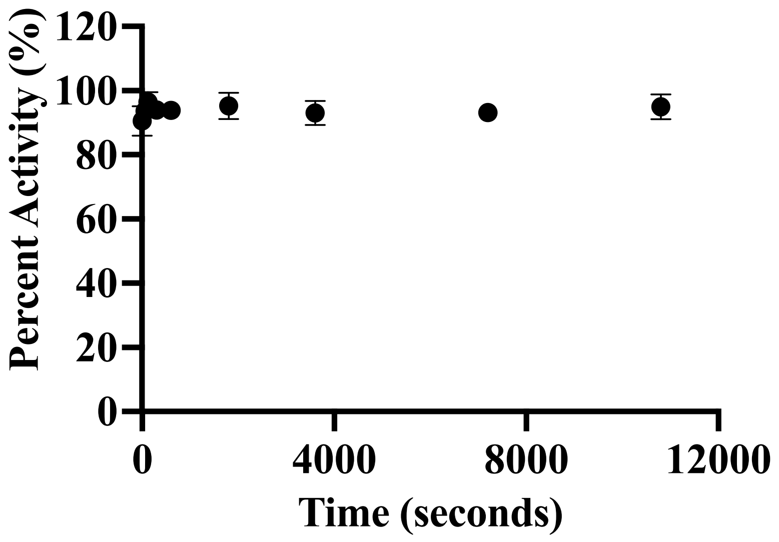

**Fig. S46**. Limited proteolysis of cGCK

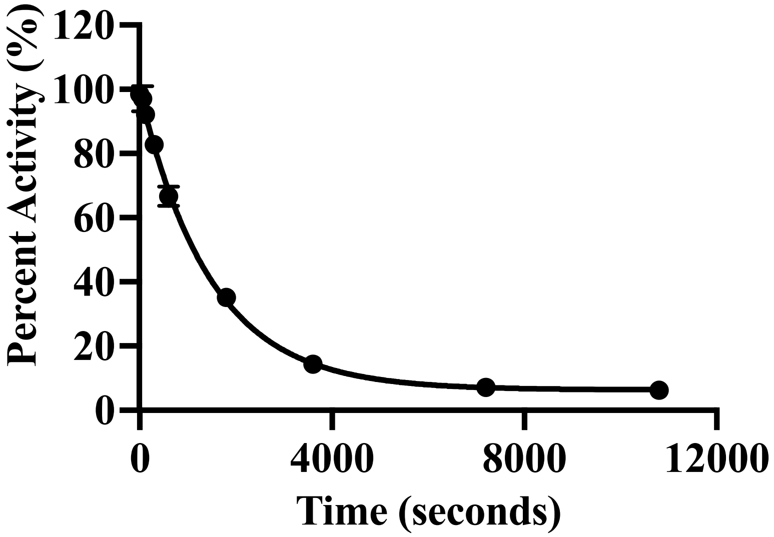

**Fig. S47**. Limited proteolysis of vGCK

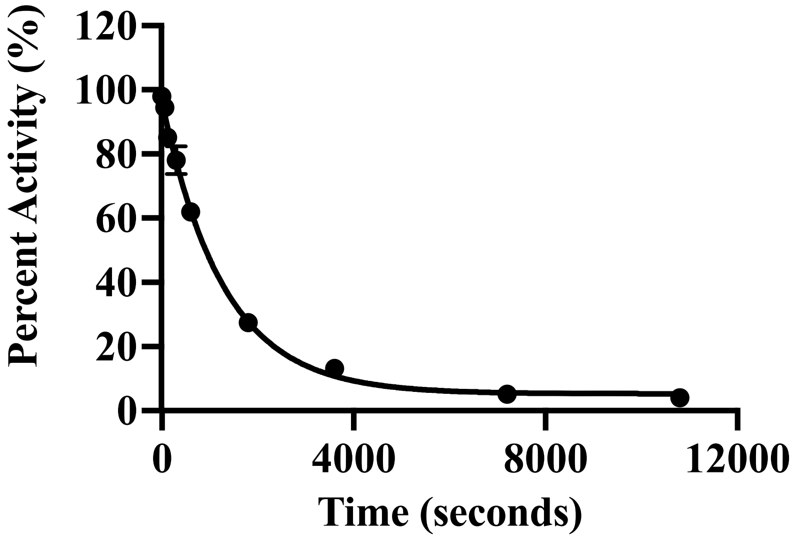

**Fig. S48**. Limited proteolysis of gGCK

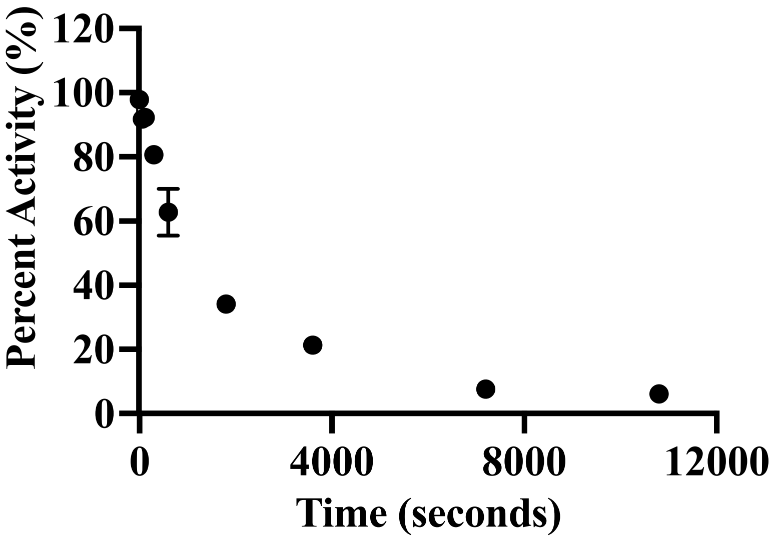

**Fig. S49**. Limited proteolysis of tGCK

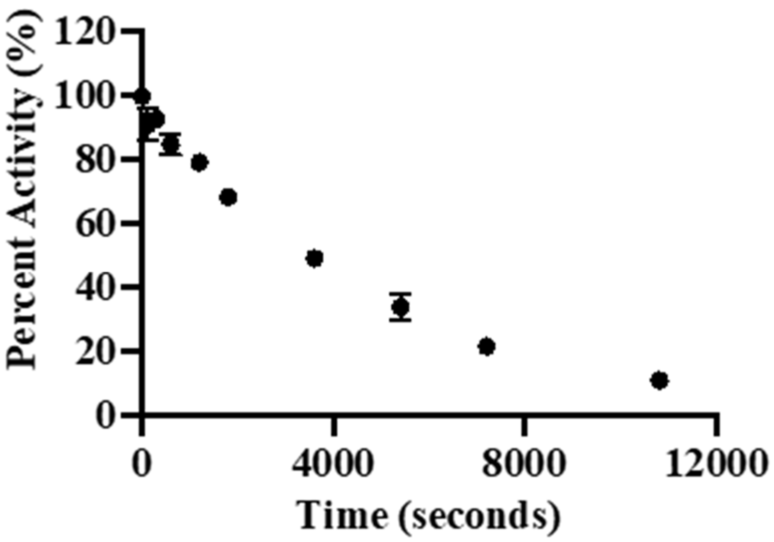

Fig. S50. Limited proteolysis of mGCK

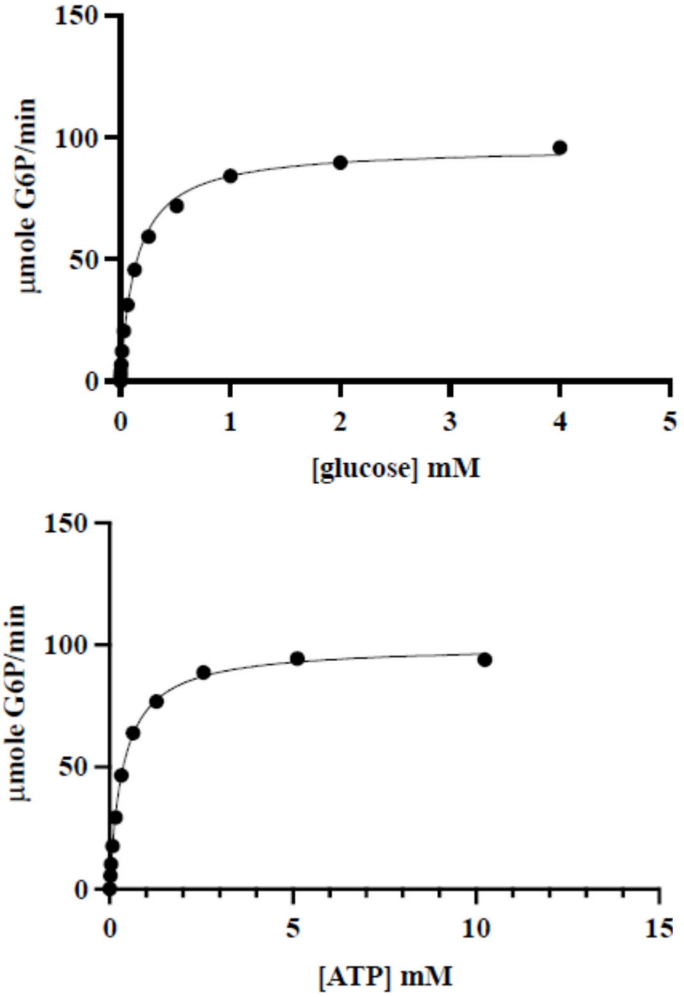

**Fig. S51**. Steady-state glucose (top) and ATP (bottom) kinetics of a variant with the disordered loop of vGCK installed into the cGCK background.

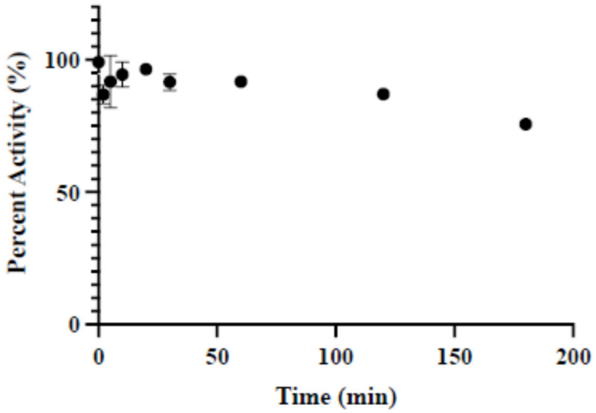

**Fig. S52**. Limited proteolysis of a variant with the disordered loop of vGCK installed into the cGCK background.

```
cGCK   1 MHHHHHHGSGSMALRE--------EKVELILDEFHLDNEELNEIMGRMHKEMEKGLRKE  51
vGCK   1 MHHHHHHGSGSMLGRRSRMEGRKGEKVEQILSEFRLDKEELEEVMRRMQREMERGLRLE  59

cGCK  52 TNEDATVKMLPTYVRSLPDGTESGDFLALDLGGTNFRVLLVKIKEGEELEGERKVEMKS 110
vGCK  60 THEEASVKMLPTYVRSTPDGSEVGDFLALDLGGTNFRVMLVKVGEDE--EGEWKVETKN 116

cGCK 111 QIYRIPEDVMTGTGEQLFDYIAECMADFLEKLGMKDRKLPLGFTFSFPCKQDGLDSASL 169
vGCK 117 QMYCIPEDVMTGTAEMLFDYIAECIADFLDKLNMKHKKLPLGFTFSFPVKHEDLDKGIL 175

cGCK 170 ITWTKGFSATGVEGKDVVKLLRDAIKRRGDFDMDIVAVVNDTVGTMMSCAFEDHDCLIG 228
vGCK 176 INWTKGFTATGAEGNNVVELLRDAIKRRGDFDMDVVAMVNDTVATMISCYYEDHNCEIG 234

cGCK 229 LIVGTGSNACYMEKMENVELLEGDKGEPNQMCINMEWGAFGDDGALDDFRTEYDREVDE 287
vGCK 235 MIVGTGCNACYMEEMRNVELVEGEEG---RMCINMEWGAFGDSGELEEFRLEYDRKVDE 290

cGCK 288 NSLNPGQQLYEKMISGMYMGELVRLVLLKLTKEGLLFGGKTSEELKTPGTFQTKYVSQI 346
vGCK 291 TSLNPGQQLYEKIISGKYMGELVRLVLLKLTNEGLLFGGKASEKLKTRGSFETKYVSQI 349

cGCK 347 ESDVPGDMTATLNILASLGLRHATEVDCEIVRQVCRAVSTRAAHLCAAGIAAVVNKMRR 405
vGCK 350 ESDDSGDMKQTYNILTTLGLQHPTELDCEIVRRVCQAVSTRAAHLCAAGMAAVVNKMRE 408

cGCK 406 NR------ITVGVDGSVYKYHPTFKELMSETVDELTPGCDVKFMLSEDGSGKGAALITA 458
vGCK 409 NRSQETLKITVGVDGSVYKLHPSFKDKFHAMVRELTPRCDITFIQSEEGSGRGAALISA 467

cGCK 459 VACRLAGK--                                                  466
vGCK 468 VACKMACMGQ                                                  477
```

**Fig. S53.** Map of common, overlapping peptide fragments between cGCK and vGCK used in hydrogen-deuterium exchange mass spectrometry analysis.

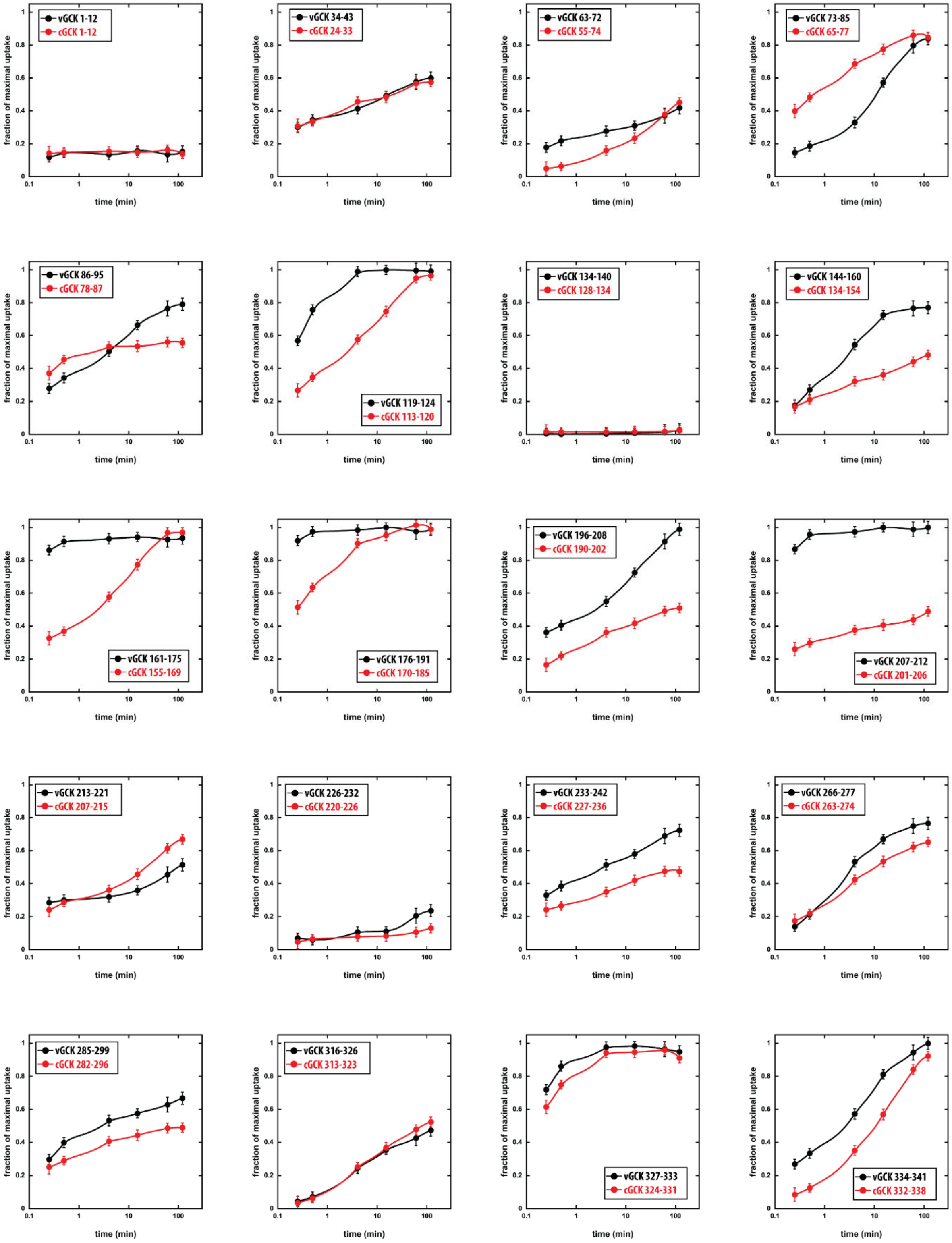
vGCK 1-12
cGCK 1-12
vGCK 34-43
cGCK 24-33
vGCK 63-72
cGCK 55-74
vGCK 73-85
cGCK 65-77
vGCK 86-95
cGCK 78-87
vGCK 119-124
cGCK 113-120
vGCK 134-140
cGCK 128-134
vGCK 144-160
cGCK 134-154
vGCK 161-175
cGCK 155-169
vGCK 176-191
cGCK 170-185
vGCK 196-208
cGCK 190-202
vGCK 207-212
cGCK 201-206
vGCK 213-221
cGCK 207-215
vGCK 226-232
cGCK 220-226
vGCK 233-242
cGCK 227-236
vGCK 266-277
cGCK 263-274
vGCK 285-299
cGCK 282-296
vGCK 316-326
cGCK 313-323
vGCK 327-333
cGCK 324-331
vGCK 334-341
cGCK 332-338
fraction of maximal uptake
time (min)

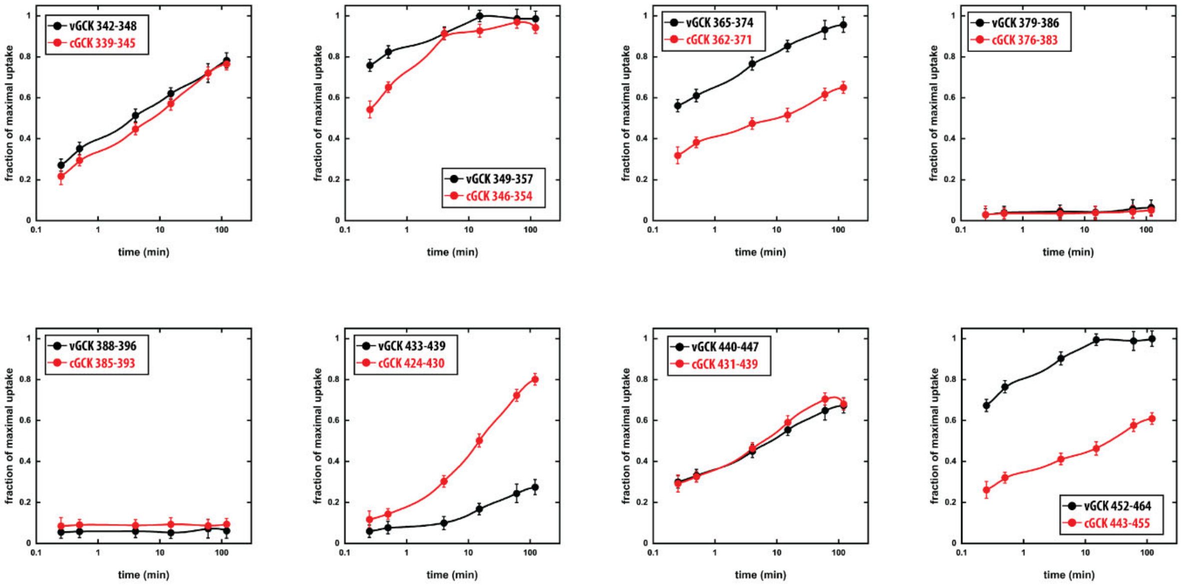

**Fig. S54**. Individual plots of fractional deuterium update as a function of time for 28 individual common peptides from cGCK (red) and vGCK (black). Data points represent the average of two independent time course experiments.

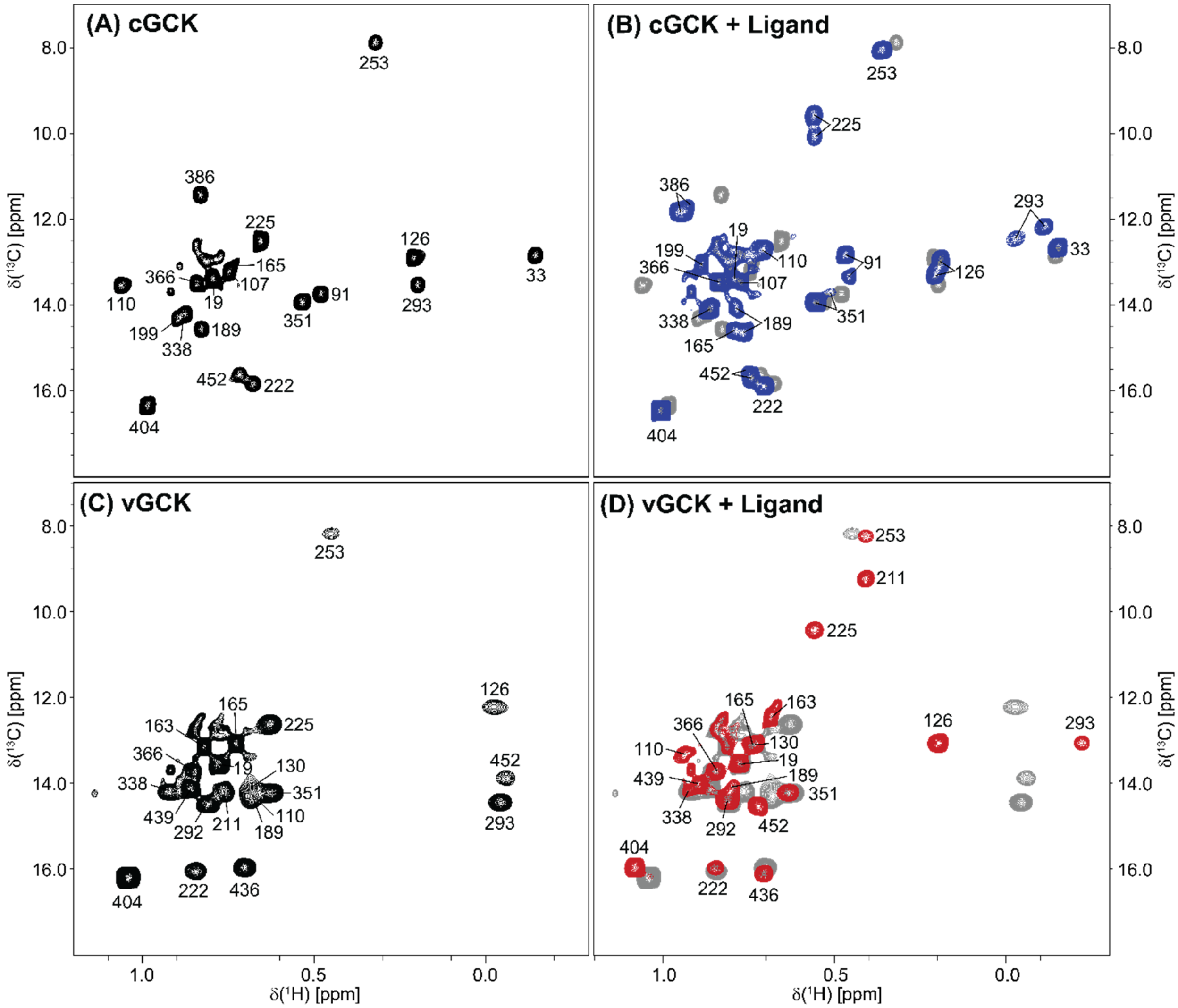

**Fig. S55**. Resonance assignment of isoleucine methyl groups in cGCK and vGCK. $^{1}$H-$^{13}$C ALSOFAST-HMQC spectra of (**A**) unliganded and (**B**) liganded cGCK are shown in black and blue, respectively. The spectrum of the unliganded cGCK is displayed in gray in panel (B) for reference. $^{1}$H-$^{13}$C ALSOFAST-HMQC spectra of (**C**) unliganded and (**D**) liganded vGCK are shown in black and red, respectively. The spectrum of the unliganded vGCK is displayed in gray in panel (B) for reference. Numbering is based on human GCK (GenBank: AAA52562.1; see SI alignment files for corresponding numbering in cGCK and vGCK).

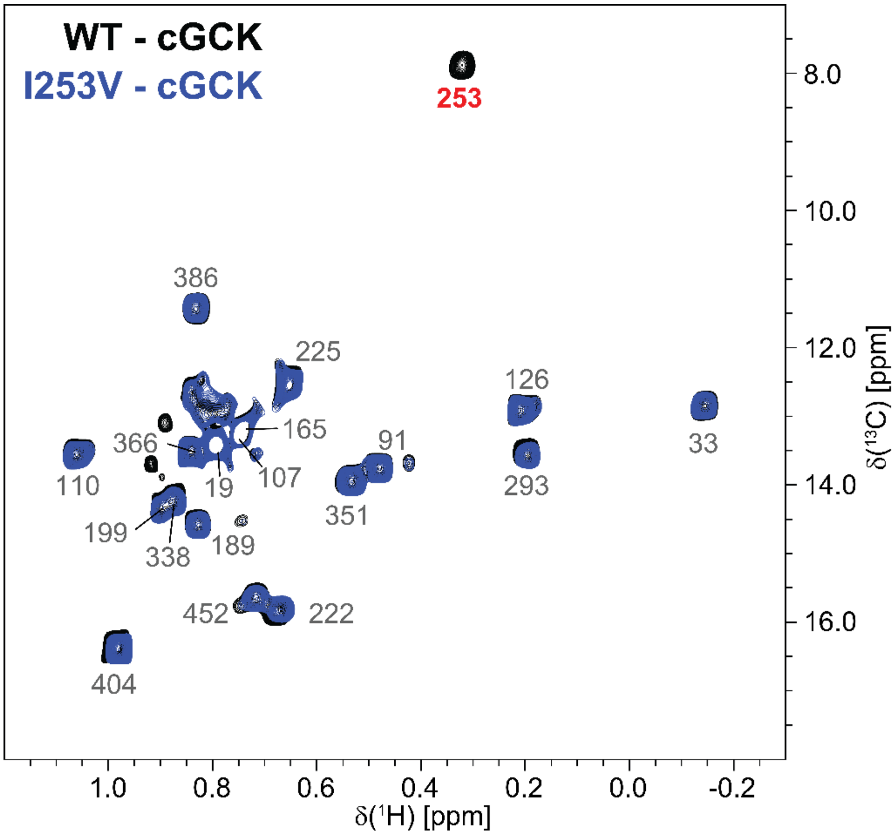

**Fig. S56**. Assignment strategy of isoleucine methyl groups (Case A). $^{1}$H-$^{13}$C ALSOFAST-HMQC spectra of unliganded wild type cGCK and unliganded cGCK-I253V are shown in black and blue, respectively. For case A, the isoleucine-to-valine mutation leads to a single peak missing (residue 253, shown in bold red), while none of the other peaks display changes in intensity or peak position. Resonance assignments without changes are shown in gray for reference. Numbering is based on human GCK (GenBank: AAA52562.1; see SI alignment files for corresponding numbering in cGCK and vGCK).

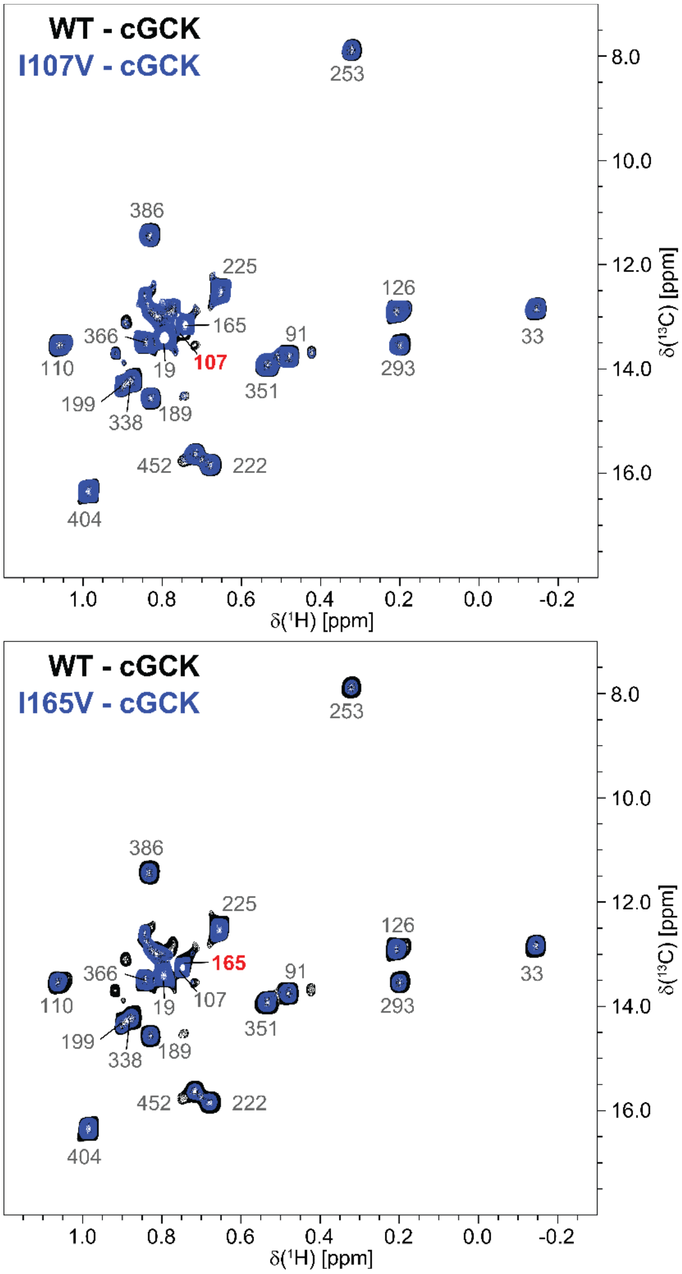

**Fig. S57**. Assignment strategy of isoleucine methyl groups (Case B). (TOP) $^{1}H$-$^{13}C$ ALSOFAST-HMQC spectra of unliganded wild type cGCK and unliganded cGCK-I107V are shown in black and blue, respectively. (BOTTOM) $^{1}H$-$^{13}C$ ALSOFAST-HMQC spectra of unliganded wild type cGCK and unliganded cGCK-I165V are shown in black and blue, respectively. For case B, the isoleucine-to-valine mutation leads to a single peak missing (bold red) similar to case A, but peaks overlap. Here, residues 107 and 165 in the top and bottom spectra, respectively, are missing, while none of the other peaks display changes in intensity or peak position. Peak intensity was evaluated and confirmed using 1D projections where necessary. Resonance assignments without changes are shown in gray for reference. Numbering is based on human GCK (GenBank: AAA52562.1; see SI alignment files for corresponding numbering in cGCK and vGCK).

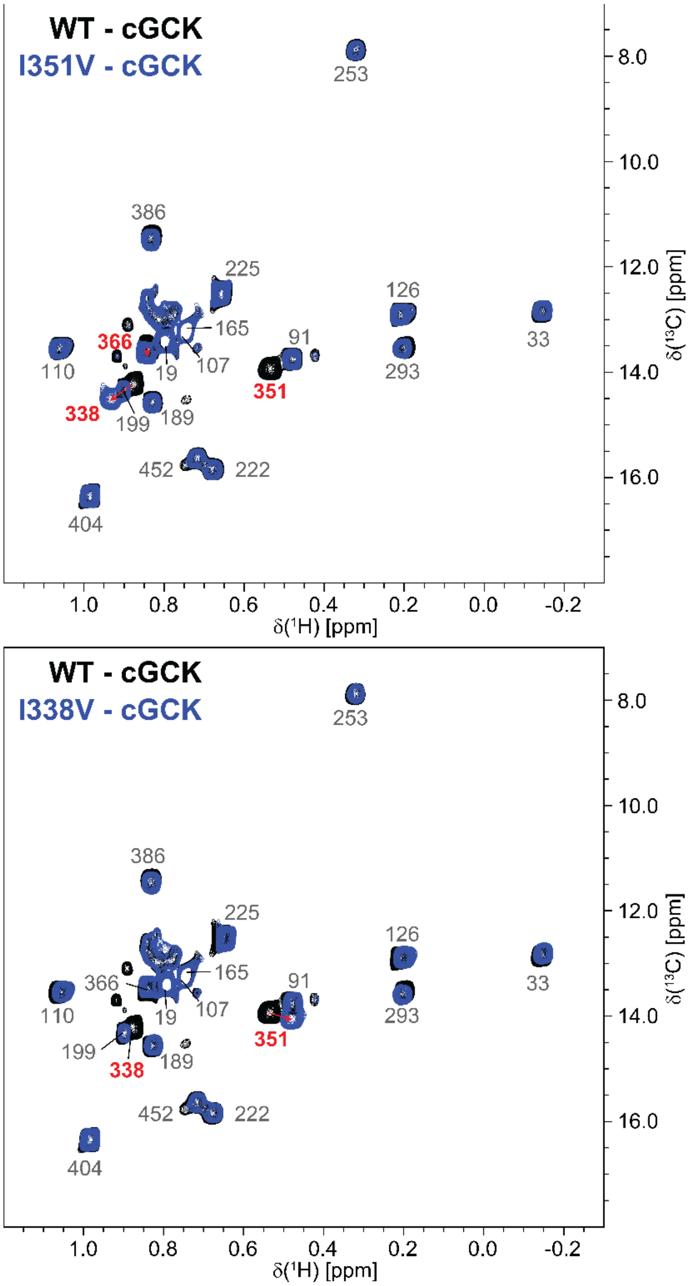

**Fig. S58**. Assignment strategy of isoleucine methyl groups (Case C). (TOP) $^{1}H$-$^{13}C$ ALSOFAST-HMQC spectra of unliganded wild type cGCK and unliganded cGCK-I351V are shown in black and blue, respectively. (BOTTOM) $^{1}H$-$^{13}C$ ALSOFAST-HMQC spectra of unliganded wild type cGCK and unliganded cGCK-I338V are shown in black and blue, respectively. For case C, the isoleucine-to-valine mutation leads to a single peak missing, but other peaks display chemical shift perturbations due to spatial proximity. In the top spectrum, the peak for residue 351 is missing and peaks 338 and 366 move as indicated by red arrows. Likewise, in the bottom spectrum the peak for residue 338 is missing and peak 351 moves as indicated by red arrows. Resonance assignments without changes are shown in gray for reference. Numbering is based on human GCK (GenBank: AAA52562.1; see SI alignment files for corresponding numbering in cGCK and vGCK).

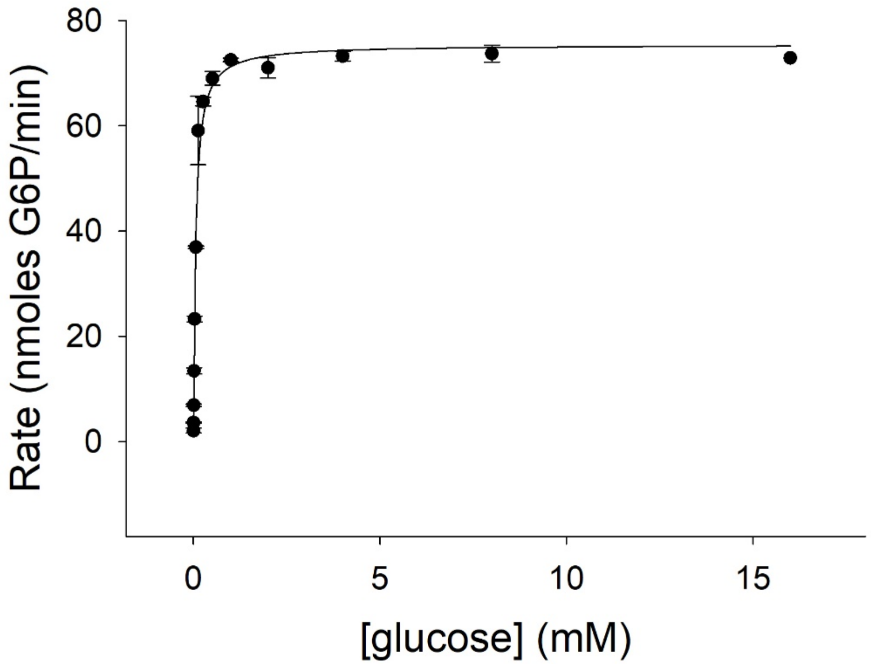

**Fig. S59**. PHHI Substitution GCK glucose kinetics.

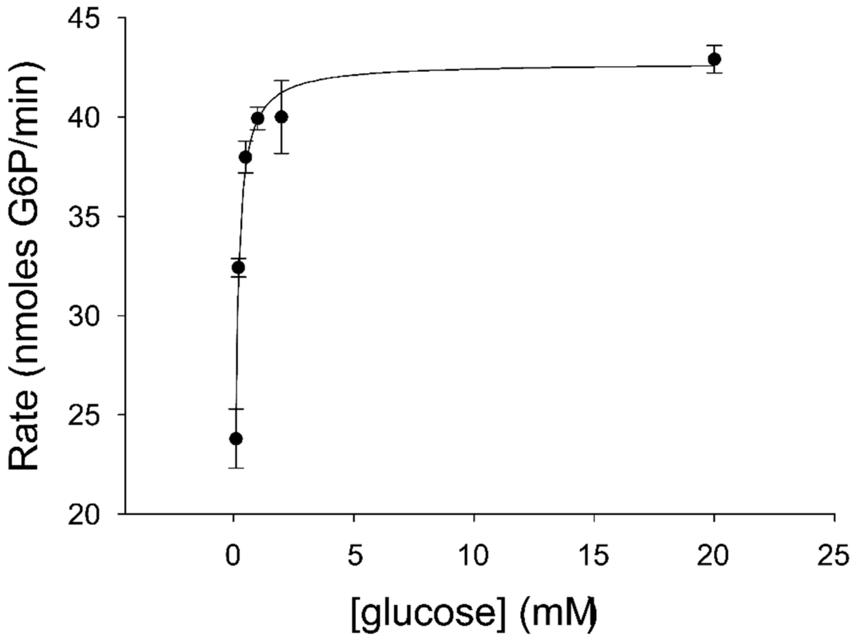

**Fig. S60**. Hybrid GCK glucose kinetics.

Replicate 1

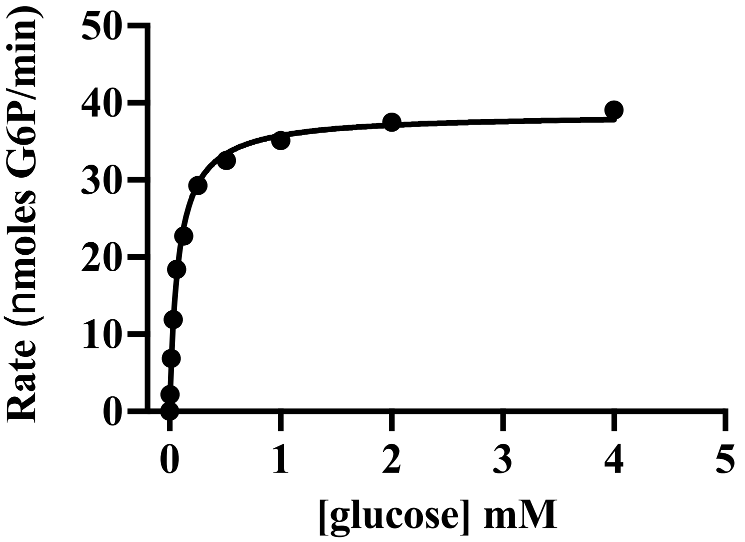

Replicate 2

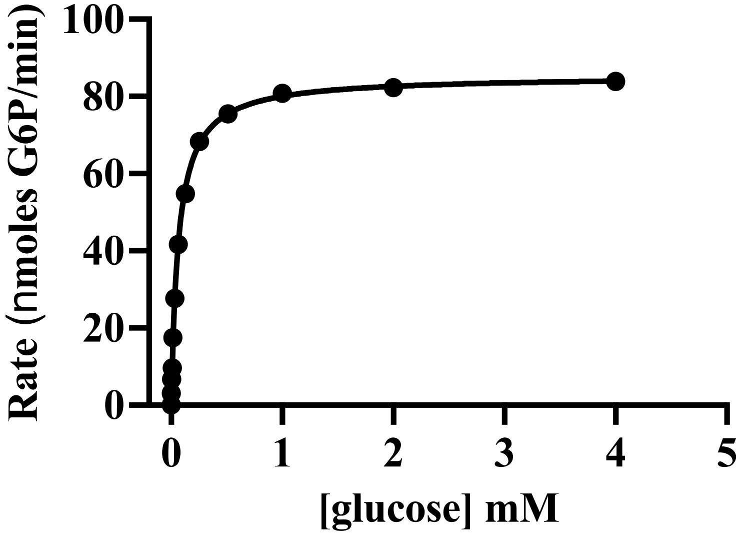

**Fig. S61**. Steady state measurements of the glucokinase reaction rates at varying glucose concentrations for FastML cGCK. Each data point represents the average of triplicate measurements of the highest rate of glucose 6-phosphate production observed over at least one minute of data collection.

Replicate 1

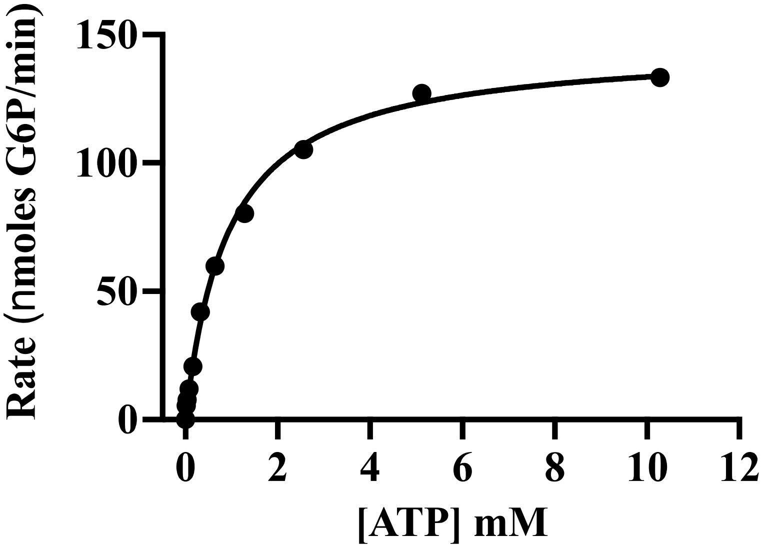

Replicate 2

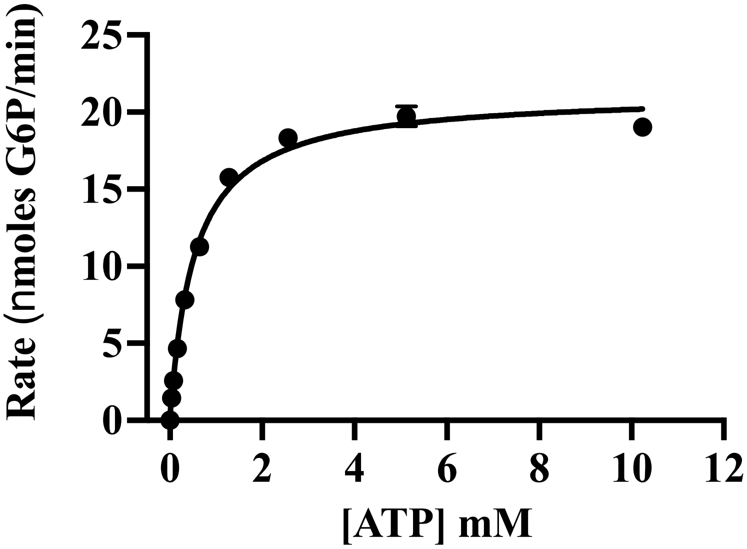

**Fig. S62**. Steady state measurements of the glucokinase reaction rates at varying ATP concentrations for FastML cGCK. Each data point represents the average of triplicate measurements of the highest rate of glucose 6-phosphate production observed over at least one minute of data collection.

Replicate 1

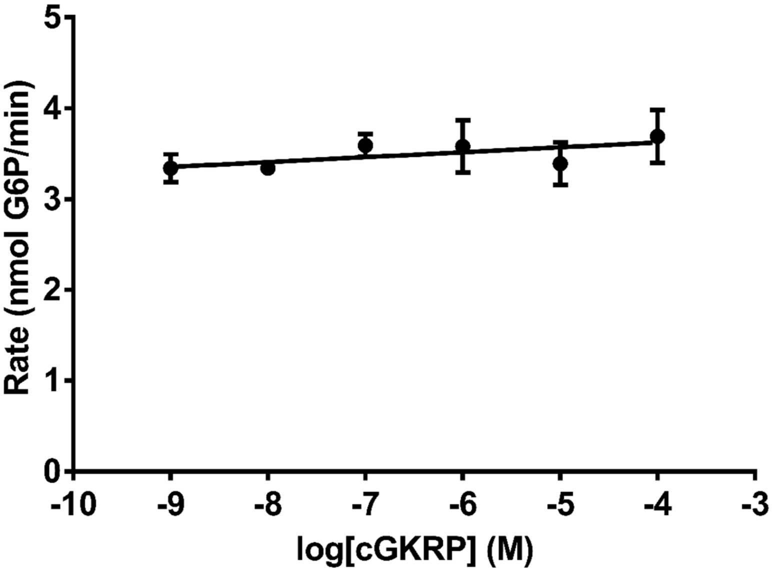

Replicate 2

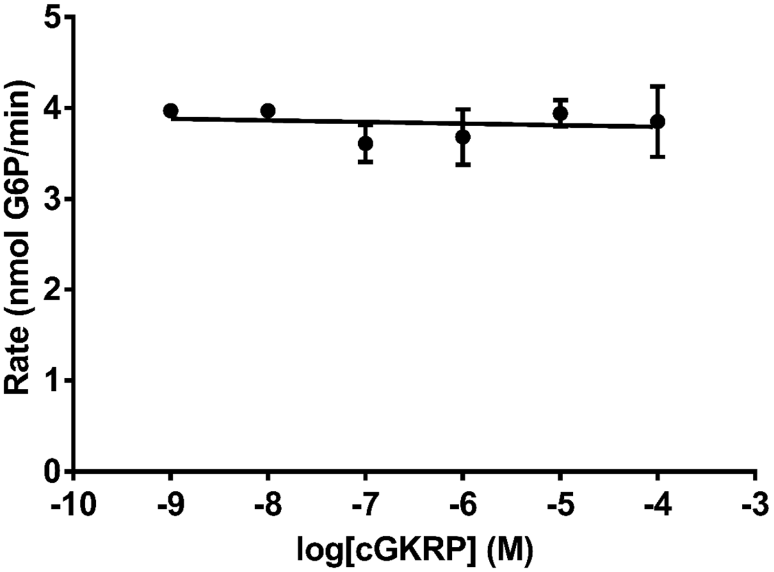

**Fig. S63**. GCK and GKRP protein pairs from the ancestral chordate, represented by cGCK and chordate GKRP (cGKRP), do not form an inhibitory interaction. Each data point represents the average of three technical replicates. Error bars represent standard deviations. Where not visible, the error bars are smaller than the data points.

Replicate 1

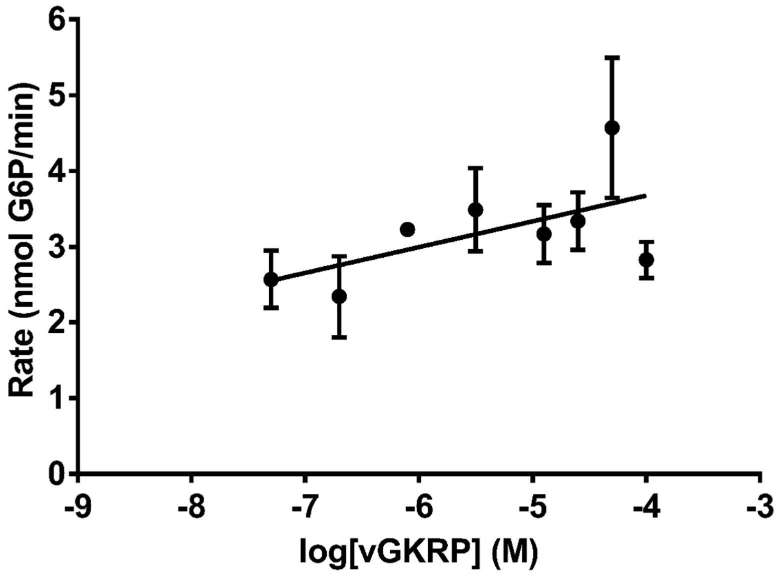

Replicate 2

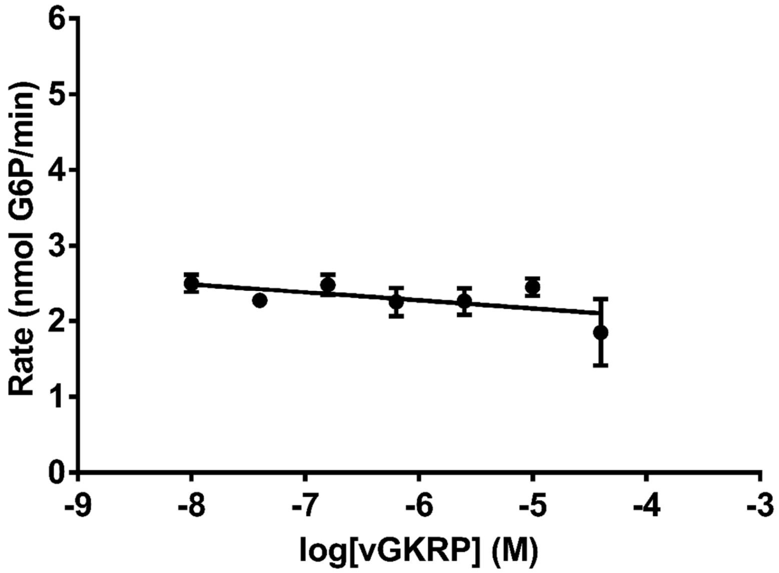

**Fig. S64**. GCK and GKRP protein pairs from the ancestral vertebrate, represented by vGCK and Vertebrate GKRP (vGKRP), do not form an inhibitory interaction. Each data point represents the average of three technical replicates. Error bars represent standard deviations. Where not visible, the error bars are smaller than the data points.

Replicate 1

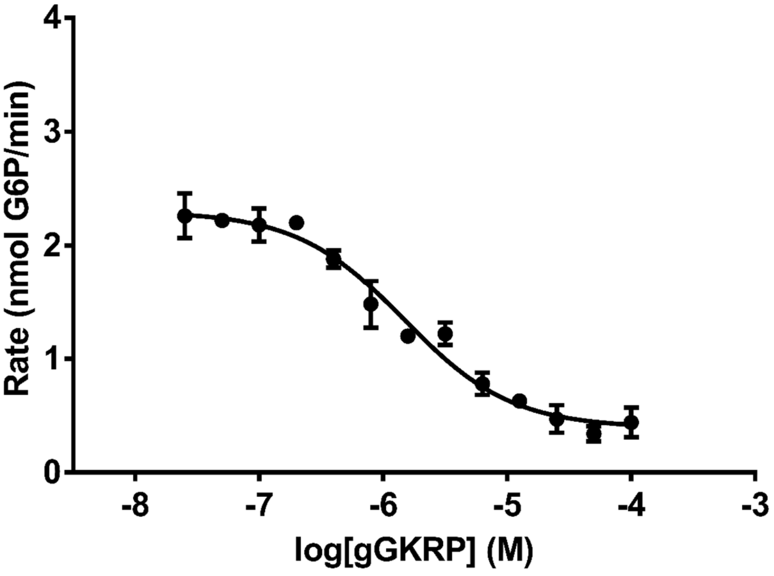

Replicate 2

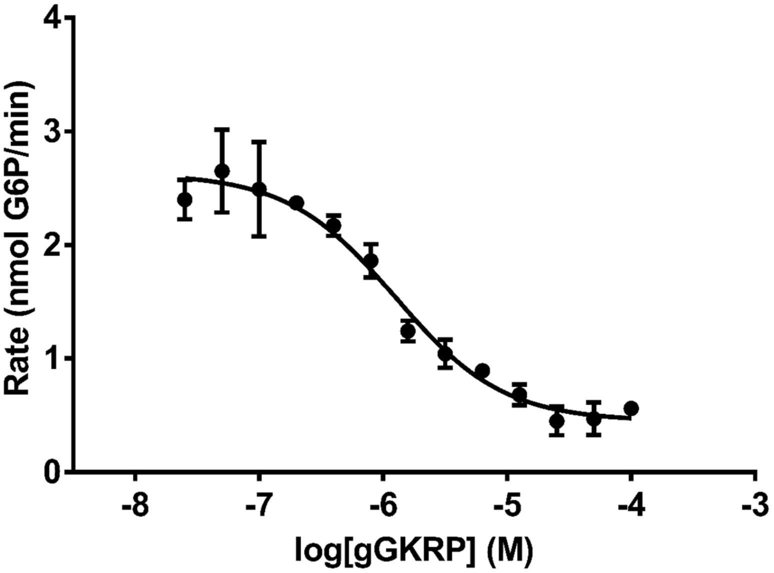

**Fig. S65**. GCK and GKRP protein pairs from the ancestral gnathostome, represented by gGCK and Gnathostome GKRP (gGKRP), form an inhibitory interaction. Each data point represents the average of three technical replicates. Error bars represent standard deviations. Where not visible, the error bars are smaller than the data points.

Replicate 1

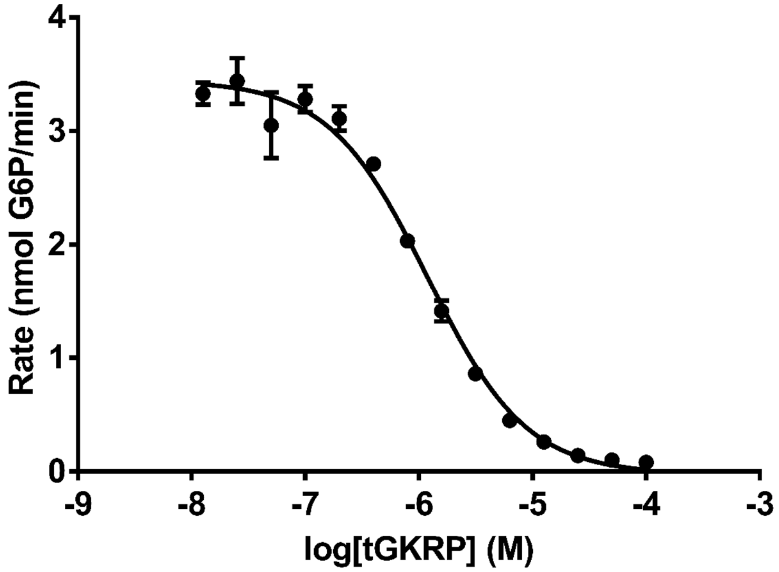

Replicate 2

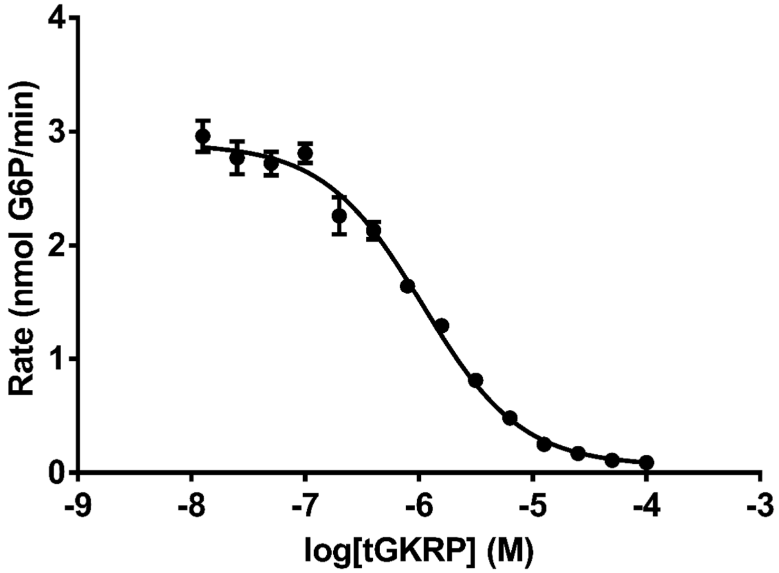

**Fig. S66**. GCK and GKRP protein pairs from the ancestral tetrapod, represented by tGCK and Tetrapod GKRP (tGKRP), form an inhibitory interaction. Each data point represents the average of three technical replicates. Error bars represent standard deviations. Where not visible, the error bars are smaller than the data points.

Replicate 1

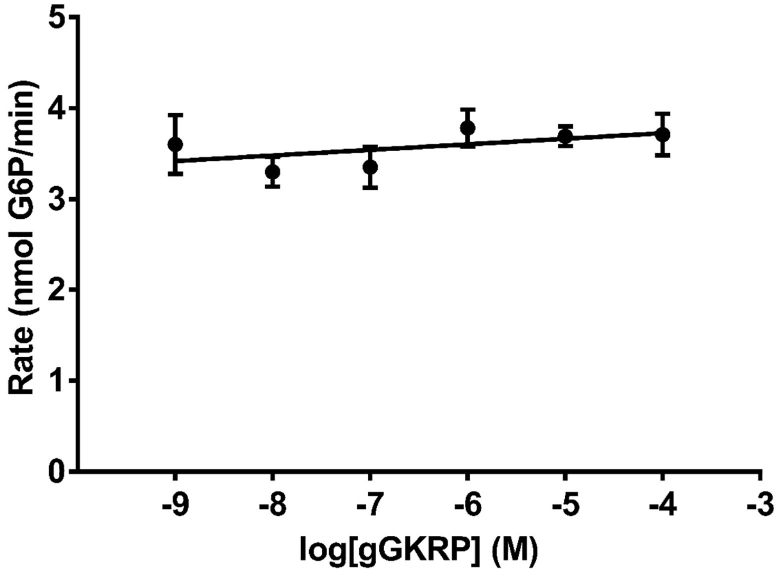

Replicate 2

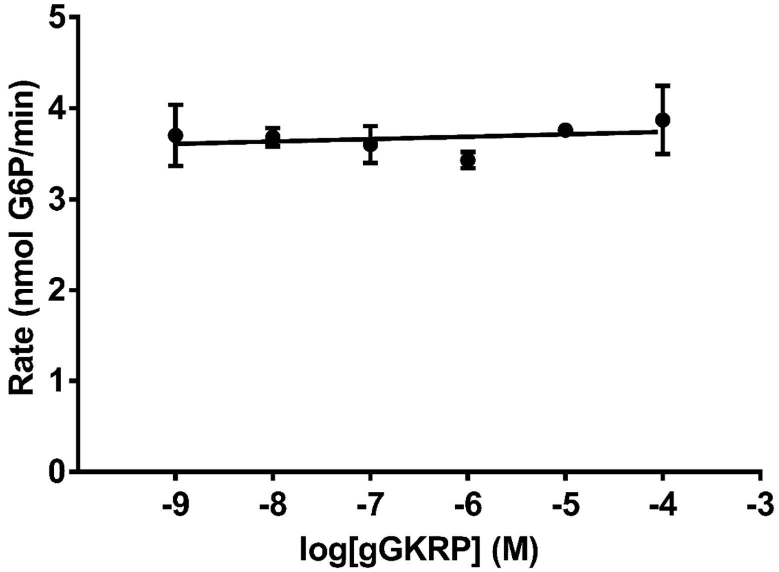

**Fig. S67**. GCK and GKRP protein pairs from the ancestral chordate and the ancestral gnathostome, represented by cGCK and gGKRP, do not form an inhibitory interaction. Each data point represents the average of three technical replicates. Error bars represent standard deviations. Where not visible, the error bars are smaller than the data points.

Replicate 1

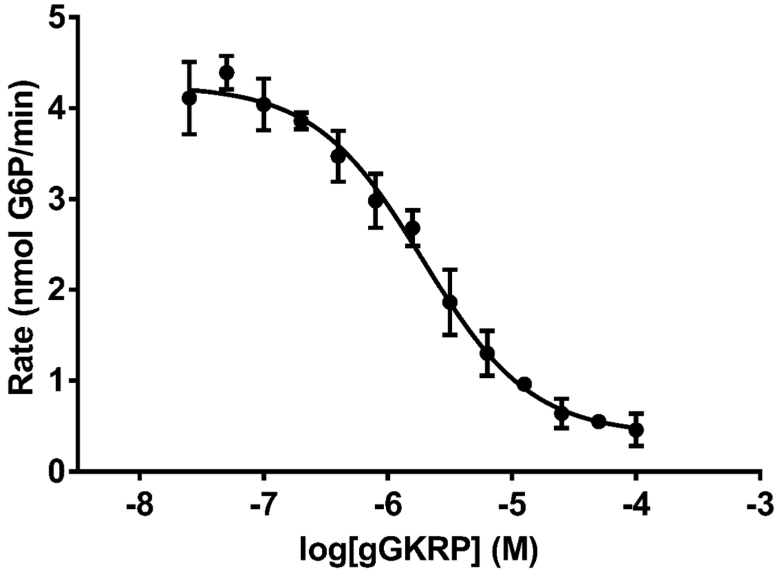

Replicate 2

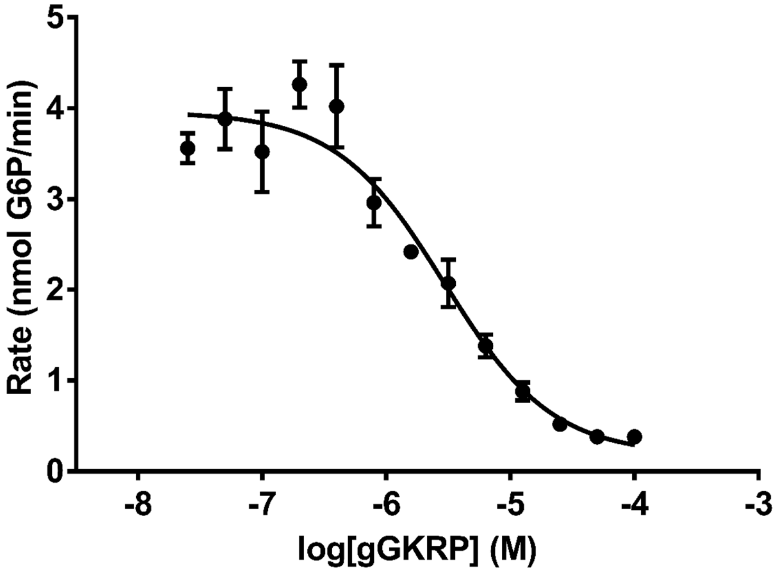

**Fig. S68**. GCK and GKRP protein pairs from the ancestral vertebrate and the ancestral gnathostome, represented by vGCK and gGKRP, form an inhibitory interaction. Each data point represents the average of three technical replicates. Error bars represent standard deviations. Where not visible, the error bars are smaller than the data points.

Replicate 1

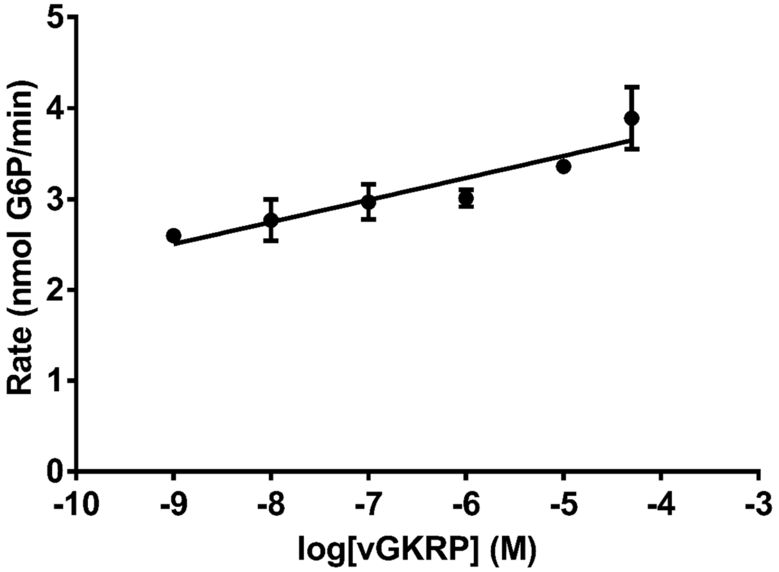

Replicate 2

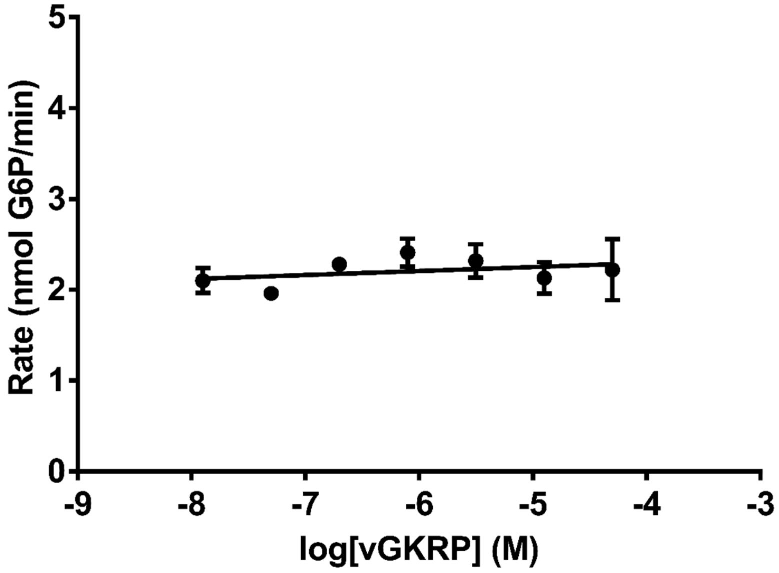

**Fig. S69.** GCK and GKRP protein pairs from the ancestral gnathostome and the ancestral vertebrate, represented by gGCK and vGKRP, do not form an inhibitory interaction. Each data point represents the average of three technical replicates. Error bars represent standard deviations. Where not visible, the error bars are smaller than the data points.

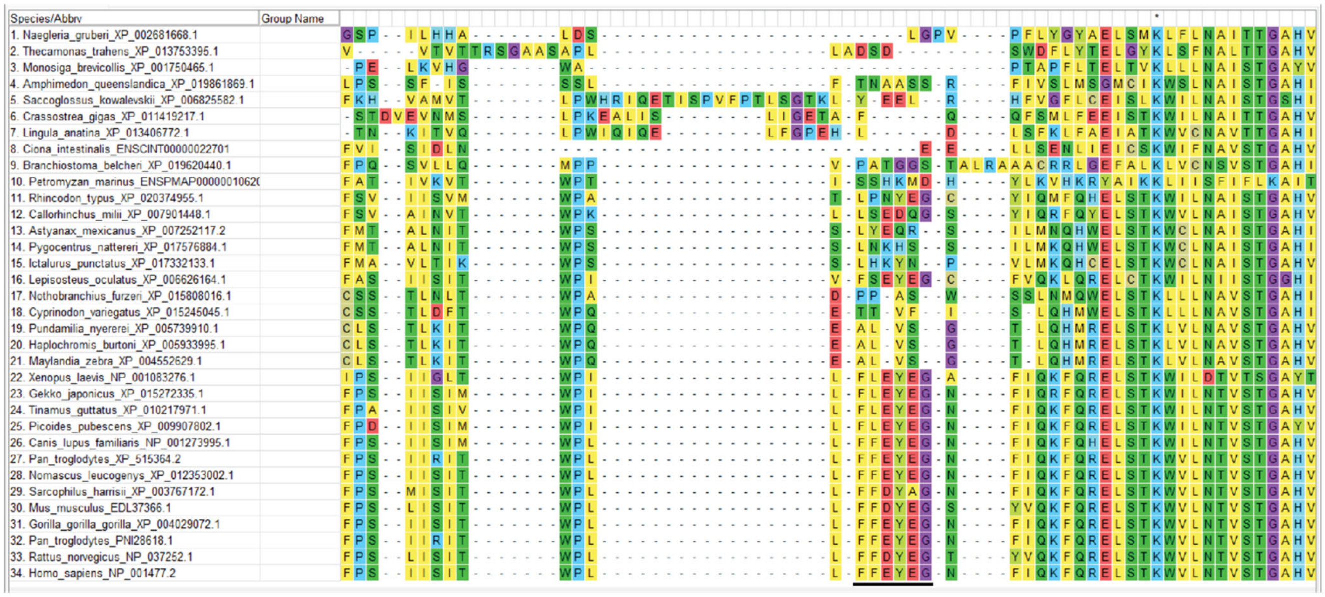

**Fig. S70**. Representative GKRP alignment showing the highly variable loop region that is conserved among jawed vertebrates. The loop in jawed vertebrates is denoted by the black bar.

Replicate 1

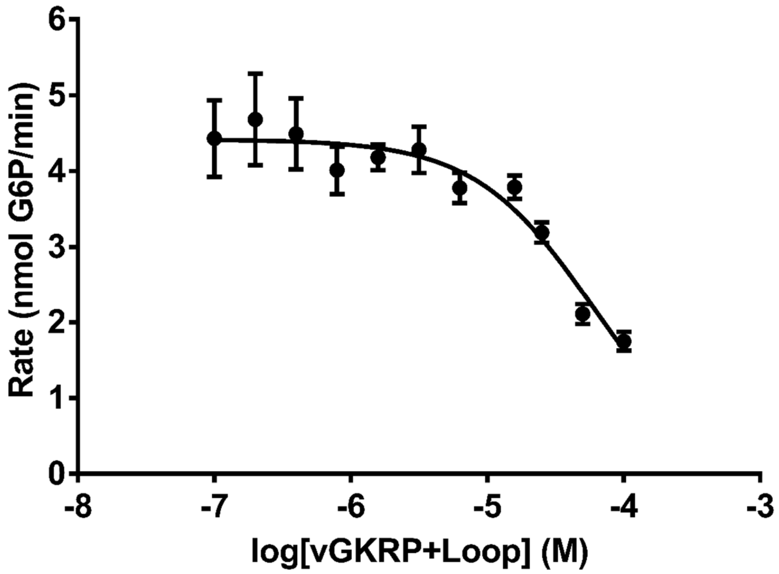

Replicate 2

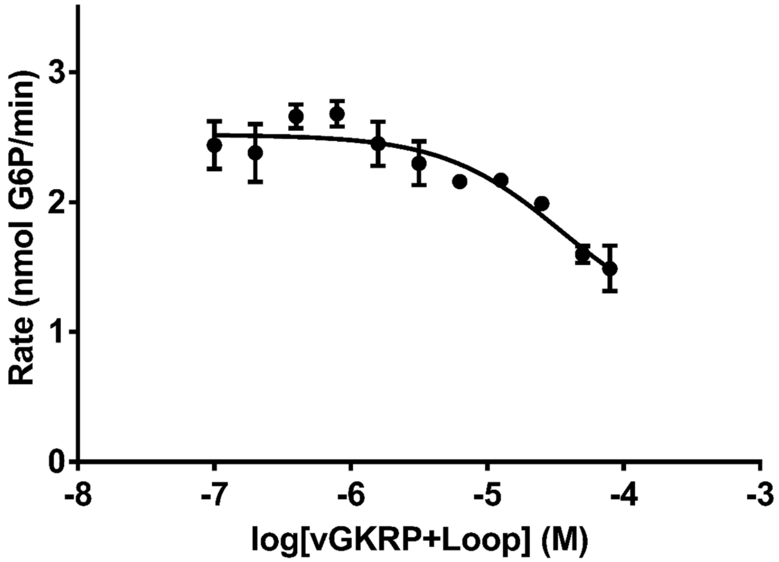

**Fig. S71**. GCK and GKRP protein pairs from the ancestral vertebrate and the ancestral gnathostome, represented by vGCK and the vGKRP+Loop variant, form an inhibitory interaction. Each data point represents the average of three technical replicates. Error bars represent standard deviations. Where not visible, the error bars are smaller than the data points.

| Protein | Full Length Average PP | Expressed Average PP |
|---|---|---|
| cGCK Best | 0.77 | 0.83 |
| vGCK Best | 0.88 | 0.88 |
| gGCK Best | 0.97 | 0.97 |
| cGKRP Best | 0.77 | 0.80 |
| vGKRP Best | 0.79 | 0.83 |
| gGKRP Best | 0.85 | 0.90 |

**Table S1.** Average posterior probabilities for reconstructed and expressed ancestors.

| | cGCK Best | | | | vGCK Best | | | | gGCK Best | |
|---|---|---|---|---|---|---|---|---|---|---|
| | % Identity | # of Diff. | | | % Identity | # of Diff. | | | % Identity | # of Diff. |
| cGCK Worst | 97.60% | 11 | | vGCK Worst | 95.45 | 10 | | gGCK Worst | 96.35 | 6 |
| cGCK AltAll | 87.25 | 58 | | vGCK AltAll | 89.06 | 40 | | | | |
| cGCK FastML | 87.91 | 55 | | | | | | | | |
| cGCK AltAll* | 85.05 | 68 | | | | | | | | |

* Indicates an AltAll version using a cutoff of 0.2 posterior probability of the 2nd best residue as the determination for using the second residue vs using the residue in the best reconstruction

Table S2. Percent identities and number of differences between expressed and characterized ancestors used for determining robustness of measured phenotypes.

| | |
|---|---|
| **Data Collection** | |
| Number of reflections | 66106 (2903 |
| Completeness (%) | 99.1 (88.3 |
| Resolution (Å) | 2.00 (2.03-2.00) |
| Overall <I/Sigma> | 6.8 (0.1) |
| Space group | $P2_12_12_1$ |
| **Cell dimensions** | |
| a | 85.29 Å |
| b | 92.83 Å |
| c | 123.05 Å |
| $\alpha$ | 90.0 ° |
| β | 90.0 ° |
| $\gamma$ | 90.0 ° |
| **Refinement** | |
| Resolution | 2.00 Å |
| R work, R free | 0.229, 0.268 |
| RMS deviations | Bond length (Å) 0.008 and bond angle (°) 1.282 |
| Ramachandran outliers | 0.0 % |
| Ramachandran favored | 97.19 % |
| Rotamer outliers | 0.6 % |
| Clash score | 4.28 |

*Values for the highest resolution shell are shown in parentheses.

**Table S3**. Diffraction and refinement statistics for FastML-cGCK crystals.*

**References**

1. V. Hanson-Smith, A. Johnson, PhyloBot: A Web Portal for Automated Phylogenetics, Ancestral Sequence Reconstruction, and Exploration of Mutational Trajectories. *PLoS Comput Biol* **12**, e1004976 (2016).
2. M. Kaltenbach, J. R. Burke, M. Dindo, A. Pabis, F. S. Munsberg, A. Rabin, S. C. L. Kamerlin, J. P. Noel, D. S. Tawfik, Evolution of chalcone isomerase from a noncatalytic ancestor. *Nat Chem Biol* **14**, 548–555 (2018).
3. G. N. Eick, J. T. Bridgham, D. P. Anderson, M. J. Harms, J. W. Thornton, Robustness of Reconstructed Ancestral Protein Functions to Statistical Uncertainty. *Mol Biol Evol* **34**, 247–261 (2017).
4. H. Ashkenazy, O. Penn, A. Doron-Faigenboim, O. Cohen, G. Cannarozzi, O. Zomer, T. Pupko, FastML: a web server for probabilistic reconstruction of ancestral sequences. *Nucleic Acids Res* **40**, W580–W584 (2012).
5. P. Pal, B. G. Miller, Activating mutations in the human glucokinase gene revealed by genetic selection. *Biochemistry* **48**, 814–816 (2009).
6. M. Larion, B. G. Miller, 23-Residue C-terminal alpha-helix governs kinetic cooperativity in monomeric human glucokinase. *Biochemistry* **48**, 6157–6165 (2009).
7. B. G. Miller, R. T. Raines, Identifying latent enzyme activities: Substrate ambiguity within modern bacterial sugar kinases. *Biochemistry* **43**, 6387–6392 (2004).
8. T. Beck, B. G. Miller, Structural basis for regulation of human glucokinase by glucokinase regulatory protein. *Biochemistry* **52**, 6232–6239 (2013).
9. J. A. Martinez, Q. Xiao, A. Zakarian, B. G. Miller, Antidiabetic Disruptors of the Glucokinase-Glucokinase Regulatory Protein Complex Reorganize a Coulombic Interface. *Biochemistry* **56**, 3150–3157 (2017).
10. A. K. Casey, B. G. Miller, Kinetic Basis of Carbohydrate-Mediated Inhibition of Human Glucokinase by the Glucokinase Regulatory Protein. *Biochemistry* **55**, 2899–2902 (2016).
11. A. C. Whittington, M. Larion, J. M. Bowler, K. M. Ramsey, R. Brüschweiler, B. G. Miller, Dual allosteric activation mechanisms in monomeric human glucokinase. *Proceedings of the National Academy of Sciences* **112**, 11553–11558 (2015).
12. B. H. Gordon, P. Liu, A. C. Whittington, R. Silvers, B. G. Miller, Biochemical methods to map and quantify allosteric motions in human glucokinase. *Methods Enzymol* **685**, 433–459 (2023).
13. Z. Otwinowski, W. Minor, Processing of X-ray diffraction data collected in oscillation mode. *Methods Enzymol* **276**, 307–326 (1997).
14. D. Liebschner, P. V. Afonine, M. L. Baker, G. Bunkoczi, V. B. Chen, T. I. Croll, B. Hintze, L. W. Hung, S. Jain, A. J. McCoy, N. W. Moriarty, R. D. Oeffner, B. K. Poon, M. G. Prisant, R. J. Read, J. S. Richardson, D. C. Richardson, M. D. Sammito, O. V. Sobolev, D. H. Stockwell, T. C. Terwilliger, A. G. Urzhumtsev, L. L. Videau, C. J. Williams, P. D. Adams, Macromolecular structure determination using X-rays, neutrons and electrons: recent developments in Phenix. *Acta Crystallogr D Struct Biol* **75**, 861–877 (2019).
15. M. Larion, R. K. Salinas, L. Bruschweiler-Li, B. G. Miller, R. Brüschweiler, Order-Disorder Transitions Govern Kinetic Cooperativity and Allostery of Monomeric Human Glucokinase. *PLoS Biol* **10**, e1001452 (2012).
16. M. Larion, A. L. Hansen, F. Zhang, L. Bruschweiler-Li, V. Tugarinov, B. G. Miller, R. Brüschweiler, Kinetic Cooperativity in Human Pancreatic Glucokinase Originates from Millisecond Dynamics of the Small Domain. *Angewandte Chemie - International Edition* **54**, 8129–8132 (2015).
